# Lipid membranes trigger misfolding and self-assembly of amyloid β 42 protein into aggregates

**DOI:** 10.1101/587493

**Authors:** Siddhartha Banerjee, Mohtadin Hashemi, Karen Zagorski, Yuri L. Lyubchenko

## Abstract

The assembly of polypeptides and proteins into nanoscale aggregates is a phenomenon observed in a vast majority of proteins. Importantly, aggregation of amyloid β (Aβ) proteins is considered as a major cause for the development of Alzheimer’s disease. The process depends on various conditions and typical test-tube experiments require high protein concentration that complicates the translation of results obtained in vitro to understanding the aggregation process in vivo. Here we demonstrate that Aβ42 monomers at the membrane bilayer are capable of self-assembling into aggregates at physiologically low concentrations, and the membrane in this aggregation process plays a role of a catalyst. We applied all-atom molecular dynamics to demonstrate that the interaction with the membrane surface dramatically changes the conformation of Aβ42 protein. As a result, the misfolded Aβ42 rapidly assembles into dimers, trimers and tetramers, so the on-surface aggregation is the mechanism by which amyloid oligomers are produced and spread.

---

Self-assembly is a widely spread phenomenon in biology and the assembly of proteins into nanoaggerates with various morphologies is a phenomenon of a special attention for decades. Evidence strongly suggests that the development of neurodegenerative diseases, such as Alzheimer’s disease (AD) and Parkinson’s disease (PD), is due to the self-assembly of amyloid β (Aβ) and α-synuclein oligomers, respectively ^1^. These oligomers appear to disrupt the cell homeostasis, by various mechanisms, resulting in the early stages of neurodegenerative diseases ^2–4^ The amyloid cascade hypothesis (ACH) ^5^ is the major model that describes the pathology of AD and other neurodegenerative diseases ^5–9^. However, several orders higher concentrations of Aβ compared with that found *in vivo* are required for the spontaneous aggregation of Aβ, which challenges the validity of ACH. We have discovered that amyloid proteins and peptides are capable of assembling into aggregates at nanomolar concentration range if the process occurs at the surface-liquid interface ^10,11^. In the present study, we monitored the aggregation of Aβ42 at its physiologically relevant low concentration (10 nM) on supported lipid bilayers (SLBs) formed by 1-palmitoyl-2-oleoyl-glycero-3-phosphocholine (POPC) and 1-palmitoyl-2-oleoyl-sn-glycero-3-phospho-L-serine (POPS).

Direct time-lapse AFM imaging revealed Aβ42 aggregation on the bilayer surfaces, while no self-assembly of Aβ42 was detected in bulk solution. We performed computational modeling of the aggregation process on the membrane surfaces to demonstrate that interaction of Aβ42 dramatically changes the conformation of Aβ42 monomers. Moreover, membrane-bound Aβ42 proteins trigger the assembly of dimers, trimers, and tetramers, propagating the misfolded states of the Aβ42 molecules. Thus, interaction with membranes results in the transition of Aβ42 into the aggregation-prone, misfolded conformations. Such conformations have not been reported in simulations or experiments performed in bulk solutions. Given that the membrane-assembled aggregates can dissociate into solution, the discovered on-membrane aggregation may be the mechanism by which amyloid oligomers, or their disease-prone seeds, are produced and spread over the organism.

## Results and discussion

We used time-lapse AFM to directly visualize the aggregation process of Aβ42 protein on supported lipid bilayers formed from 1-palmitoyl-2-oleoyl-glycero-3-phosphocholine (POPC) and 1-palmitoyl-2-oleoyl-sn-glycero-3-phospho-L-serine (POPS) using the approaches described in ^10–13^. A solution of 10 nM Aβ42 was placed on top of the SLBs, assembled on the mica substrate, and imaged at room temperature. SLBs assembled from POPC, POPS, and 1:1 (mol) mixture of POPC:POPS were used. Assembly of SLB with smooth surface topography that remains stable during hours of observation is critical in these studies and we used methodology recently developed in the lab ^11,12^. Typical examples of AFM images of POPC, POPS and POPC:POPS bilayers are shown in Supplementary Fig. 1; cross-section profiles in this figure reveal the height value of 4.2-4.5 nm validating the assembly of the bilayer. The time-dependent aggregation of 10 nM Aβ42 on POPC bilayer is shown in Fig. 1a. Aβ42 aggregates appearing on the SLB after 6 hrs and 9 hrs of incubation are shown as frames (ii) and (iii), respectively. The quantitative analysis of the number and sizes of aggregates shown in Fig. 1b demonstrate that these parameters increase gradually over time, which is similar to our earlier results obtained for Aβ42 aggregation on mica surface ^10^. To characterize the role of electrostatics in the aggregation of Aβ42 on POPC, we performed experiments in the presence of 150 mM NaCl. Supplementary Fig. 2 frames (iii-v) show that ionic strength promotes the aggregation process. Aggregates assemble within 1 hr (Supplementary Fig. 2, frame ii) compared to 5-6 hours in the absence of 150 mM NaCl. The quantitative analysis (Supplementary Fig. 3, a and b) demonstrates that aggregates with volume ~120 nm^3^ appear after 6 hrs in the presence of 150 mM NaCl, while in the absence of NaCl they are formed after 9 hrs.

**Fig. 1.**
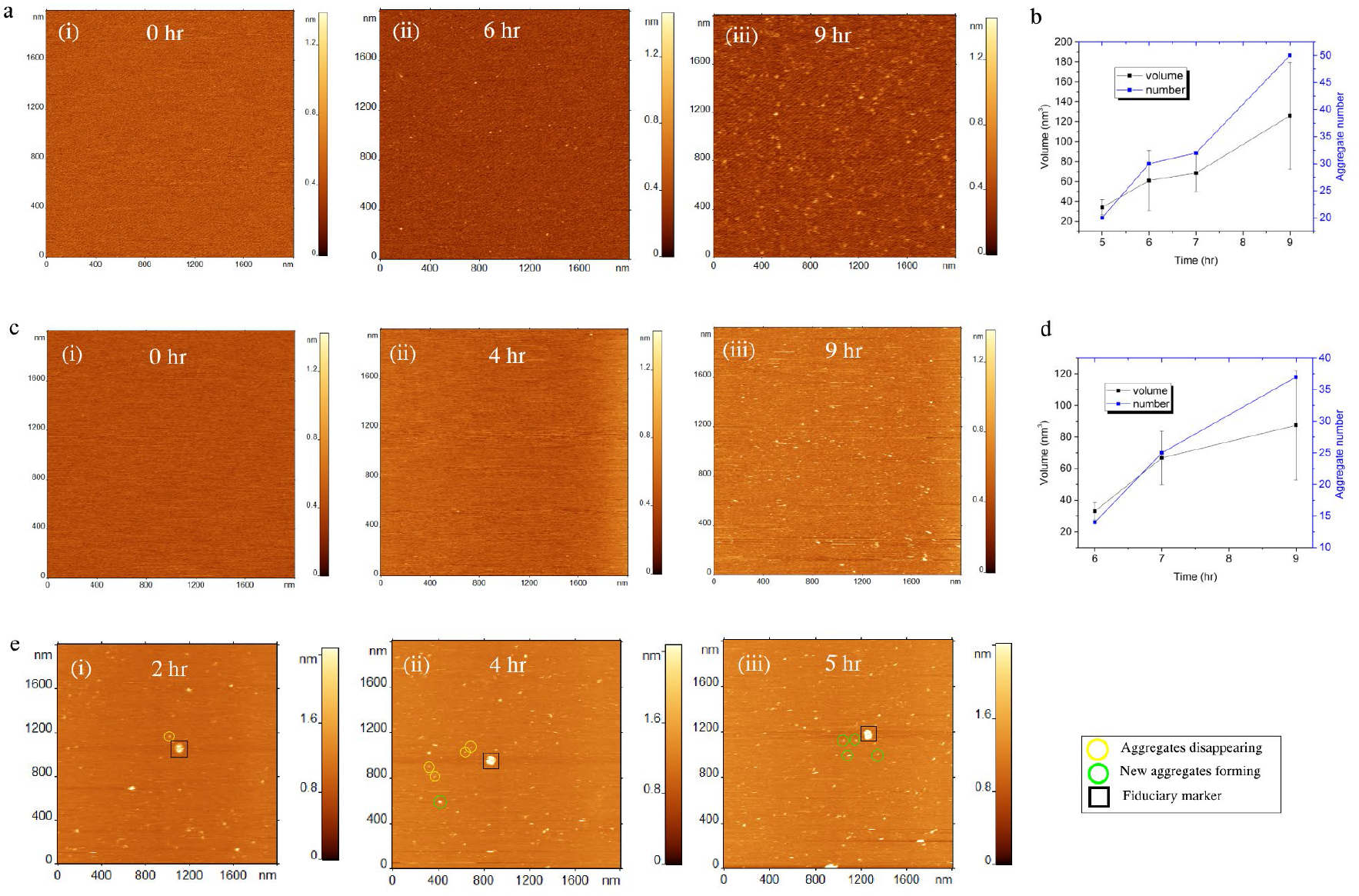
Aggregation of 10 nM Aβ42 on supported lipid bilayers. (**a**) AFM topographic images showing the aggregation on POPC SLB surface. Frames (i)-(iii) show images taken at different time points. More aggregates are visible at later time points. (**b**) The plot shows the increase in number and volume of the aggregates with increase in time. (**c**) Time lapse imaging of Aβ42 aggregation on POPS SLB. (**d**) shows the progress of aggregation as increase in both aggregate number and volume is observed in the plot. (**e**) Appearance and disappearances of aggregates from POPS SLB. Representative examples are shown by circles. Yellow circles indicate that the particular aggregate would leave the surface in the next frame, whereas green circles indicate the appearance of new aggregates which was not present in the previous frame.

Next, we performed similar time-lapse studies for the negatively charged POPS bilayer to test the effect of the bilayer type on the Aβ42 aggregation. Figure 1c, frames (i)-(iii) present typical time-lapse images of the aggregates assembled by 10 nM Aβ42 on POPS SLB. Aggregates start appearing after 4 hrs (frame ii, Fig. 1c) and grow in numbers over time (frame iii, 9 hrs). The time-dependent values of aggregates sizes (volumes of particles) and their numbers are plotted in Fig. 1d and Supplementary Fig. 3c; they demonstrate that both values increase gradually with time. The effect of 150 mM NaCl on the aggregation kinetics of Aβ on POPS SLB was studied as well (Supplementary Fig. 4). Like on POPC, the aggregation process is accelerated in the presence of salt (Supplementary Fig. 3d).

Similar experiments were performed for SLB formed by the equimolar mixture of POPC and POPS. Aggregates appear after 5 hrs of incubation and grow over time (Supplementary Fig. 5a). Ionic strength (150 mM NaCl) accelerates the aggregation (Supplementary Fig. 5b); aggregates formed after 5 hrs of incubation are several times larger compared to those formed in in the absence of NaCl (Supplementary Fig. 5b, frame vi). The cumulative graph illustrating the effect of NaCl for all SLB systems investigated is presented in Supplementary Fig. 6. It demonstrates that NaCl facilitates the formation of aggregates, which appear ~1 hr earlier for all the SLB compositions. Furthermore, aggregates formed on each SLB are larger in presence of 150 mM NaCl.

Time-lapse AFM imaging allowed us to directly monitor the assembly-disassembly process for each individual aggregate (Fig. 1e, frames (ii) and (iii)). The aggregates marked in yellow circles are absent in the next frame and aggregates marked with green circles have appeared. These images demonstrate that assembled Aβ aggregates can dissociate from the surface and go into the solution, so the concentration of aggregates in the solution above the bilayer will increase. To test this conclusion, aliquots from the solution above the bilayer were taken, deposited on mica and imaged. A set of images corresponding to different time points along with the controls (samples without bilayer) demonstrates the accumulation of aggregates in the solution above the bilayer (Supplementary Fig. 7a) with no aggregates in the control (Supplementary Fig. 7b). Quantitatively these data assembled as histograms are shown in Supplementary Fig. 7c.

To determine the underlying mechanism of interaction between Aβ42 and lipid bilayers, we employed extensive MD simulations. POPC and POPS bilayers were assembled using 512 lipid and Aβ42 molecules were added to the bilayers at a distance of 5 nm from the bilayer center. The systems were then simulated for 5 μs each and the interaction of the Aβ molecules with the bilayers was monitored.

Figure 2a shows a few snapshots illustrating the structural change of Aβ monomers induced by its interaction with POPC bilayer. The initial conformation of Aβ monomer (snapshot (i)), taken from our recent paper^14^, has essentially random coiled configuration. However, already after ~ 500 ns, two β-strands appear at the C- and N-termini of the molecule (snapshot (ii)). These segments increase in size followed by the formation of two β-strand regions in the central part of the molecule (snapshot (iii)). The most interesting feature is a cooperative transition within the monomer accompanied by the extension of the β-hairpin structure to the C-terminal region (snapshot (iv)) and this structure remains stable until the end of the 5 μs simulation. The entire conformational transition of the Aβ monomer induced by the interaction with the POPC bilayer is shown graphically in Fig. 2b and Supplementary Fig. 8a and can be viewed in Supplementary Movie 1. Importantly, such dramatic conformational transitions within the Aβ monomer are induced by transient interactions between the POPC surface and peptide residues, with the primary interactions being through residues 7-12 (Supplementary Fig. 8b). A detailed analysis of the peptide-bilayer interaction is given in Supplementary Note 1.

**Fig. 2.**
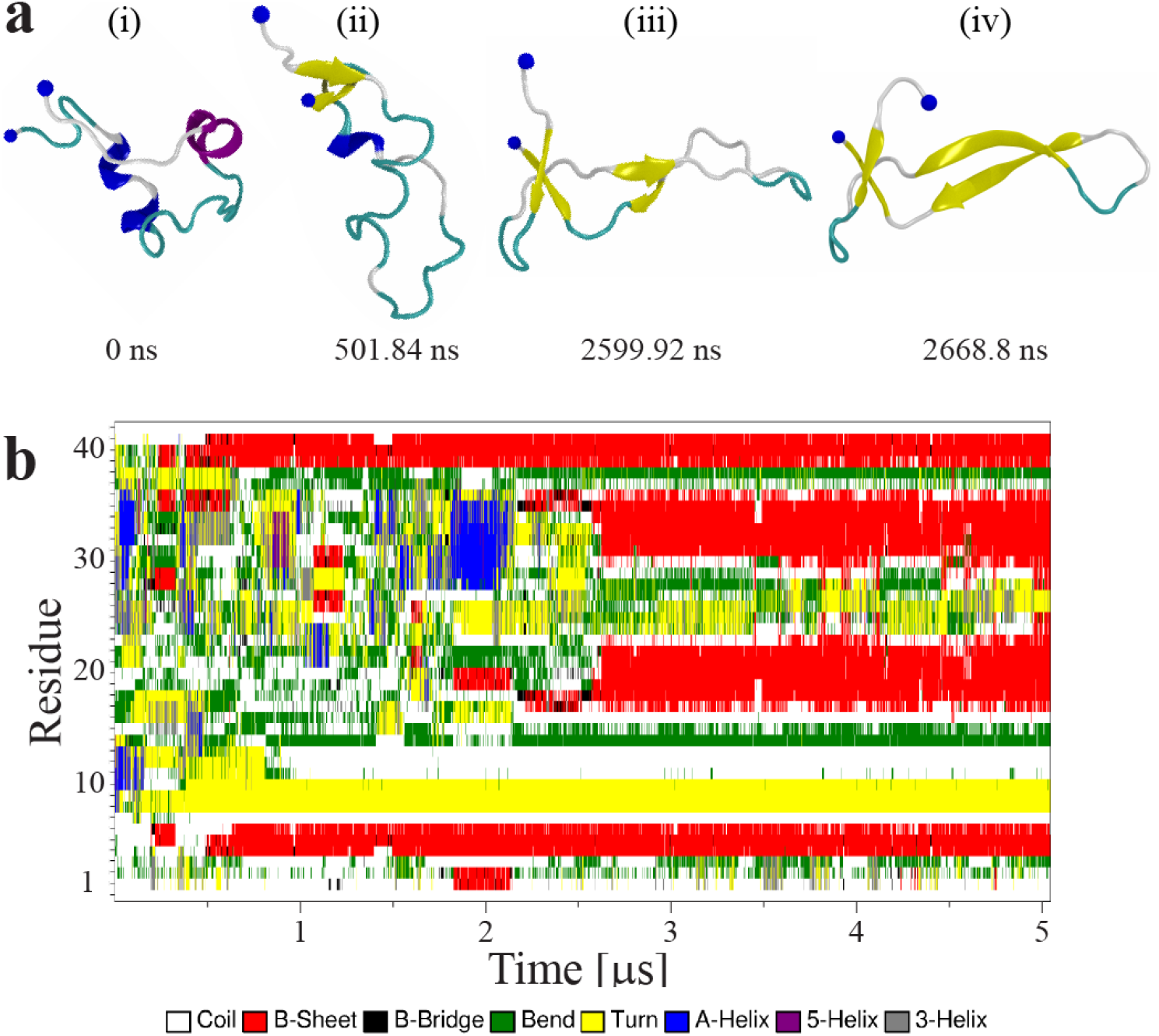
Molecular dynamics simulations of interaction of Aβ42 monomer with POPC bilayer. (**a**) Snapshots showing change of Aβ42 conformation and secondary structure at different timepoints while interacting with POPC bilayer. The monomer is re-arranged throughout the simulation and gradually adopts higher β-structure content. Protein is depicted as cartoon following VMD coloring scheme (yellow β-strands and purple α-helices), N- and C-terminal Cα are presented as a large and a small blue sphere respectively. (**b**) Map of protein secondary structure showing change versus time as determined by DSSP (v3). The N-terminus of the monomer adopts a stable β-strand after 500 ns which remains for the rest of the simulations. Similarly, the central and C-terminal segments experience dramatic changes throughout the simulation and finally settle on three β-strands after ~2.6 μs.

To elucidate the role of conformational transitions within the Aβ monomer in the aggregation process, we modeled the assembly of a dimer by the interaction of an Aβ monomer at the surface with a free monomer. The results in Fig. 3 and Supplementary Movie 2 reveal a number of features of the dimer assembly process. First, in contrast to a slow Aβ42 dimer formation in solution^14^, the dimer on the surface assembles rapidly, after ~23 ns of simulation (Fig. 3a). Second, both monomers within the dimer undergo conformational transitions (Fig. 3b and Supplementary Fig. 9). Third, the conformational transitions are accompanied by the formation of multiple contacts between the two monomers, in which the central part of the surface bound monomer, Mon 1, and the C-terminal segment of the free monomer, Mon 2, play the major role (Fig. 3c and Supplementary Fig. 10). Fourth, similar to the results for the monomer (Fig. 2), transient interactions of the dimer with the bilayer, primarily via Mon 1, are accompanied by conformational changes.

**Fig. 3.**
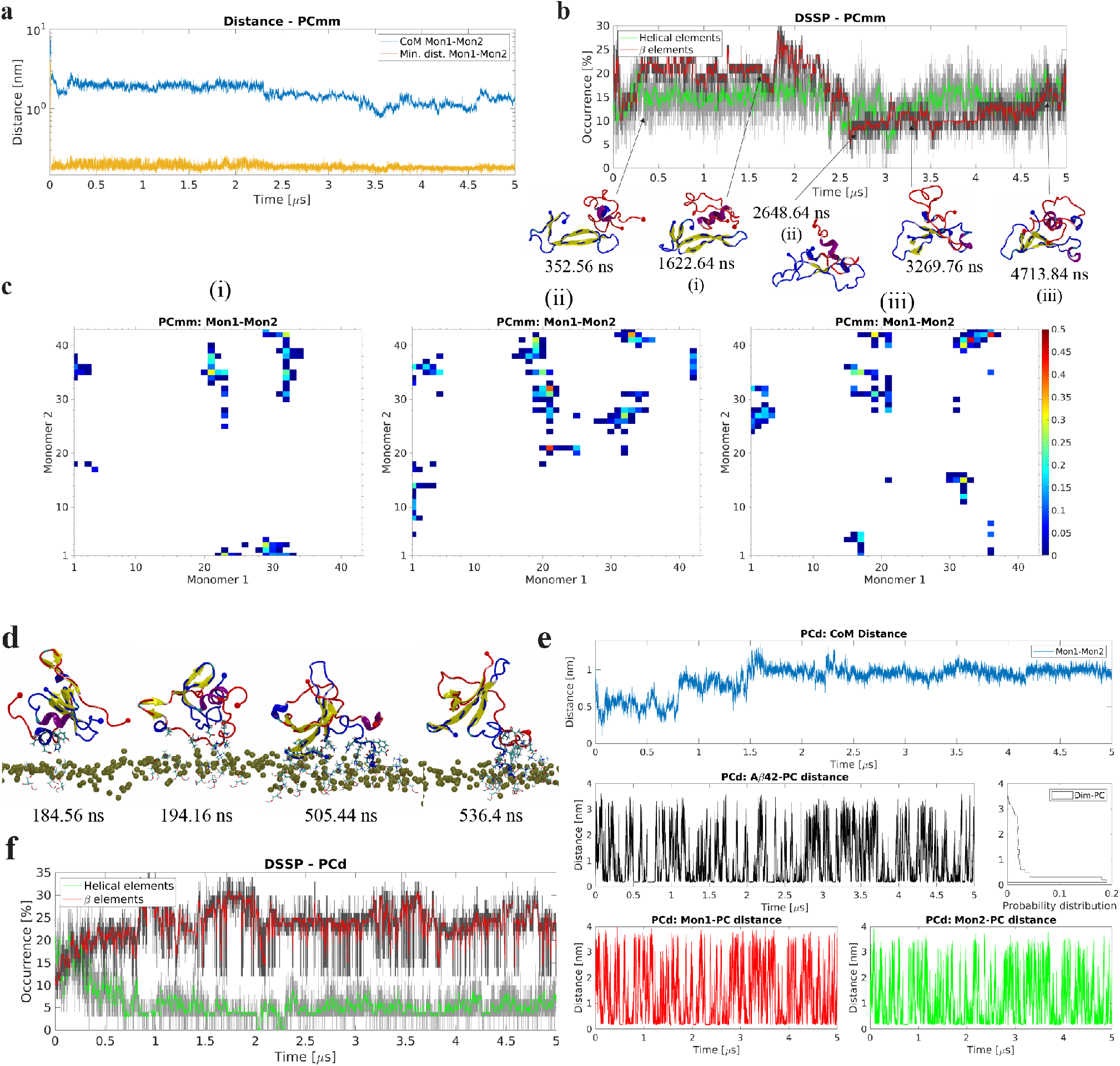
Dynamics of on-surface dimerization of Aβ42 monomers and interaction of preformed dimer with POPC bilayer. (**a**) Time-dependent center of mass (CoM) distance and minimum distance between the two Aβ42 monomers reveal that the dimer rapidly forms and remains stable for the rest of the simulation. (**b**) Evolution of secondary structure β- (sheet and bridge), red, and helical (α-, π-, and 3/10), green, elements, as determined by DSSP, show that the on-surface dimer initially sees an increase in β-content that then deceases and finally recovers and increases to ~20%. The graphs are moving averages using a 1 ns window; raw data is presented as dark and light grey graphs, respectively. Key transition points are highlighted with cartoon representation of the protein structures. Initial surface bound monomer is depicted in blue, with N- and C-terminal Cα as large and small spheres respectively, and the initially free Aβ42 monomer in red. Secondary structure elements are depicted using VMD color scheme (yellow β-strands and purple α-helices). (**c**) Contact maps showing interactions (heavy atom distance <6 Å) between residues of the initially surface-bound monomer, Mon1, and the free monomer, Mon2. Color legend depicts normalized total number of contacts between individual residues. Panel numbers correspond to dimer structures with same numbering in b. (**d**) Snapshots showing change of secondary structure of Aβ42 dimer while interacting with POPC bilayer. Protein side-chains and POPC head groups are shown for interacting residues (distance <6 Å), protein is depicted as cartoon following VMD coloring scheme (yellow β-strands and purple α-helices), N- and C-terminal Cα are represented as large and small spheres in blue and red for Mon1 and Mon2, respectively, POPC P atoms are depicted as gold spheres. (**e**), Evolution of β-(sheet and bridge), red, and helical (α-, π-, and 3/10), green, elements of the protein secondary structure over time show an increase in β-structure content as the simulation progresses, while helical content decreases and plateaus around 5%. The graphs are moving averages using a 1 ns window; raw data is presented as dark and light grey graphs, respectively. (**f**) Top, center of mass distance between the two monomers within the dimer showing that as the dimer interacts with the bilayer it adopts a more expanded conformation. Center, minimum distance between the residues in the dimer and the P atoms of the lipid headgroups versus time showing the transient nature of the dimer-membrane interactions. Bottom, the minimum distance between Aβ42 residues and the P atoms from lipid headgroups showing almost identical plots for the two monomers, suggesting that the dimer does not prefer interactions through a particular monomer.

Previous comprehensive MD simulations of Aβ42 dimer did not reveal extended β-sheets^14^. Here we investigated whether POPC bilayer is capable of changing the conformation of the dimer assembled in solution ^14^. Results shown in Fig. 3d demonstrate dramatic conformational changes within the dimer. Snapshots in Fig. 3d along with conformational analysis in Fig. 3e and Supplementary Fig. 11 demonstrate a rapid formation of β-sheet within the central and C-terminal segments of one monomer (snapshot (i)), followed by the β-strand formation in C-terminal segment in the other monomer (snapshot (ii)). The latter includes a nine-residue strand that remains stable till the end of simulation (snapshot (iii). The dimer remains stable during the entire simulation (Fig. 3f) and is stabilized by contacts in the termini and the central regions (Supplementary Fig. 12). Thus, interaction of Aβ42 monomer and dimer with POPC bilayer dramatically changes the secondary structure of monomers, allowing the formation of extended β-sheets that are also present in amyloid fibrils, but not found in previous simulations of free Aβ42 monomers and dimers in solution. We also simulated assembly of trimers and tetramers on POPC bilayer (Supplementary Figs. 13-21). These data show that oligomers assemble rapidly followed by a dramatic change of the secondary structure of initially free Aβ42 molecules. Detailed structural changes depend on the bilayer type but are generally limited to central segments of Aβ42. Full details and extended technical explanation are presented in Supplementary Note. 2.

Simulations were also performed with POPS bilayers to identify the role of the bilayer composition on the Aβ42 interaction and aggregation. The results shown in Supplementary Figs. 22-37, and their description in Supplementary Note 3, demonstrate that POPS also facilitates β-sheet formation, although Aβ42 molecules contain shorter β-strands compared to conformations formed on POPC bilayer. Additionally, oligomerization is accompanied by conformational transition of the monomers to conformations with greater β-structure (Supplementary Text).

Overall, bilayers dramatically facilitate aggregation of Aβ42 protein allowing the self-assembly to occur at concentration close to physiological ones. Computational modeling revealed a dramatic conformational change in Aβ42 monomer leading to the elevated ability of Aβ42 to assemble into oligomers, suggesting that interaction with membrane surfaces plays a critical role in amyloid aggregation at the physiologically relevant concentrations. Based on these data we hypothesize that self-assembly of the disease-prone Aβ aggregates is driven by the interaction of Aβ with cellular membranes followed by the assembly of dimers first and larger oligomers at later stages. Oligomers can dissociate from the surface into the bulk solution to initiate the oligomer-mediated neurotoxicity of Aβ aggregates. This process is schematically shown in Fig. 4. This model suggests that potential preventions and treatment for AD can be focused on controlling the interaction of amyloids with membranes.

**Figure 4.**
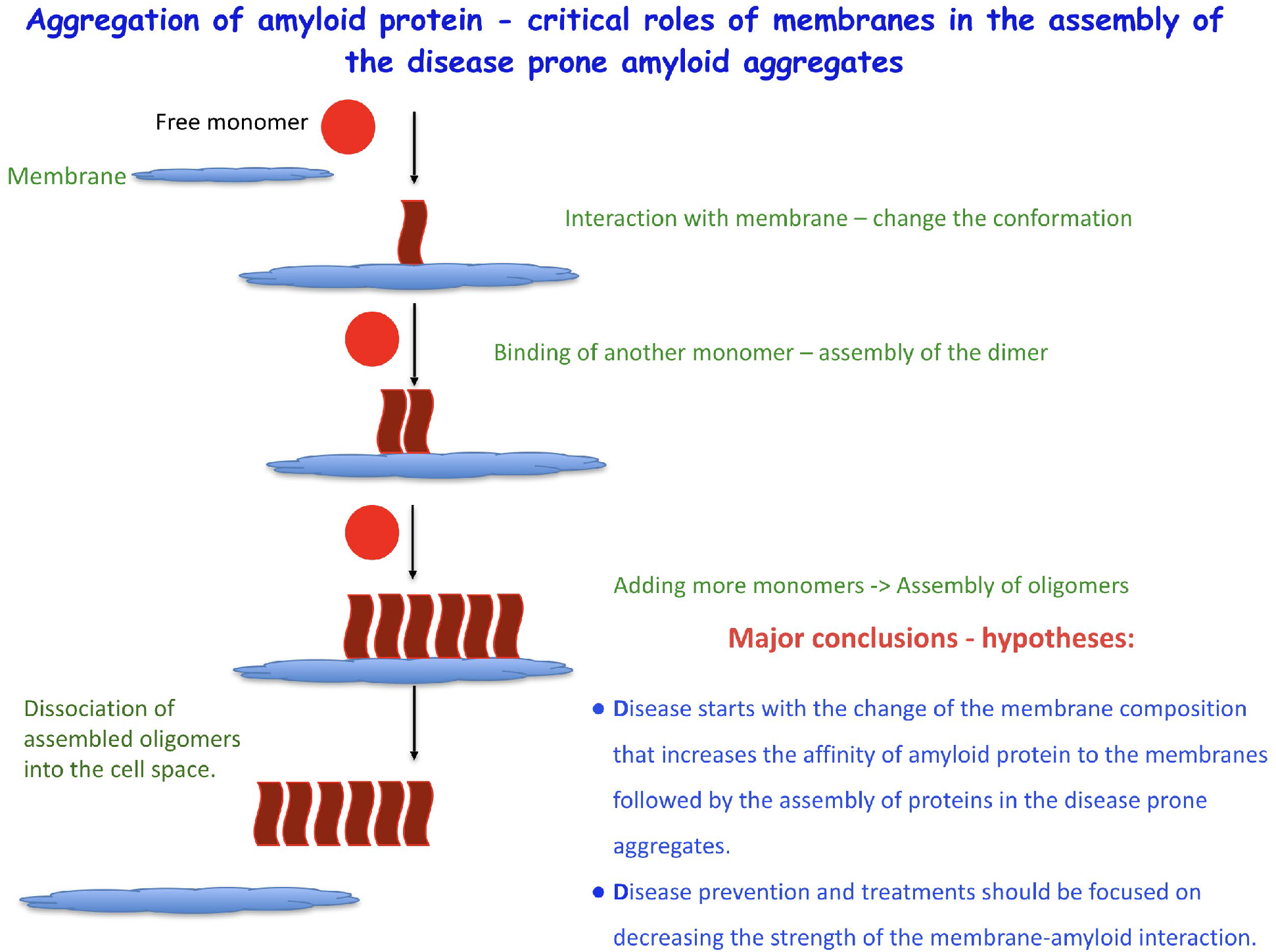
Model of the membrane-mediated aggregation of Aβ protein.

## Methods

### Materials

Amyloid beta-42 (Aβ42) was purchased from AnaSpec, Fremont, CA. 1-Palmitoyl-2-oleoyl-sn-glycero-3-phosphocholine (POPC) and 1-palmitoyl-2-oleoyl-sn-glycero-3-phospho-L-serine (POPS) were obtained from Avanti Polar Lipids, Inc, Alabama, US; Chloroform (Sigma Aldrich Inc.); sonicator (Branson 1210, Branson Ultrasonics, Danbury, CT). Mainly two buffer solutions were used. 10 mM sodium phosphate buffer, pH 7.4 (for without salt condition), 10 mM sodium phosphate, 150 mM NaCl, pH 7.4 (for with salt condition). Deionized water (18.2 MΩ, filter pore size: 0.22 μm; APS Water Services Corp., Van Nuys, CA) was used in all the experiments wherever required. Glass vial and glass pipettes (Fisher Scientific, Waltham, Massachusetts, USA) were used to handle the lipid solution.

### Preparation of Aβ42 solution

Protein solution was prepared as described previously ^10^. Briefly, measured amount of Aβ42 was dissolved and sonicated for 5 min in 100 μl of 1,1,1,3,3,3-hexafluoroisopropanol (HFIP) to remove any preformed oligomers. The tubes were then put into vacufuge until the solvent is completely evaporated. The stock solution was prepared in DMSO and kept at −20 °C. 10 nM solution was prepared from the stock in 10 mM sodium phosphate, pH 7.4 just before the experiment.

### Preparation of supported lipid bilayers (SLBs)

The similar methodology was obtained as described previously ^15^. Briefly, a 25 mg/ml stock solution of POPC and POPS was prepared in chloroform. Glass vial and glass pipettes were used to handle the lipid solution. Stock solution was stored in −20 °C. An aliquot of 20 μl was taken in another glass vial and dried with the flow of Ar and kept overnight in a vacuum chamber to remove any trace of chloroform. Then the dried lipid is resuspended in 1 ml 10 mM sodium phosphate, pH 7.4 buffer solution to prepare a 0.5 mg/ml solution which was used for SLB preparation. The solution is gently vortexed for 1 min and then sonicated until the solution became clearer. This solution was then deposited onto a freshly cleaved mica surface, which was attached to a glass slide and incubated for 1 hr at 60 °C. After the incubation, the slide was cooled to the room temperature, the excess of the lipid solution was removed, and the substrate was rinsed with the buffer gently. The prepared SLB was never allowed to dry by keeping ~300 μl buffer on top of it.

### AFM imaging

Time-lapse AFM imaging was performed in MFP-3D (Asylum Research, Santa Barbara, CA) instrument in tapping mode. MSNL AFM probe (cantilever ‘E’) was used for all the experiments. The nominal resonance frequency and the spring constant of the probe were 7-9 kHz and 0.1 N/m, respectively. Typical scan speed was kept at 1-2 Hz. For time-lapse imaging, images of the same area of the surface were acquired at different time points. Typically, continuous scanning was avoided. Between each time-point the cantilever was electronically retracted by the software and engaged again to record the image for the next time point.

### Analysis of AFM data

All the AFM images shown were subjected to minimum processing. Only flattening (1^st^ order polynomial) was performed by Femtoscan software (Advanced Technologies Center, Moscow, Russia). The volume of the aggregates was measured by the ‘Grain analysis’ tool of the software and then the distribution of the volume was obtained by plotting them in histogram and fitting the histogram with Gaussian distribution using Origin Pro software (OriginLab, Northampton, MA). Standard deviation was obtained as half-width of the distribution.

### Computational methods

#### Bilayer assembly

To generate the initial bilayers of POPC and POPS used for all simulation systems, we employed CHARMM-GUI ^16^ to produce bilayers consisting of 512 lipid molecules with 40 TIP3P waters ^17^ per lipid. Lipid systems were neutralized and kept at 150 mM ionic concentration using Na and Cl counterions and converted to AMBER format using the lipid17 force field (an extension and refinement of lipid14 ^18^). After which 150 ns NPT MD simulations were performed using a 2 fs integration time step. The simulations employed periodic boundary conditions with a semi-isotropic pressure coupling at 1 bar, a constant temperature of 300 K, non-bonded interactions truncated at 10 Å, and electrostatic interactions treated using particle-mesh Ewald ^19^. Simulations were performed using the Amber16 package ^20^.

#### Interaction of Aβ42 with bilayers

To investigate the interaction of Aβ42 monomer with the bilayers, we extracted the POPC and POPS bilayers from the final frame of the pure bilayer simulations, added Aβ42 molecules (monomer or dimer conformations taken from ref. ^14^) at 5 nm center-of-mass (CoM) from the bilayer center, solvated in TIP3P water (in an orthorhombic box [a=b≠c] with side:height ratio 0.75), neutralized with NaCl counter ions, and maintained a final NaCl concentration of 150 mM. Proteins were described using the Amber ff99SB-ILDN force field ^21^. Each system then underwent H-mass repartitioning to increase H mass to 3.024 Da allowing for 4 fs time steps ^22^, following which the systems were simulated as an NPT ensemble for 50 ns (using the same parameters as previous bilayer simulations) before being submitted to the special purpose supercomputer Anton2 for long production runs. Simulations on Anton2 employed the multigrator algorithm and treated electrostatics using the Gaussian split Ewald method.

#### Interaction between membrane-bound and free Aβ42

To investigate the interaction between membrane-bound and free Aβ42 species, we used the last membrane-bound conformation of the previous simulation systems and added a monomer or a dimer to the simulation systems following the same procedure as used initially to add Aβ42 molecules to the bilayer systems. Newly added molecules were placed 4 nm CoM with respect to the membrane-bound molecules and at 5 nM distance to the membrane core. Simulation parameters and steps were the same as the initial Aβ42-bilayer simulations.

**Table S1.** Description and nomenclature of simulated bilayer systems.

| **Table S1. Description and nomenclature of simulated bilayer systems.** |  |  |  |
| --- | --- | --- | --- |
| **Name** | **Bilayer** | **Aβ42 species** | **Duration [μs]** |
| PC | POPC | - | 0.150 |
| PS | POPS | - | 0.150 |
| PCm | POPC | Monomer | 5 |
| PSm | POPS | Monomer | 5 |
| PCmm | POPC | Bound monomer + monomer | 5 |
| PCmd | POPC | Bound monomer + dimer | 5 |
| PSmm | POPS | Bound monomer + monomer | 5 |
| PSmd | POPS | Bound monomer + dimer | 5 |
| PCd | POPC | Dimer | 5 |
| PSd | POPS | Dimer | 5 |
| PCdm | POPC | Bound dimer + monomer | 5 |
| PCdd | POPC | Bound dimer + dimer | 5 |
| PSdm | POPS | Bound dimer + monomer | 5 |
| PSdd | POPS | Bound dimer + dimer | 5 |

#### Analysis of MD trajectories

Gromacs suite of programs (v2016) ^23^ was used to analyze the obtained simulation trajectories. In addition, the Python-based TimeScapes package ^24^ was used to identify residues important for the stability of the Aβ oligomers. The analysis is based on pivot residue pseudodihedral angles defined by four consecutive Cα.

## Acknowledgments

The work at the University of Nebraska Medical Center (UNMC) was supported by grants from the National Institutes of Health to Y.L.L. (GM096039, GM118006). Anton 2 computer time was provided by the Pittsburgh Supercomputing Center (PSC) through Grant R01GM116961 from the National Institutes of Health. The Anton 2 machine at PSC was generously made available by D.E. Shaw Research.

## Author contributions

Y.L.L., S.B. and M.H. designed the project. S.B and K.Z. performed the AFM experiments; M.H. performed the molecular dynamics simulations. All authors wrote and edited the manuscript.

## Supplementary Information

Supplementary Notes 1-3

Figures S1-S37

Movies S1-S2

## References

1 Chiti, F. & Dobson, C. M. Protein Misfolding, Amyloid Formation, and Human Disease: A Summary of Progress Over the Last Decade. Annual review of biochemistry 86, 27–68, doi:10.1146/annurev-biochem-061516-045115 (2017).

2 Venda, L. L., Cragg, S. J., Buchman, V. L. & Wade-Martins, R. alpha-Synuclein and dopamine at the crossroads of Parkinson’s disease. Trends Neurosci 33, 559–568, doi:10.1016/j.tins.2010.09.004 (2010).

3 Bate, C. & Williams, A. Monomeric amyloid-β reduced amyloid-β oligomer-induced synapse damage in neuronal cultures. Neurobiology of disease 111, 48–58, doi:https://doi.org/10.1016/j.nbd.2017.12.007 (2018).

4 Zhao, Y. et al. TREM2 Is a Receptor for beta-Amyloid that Mediates Microglial Function. Neuron 97, 1023–1031.e1027, doi:10.1016/j.neuron.2018.01.031 (2018).

5 Hardy, J. A. & Higgins, G. A. Alzheimer’s disease: the amyloid cascade hypothesis. Science 256, 184–185 (1992).

6 Mohamed, T., Shakeri, A. & Rao, P. P. N. Amyloid cascade in Alzheimer’s disease: Recent advances in medicinal chemistry. European Journal of Medicinal Chemistry 113, 258–272, doi:https://doi.org/10.1016/j.ejmech.2016.02.049 (2016).

7 A. Armstrong, R. Review paper<br>A critical analysis of the ‘amyloid cascade hypothesis’. Folia Neuropathologica 52, 211–225, doi:10.5114/fn.2014.45562 (2014).

8 Hardy, J. Has the amyloid cascade hypothesis for Alzheimer’s disease been proved? Current Alzheimer research 3, 71–73 (2006).

9 Hardy, J. Alzheimer’s disease: the amyloid cascade hypothesis: an update and reappraisal. Journal of Alzheimer’s disease: JAD 9, 151–153 (2006).

10 Banerjee, S. et al. A novel pathway for amyloids self-assembly in aggregates at nanomolar concentration mediated by the interaction with surfaces. Sci Rep 7, 45592, doi:10.1038/srep45592 (2017).

11 Lv, Z., Banerjee, S., Zagorski, K. & Lyubchenko, Y. L. Supported Lipid Bilayers for Atomic Force Microscopy Studies. Methods Mol Biol 1814, 129–143, doi:10.1007/978-1-4939-8591-3_8 (2018).

12 Lv, Z. et al. Phospholipid membranes promote the early stage assembly of asynuclein aggregates. bioRxiv preprint, doi:https://doi.org/10.1101/295782 (2018).

13 Lyubchenko, Y. L. Direct AFM visualization of the nanoscale dynamics of biomolecular complexes. Journal of Physics D: Applied Physics 51, 403001 (2018).

14 Zhang, Y., Hashemi, M., Lv, Z. & Lyubchenko, Y. L. Self-assembly of the full-length amyloid Abeta42 protein in dimers. Nanoscale 8, 18928–18937, doi:10.1039/c6nr06850b (2016).

15 Lv, Z. et al. Phospholipid membranes promote the early stage assembly of α-synuclein aggregates. bioRxiv (2018).

16 Jo, S., Kim, T., Iyer, V. G. & Im, W. CHARMM-GUI: a web-based graphical user interface for CHARMM. J Comput Chem 29, 1859–1865, doi:10.1002/jcc.20945 (2008).

17 Jorgensen, W. L., Chandrasekhar, J., Madura, J. D., Impey, R. W. & Klein, M. L. Comparison of simple potential functions for simulating liquid water. J. Chem. Phys. 79, 926–935, doi:doi:http://dx.doi.org/10.1063/1.445869 (1983).

18 Dickson, C. J. et al. Lipid14: The Amber Lipid Force Field. J Chem Theory Comput 10, 865–879, doi:10.1021/ct4010307 (2014).

19 Darden, T., York, D. & Pedersen, L. Particle mesh Ewald: An N·log(N) method for Ewald sums in large systems. The Journal of Chemical Physics 98, 10089–10092, doi:10.1063/1.464397 (1993).

20 D.A. Case, V. B., J.T. Berryman, R.M. Betz, Q. Cai, D.S. Cerutti, T.E. Cheatham, III, T.A. Darden, R.E. Duke, H. Gohlke, A.W. Goetz, S. Gusarov, N. Homeyer, P. Janowski, J. Kaus, I. Kolossváry, A. Kovalenko, T.S. Lee, S. LeGrand, T. Luchko, R. Luo, B. Madej, K.M. Merz, F. Paesani, D.R. Roe, A. Roitberg, C. Sagui, R. Salomon-Ferrer, G. Seabra, C.L. Simmerling, W. Smith, J. Swails, R.C. Walker, J. Wang, R.M. Wolf, X. Wu and P.A. Kollman. (University of California, San Francisco, 2016).

21 Lindorff-Larsen, K. et al. Improved side-chain torsion potentials for the Amber ff99SB protein force field. Proteins 78, 1950–1958, doi:10.1002/prot.22711 (2010).

22 Hopkins, C. W., Le Grand, S., Walker, R. C. & Roitberg, A. E. Long-Time-Step Molecular Dynamics through Hydrogen Mass Repartitioning. J Chem Theory Comput 11, 1864–1874, doi:10.1021/ct5010406 (2015).

23 Abraham, M. J. et al. GROMACS: High performance molecular simulations through multi-level parallelism from laptops to supercomputers. SoftwareX 1–2, 19–25, doi:10.1016/j.softx.2015.06.001 (2015).

24 Wriggers, W. et al. Automated Event Detection and Activity Monitoring in Long Molecular Dynamics Simulations. J Chem Theory Comput 5, 2595–2605, doi:10.1021/ct900229u (2009).

