## Supplementary Information for "Lipid membranes trigger misfolding and self-assembly of amyloid β 42 protein into aggregates"

**This PDF file includes:**

Supplementary Text
Captions for Supplementary Movies 1 and 2

**Other Supplementary Materials for this manuscript include the following:**

Supplementary Movies 1 and 2

**Supplementary Note 1**

Interaction of Aβ42 monomer with POPC bilayer

Immediately after interaction with the surface the secondary structure of the Aβ42 monomer changes, with a steady decline in helical content and a stepwise plateau-like increase in β-content, Supplementary Fig. 8a. The monomer starts with very little β-structure and ~12% helical content and quickly undergoes transition to conformations with 5-10% β-content, followed by an abrupt transition around ~2.5 μs to ~17% β-content that remains for the rest of the simulation. Interactions between the Aβ42 monomer and bilayer surface are transient, with each interaction occurring at different regions of the monomer, Fig. S8B. However, interactions between the Aβ monomer and the bilayer tend to be focused around the N-terminal and residues 20-30, these segments also experience the largest change in structure and adopt β-strands throughout the simulations (Fig. 2).

**Supplementary Note 2**

Dimerization on POPC membrane surface

To investigate dimer formation, a surface-bound Aβ monomer (Mon1) conformation was adopted from the Aβ42-POPC simulation, specifically the frame at ~4.9 μs, to which a free Aβ monomer (Mon2) was added at a CoM distance of 4 nm between the two molecules. The free monomer rapidly interacts with the surface-bound monomer, within ~23 ns, following which the newly formed dimer undergoes structural transition and dissociates from the membrane, Figs. 3a and Supplementary Figs. 9-10. Mon1 retains a small N-terminal β-strand throughout the simulation, while the larger central and C-terminal strands undergo dramatic change as the simulation progresses, Supplementary Fig. 9. The structure of the initially free monomer, Mon2, fluctuates widely but remains largely helical throughout the simulation.

The dimer transiently interacts with the membrane for the next ~5 μs; interaction map between the individual Aβ42 residues and the POPC headgroups further clarifies the transient nature of the membrane-interactions and reveal that interactions primarily happen at the N-terminal and central regions of the Aβ42 protein, Supplementary Fig. 10a. Normalized contact map between the two monomers revel that the interaction of monomers within the dimer primarily occurs between the central and N-terminal residues and central and C-terminal regions, Supplementary Fig. 10b. Further evaluation of contact maps at different simulation times reveal that the interaction pattern between the two monomers change as the dimer stabilizes and converges toward a β-rich conformation, Supplementary Fig. 3c. To further identify residue interactions important for dimerization we used the correlation of the pivot residue dihedral angles to the change of protein conformation, which revealed that interactions involving charged and aromatic residues (in particular residues D, H, R, Y, and I) are important for the newly formed dimer, Supplementary Fig. 10c.

Interaction of preformed dimer with POPC

To determine if a dimer interacts differently with the bilayer compared to the monomer, we simulated the interaction of a preformed Aβ dimer with POPC bilayer. In presence of POPC the dimer interacts with the bilayer surface ~185 ns after the start of the simulation, Figure 4c. Similar to the monomer, interactions with the bilayer causes change in the dimer conformation that leads to change in the orientation and secondary structures of the monomers within the dimer, Figs. 4 and Supplementary Fig. 11. Mon1 of the dimer experiences large changes in the termini, with the N-terminal adopting a small helical segment while the C-terminal loses a short β-turn; the central β-sheet remains stable throughout the simulation, Supplementary Fig. 11. Mon2 on the other hand, gains a large β-strand in the C-terminal region and a small strand in the central region. Most strikingly, the change in conformation leads to a dimer interface that is dominated by β-structure, finally culminating in an anti-parallel inter-chain β-sheet. Additionally, while the dimer-bilayer interaction is transient, periods of membrane interactions are longer compared to the monomer-POPC interactions (Figs. 4c and Supplementary Fig. 12a). Similar to the monomer, the dimer interacts with the membrane almost always through the N-terminal and residues 20-30, Supplementary Fig. 12a.

Compared to the on-surface formed dimer (Fig. 3) the preformed dimer possesses longer regions with β-structure (Supplementary Fig. 11) as well as a different interaction pattern between the monomers with in the dimer, Supplementary Fig. 12b. While interactions between the monomers do occur through the C-central and N-central regions, a significant portion also occur through the N-C regions. The difference in interaction pattern between the two dimers is also clear from the correlation of the pivot dihedrals, which places larger emphasis on charged residue interactions compared to the on-surface formed dimer (in particular D, H, and R) Supplementary Fig. 12c.

Two pathways of trimer formation on POPC

On-surface trimerization was investigated using two different pathways, a membrane-bound monomer and a free dimer (Trimer1) and a bound dimer and a free monomer (Trimer2). Simulation of the interaction between a membrane-bound monomer (Mon1) and a free dimer (Mon2 and Mon3) shows that the dimer rapidly finds the surface-bound monomer and forms a trimer, Supplementary Fig. 13a. Furthermore, in line with previous simulations, the trimer undergoes structural transitions while interacting with the membrane, this leads to the formation of long β-strands within the monomers, that throughout the simulation transitions to inter-chain β-sheet, Supplementary Figs. 13 and 14. Mon1, interacts with the dimer through both monomers in the dimer, as the simulation progresses the inter-peptide β-sheet is disrupted due to Mon3 being pushed away from the interaction interface as the trimer adopts a triangular conformation. Additionally, the trimer does not stay bound to the surface for extended amount of time, Supplementary Fig. 15a. Interactions between Mon1 and Mon2 occur through the N-terminal residue of Mon1 and the central residues of Mon2, Supplementary Fig. 15b, while interaction with Mon3 occur through the central and C-terminal regions of Mon1, Supplementary Fig. 15c. Mon2 and Mon3 interaction through central and C-terminal regions, Fig. S15D. Analysis of the pivot dihedrals reveal a residue-wise interaction pattern reminiscent of the preformed dimer, Supplementary Fig. 15e.

Addition of a free Aβ monomer (Mon3) to the membrane-bound dimer (Mon1 and Mon2) results in the formation of a trimer on the surface of the bilayer within 36 ns, Supplementary Fig 16a. Throughout the simulation, the initial dimer remains stable and does not experience significant conformational change; snapshots showing change of secondary structure of the trimer reveal that the initially free monomer undergoes significant conformational change as it interacts with the initial dimer, Supplementary Fig. 16a. Unlike previously formed oligomers however, the trimer does not experience a large degree of change in its secondary structure, Supplementary Figs. 16b and 17; in particular the monomers within the existing dimer show little change in secondary structure. As has been the case for the previous on-surface formed dimer and trimers, the newly formed trimer only transiently interacts with the bilayer through the N-termini of the monomers, Supplementary Fig. 18a. While the trimer interacts with the membrane, the monomers within interact with each other as follows: Mon1 and Mon2 interact through N-C and C-central regions while Mon1 and Mon3 interact only through the residues in the C-central segments. Mon2-Mon3 interactions are facilitated through the central segments of the monomers with contributions from N-C interactions. The free monomer, Mon3, interacts with the dimer through its central residues, in particular the KLVFFA residues, Supplementary Fig. 18 b-d. Analysis of individual residue interactions do not show large differences between Trimer1 and Trimer2, Supplementary Fig. 18e.

Tetramer formation on POPC surface

The interaction of two dimers, a surface-bound (Mon1 and Mon2) and a free dimer (Mon3 and Mon4), results in the formation of a tetramer in presence of POPC bilayer, Supplementary Fig. 19. Unlike the formation of previous oligomers, the tetramer forms off surface following the dissociation of the bound dimer. The tetramer is re-organized and the monomers undergo change, both in conformation and secondary structure, Supplementary Figs. 19 and 20. Mon1 and Mon2, largely maintains their initial structure and only shows dynamic behavior in the N-termini of the monomers within the dimer. Whereas the initially free dimer, Mon3 and Mon4, is more dynamic, in particular Mon4 experiences significant change in secondary structure adopting small β-structure and helical segments before finally assuming a conformation with a small C-terminal β-strand, Fig. S20. Once formed, the tetramer interacts with the bilayer surface through two of the monomers, these monomers work as anchors and allow the tetramer to adopt an extended conformation, Supplementary Fig 21a. Consequently, the monomers interacting with the surface experience the greatest amount of change. Moreover, like previous oligomers the tetramer only transiently interacts with the surface and undergoes many association-dissociation events. The monomers within the tetramer largely interact with each other through the terminal and central regions, with frequent interactions occurring with their initial monomer partners (i.e. Mon1-Mon2 and Mon3-Mon4), Supplementary Figs. 21b-g.

**Supplementary Note 3**

Interaction of Aβ42 with POPS bilayer

We investigated the interactions between Aβ42 and POPS bilayer. Aβ42 monomer rapidly interacts with POPS and undergoes structural transition into a conformation with β-structure, Supplementary Fig. 22a. The monomer has a smaller degree of β-structure compared to the monomer interacting with POPC, Supplementary Fig. 22 a-b. The monomer interacts frequently with the bilayer, with the majority of interactions being transient, Supplementary Fig. 23a, however stable binding (around 2.4 μs for ~600 ns duration) to the bilayer is also observed.

Additionally, interactions are primarily through the N-terminal and residues 23-33, Supplementary Fig. 23b.

Dimerization of Aβ42 on POPS

Interaction between a POPS-bound (Mon1) and a free (Mon2) Aβ42 monomer happens after ~520 ns and results in a stable dimer, Supplementary Fig. 24a. The formation of the dimer causes a dramatic increase of the β-structure content and a decrease in the helical content, Supplementary Fig. 24b. This change is further enhanced around ~2.6 μs and results in a final β-structure content of ~20%. Mon1 maintains its structure with little change for the first ~1.8 μs of the simulation but then loses the N-terminal β-strand while the central and C-terminal strands become smaller, Supplementary Fig. 25. During the same time, after the dimer has formed, Mon2 undergoes a helix-to-strand conversion that starts with the loss of the N-terminal and central helix, followed by appearance of a C-terminal helix and β-strand and an N-terminal strand. The newly formed strands remain stable for the remaining simulation time, while the C-terminal helix is converted into two additional strands in the central and C-terminal regions, Supplementary Fig. 25. The dimer transiently interacts with the POPS bilayer through the N- and central regions of a single monomer, Supplementary Fig. S26a. Similar to the monomer, the dimer experiences a prolonged interaction (~ 1.6 μs duration) with the membrane toward the end of the simulation (Supplementary Fig. 26a), which sees the dimer adopt higher β-structure content as well as a small transient helix, Supplementary Fig. 25. The dimer is stabilized primarily by N-C and C-C terminal interactions, Supplementary Fig. 26b. The pivot dihedral analysis yields a similar picture as the on-surface dimerization on POPC, with emphasis on charged and hydrophobic residues being important for the dimer stability, Supplementary Fig. 26c.

Interaction of preformed Aβ42 dimer with POPS

During the interaction of an already formed dimer with the POPS bilayer, the dimer undergoes structure transitions with both monomers within the dimer experiencing an increase in β-structure, Supplementary Fig. 27. Which leads to a dimer interface with inter-peptide β-sheet. Surface interactions are transient and primarily involve residues in the N-terminal and central regions of the proteins, Supplementary Fig. 28a. Interactions between the monomers within the dimer are primarily focused around the residues of N-C termini and central-central region, Supplementary Fig. 28b. Residue correlation map shows that the dimer is stabilized by interactions involving residues A, D, E, F, I, K, L, and N, Supplementary Fig. 28c.

Aβ42 trimers on POPS

Similar to the POPC bilayer, trimer formation was explored through two pathways. On-surface formation of a trimer on POPS, from a membrane-bound monomer (Mon1) and a free dimer (Mon2 and Mon3), occurs rapidly, Supplementary Fig. 29a, with the monomers experiencing increase in β-structure content, Supplementary Fig. 29b. Mon1, does not experience a significant change in structure as it maintains the small β-strands in the termini and central region, Supplementary Fig. 30. Mon2 maintains three small b-strands in the central and C-terminal regions while it experiences fluctuations in the secondary structure of the N-terminal region. Mon3 is the most dynamic monomer in the system and experiences significant changes in all regions, however it also has short β-strands in the C-terminal and central (17-21) regions, Supplementary Fig. 30.

After assembly, the Aβ42 trimer dissociates from the surface and for the majority of the simulation transiently interacts with the bilayer through a few residues in the N-termini of the monomers within the trimer, Supplementary Fig. 31a. The monomers within the trimer interact through terminal residues as well as central residues: Mon1-Mon2 interact through their N- and C- termini; Mon1-Mon3 interact almost exclusively through C-C terminal interactions; while Mon2-Mon3 interact through the central regions as well as the N- and C-termini, Supplementary Fig. 31 b-d. The residue-wise interactions important for the stability of the trimer resembles the previously observed trimers formed on POPC with emphasis on H, K, Q, S, and Y, Supplementary Fig. 31e.

A POPS-bound dimer (Mon1 and Mon2) rapidly interacts with the free monomer (Mon3) and forms a trimer, Supplementary Fig. 32a. The newly formed trimer undergoes secondary structure transition with a gradual increase of β-structure elements, Supplementary Figs. 32b and 33, during which the trimer experiences change in the organization of the monomers. Unlike on POPC and the previous trimer on POPS, the change in the secondary structure is significant and for two of the monomers means an increase in β-structure content, Supplementary Fig. 33. Interaction with the POPS bilayer is transient for the majority of the simulation, however between ~2.2-4.4 μs the monomer remains on the surface and interacts with the membrane through the N-termini of Mon1 and Mon3 and to a lesser extent the C-terminal residues of Mon2, Supplementary Fig. 34a. The interaction of the monomers within the trimers are significantly different compared to the trimers formed by the first pathway: Mon1-Mon2 interactions are facilitated by the central regions of the monomer as well as the C-termini. Mon1-Mon3 interaction happen through N-C interactions and N-central regions, while Mon2-Mon3 interactions occur primarily through the N-C termini of the monomers, Supplementary Figs. 34b-d. Charged and aromatic residues (A, D, F, H, S, and Y) are the primary contributors to the stability of the trimer, Supplementary Fig. 34e.

Tetramer formation on POPS

In the presence of POPS the surface-bound dimer (Mon1 and Mon2) dissociates from the bilayer surface and interact with the initially free dimer (Mon3 and Mon3), which has become membrane-bound, to form a tetramer, Supplementary Fig. 35. After formation both dimers undergo structural transition, initially small segments within the N-termini of the Mon3 and Mon4 transition to α-helices, while the monomers Mon1 and Mon2 tend toward increased β-structure content, Supplementary Fig. 35a. The β-content increases a small amount while the helical content slowly decreases as the simulation progresses, Supplementary Fig. 35b. As the stable tetramer is formed, the transition in structure tend toward helical for the N-termini of two monomers, Mon2 and Mon4, while the C-termini tend toward β -structures; in particular the monomers Mon1 and Mon2 form stable anti-parallel β-sheets, Supplementary Fig. 36. As seen with the other oligomers, the tetramer only sporadically interacts with the membrane surface, Supplementary Fig. 37a, with the residues involved being from the N-terminal and central regions. Interactions within the tetramer occur through residues in the termini and the central regions, with high probability being given to the N-C and central-central residues, Supplementary Figs. 37 b-g. Mon1-Mon2 interactions primarily happen through C-C termini and the central segments; Mon1-Mon3 interact through N-C- and C-C termini; Mon1-Mon4 interact through a small number of residues in the N- and C-termini and C-C termini; Mon2 interacts with Mon3 through residues in the N-terminus and central segment and with Mon4 through C-C terminal interactions. Mon3-Mon4 interactions are dominated by residues from the central regions of the monomers and C-central interactions

**Supplementary Movie 1. Interaction of Aβ42 monomer with the POPC bilayer surface.** The bilayer surface facilitates the transition of the protein structure from a conformation with small β-structure content to a conformation with an extended β-sheet consisting of two strands, which remains stable for the rest of the simulation.

**Supplementary Movie 2. Interaction of membrane-bound Aβ42 monomer with a free monomer on POPC.**

Transient interaction between the initially free monomer and the membrane surface are observed, but the monomer remains free, which is followed by diffusion above the bilayer toward the bound Aβ42 monomer. After a short period, the two monomers interact and for a dimer. The conformation of both monomers changes dramatically during the on-surface dimerization.

### Supplementary Figures for

#### Lipid membranes trigger misfolding and self-assembly of amyloid β 42 protein into aggregates

Siddhartha Banerjee[1,†], Mohtadin Hashemi[1,†], Karen Zagorski[1] and Yuri L. Lyubchenko[1,*]

†These authors contributed equally

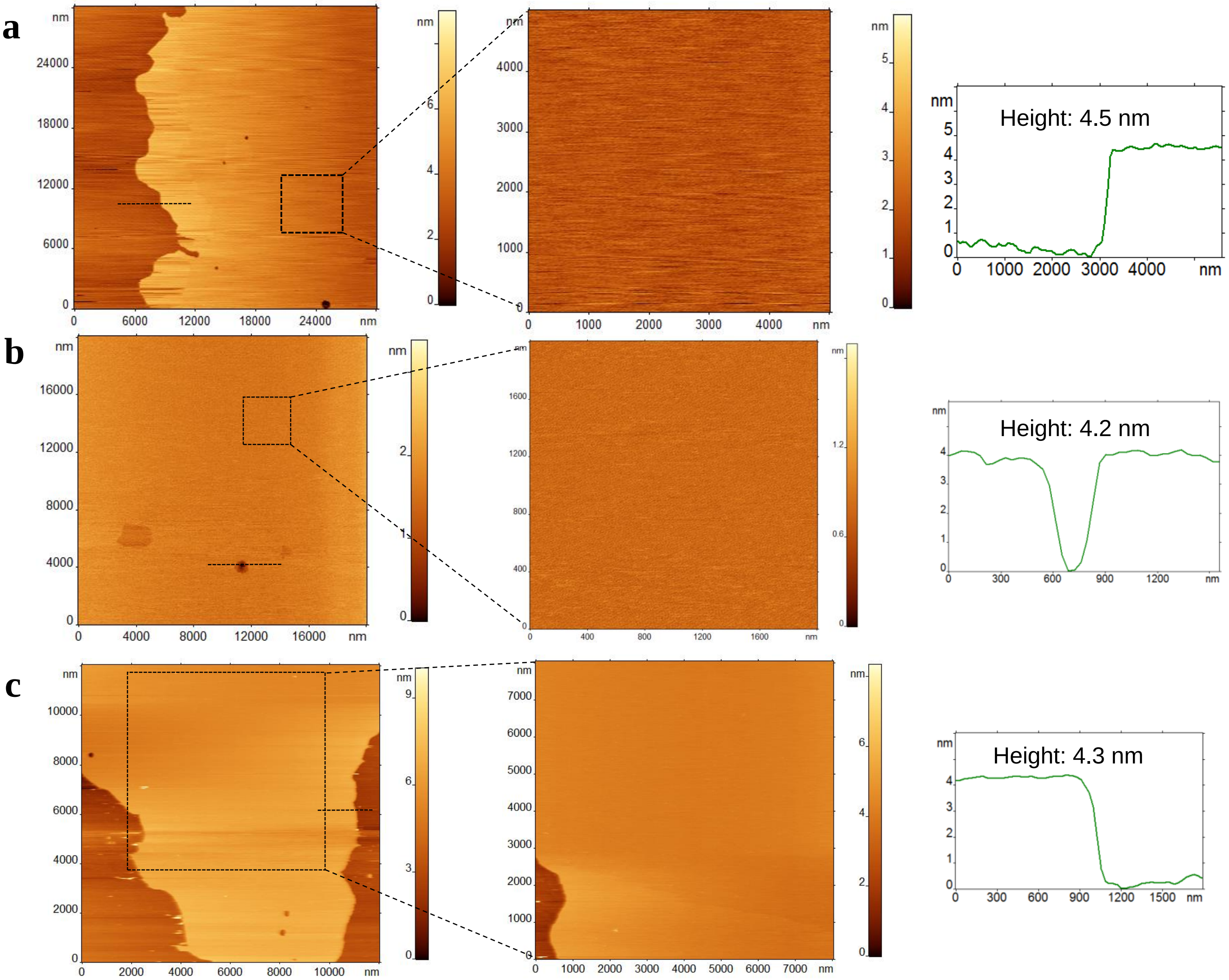


**Supplementary Figure 1. Representative AFM images of supported lipid bilayers used in this study.** AFM topographic images showing large, smooth SLB surfaces of (**a**) POPC, (**b**) POPS and (**c**) POPC-POPS mixture (1:1 mol). The first column shows the large scans and scans for zoomed area have been shown in the second column. The SLB surfaces are smooth and devoid of any unruptured or trapped vesicles. Third column shows the cross-section profiles, which indicate the height of the prepared bilayer. The height of the bilayer remains 4.2-4.5 nm.

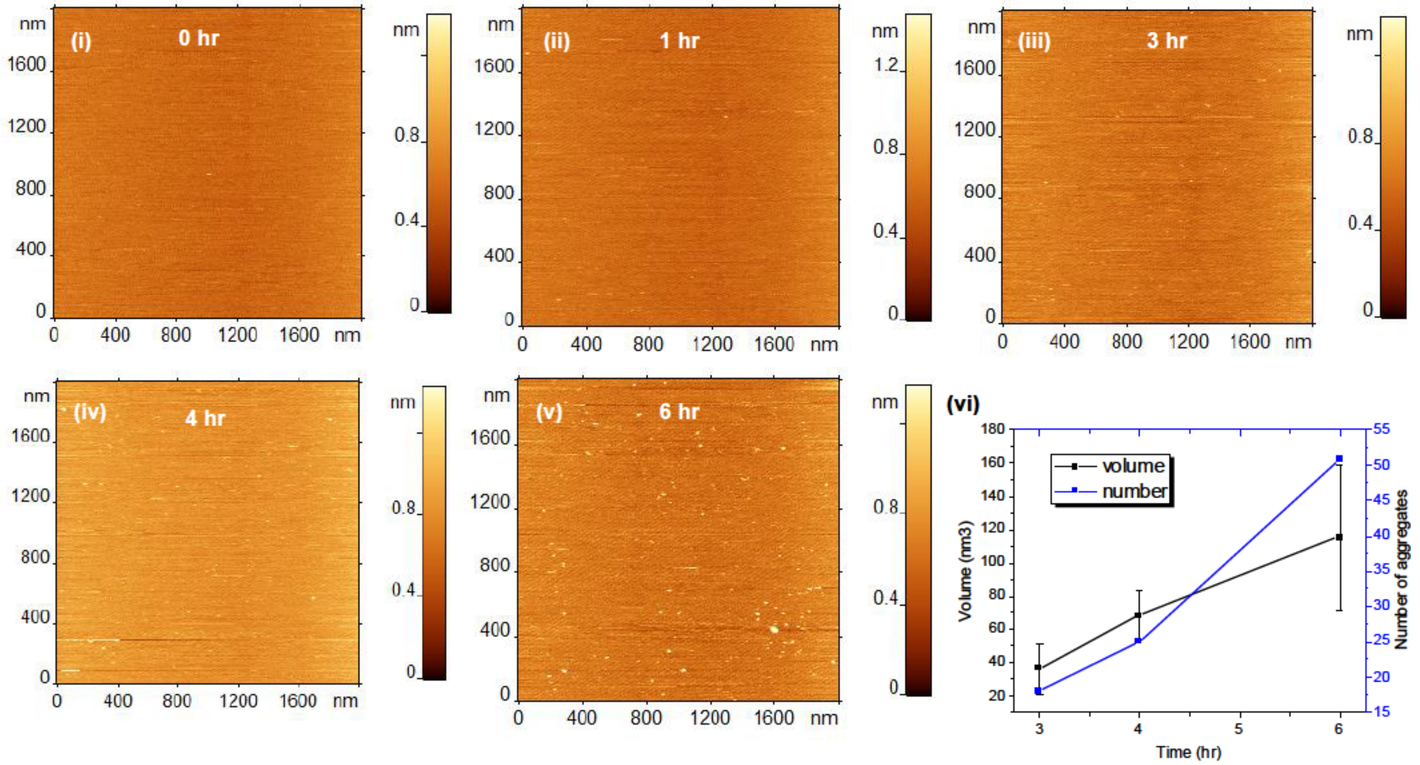


**Supplementary Figure 2. Time-lapse AFM imaging of 10 nM Aβ42 on POPC SLB, in presence of 150 mM NaCl.** Frames (i)-(v) show images acquired at different points. The features on the surface are Aβ42 aggregates. Initial frame (Frame i) shows clean surface. The formation of aggregates is seen in consecutive frames as time progresses. Frame (vi) represents the number and volume of aggregates as a function of time.

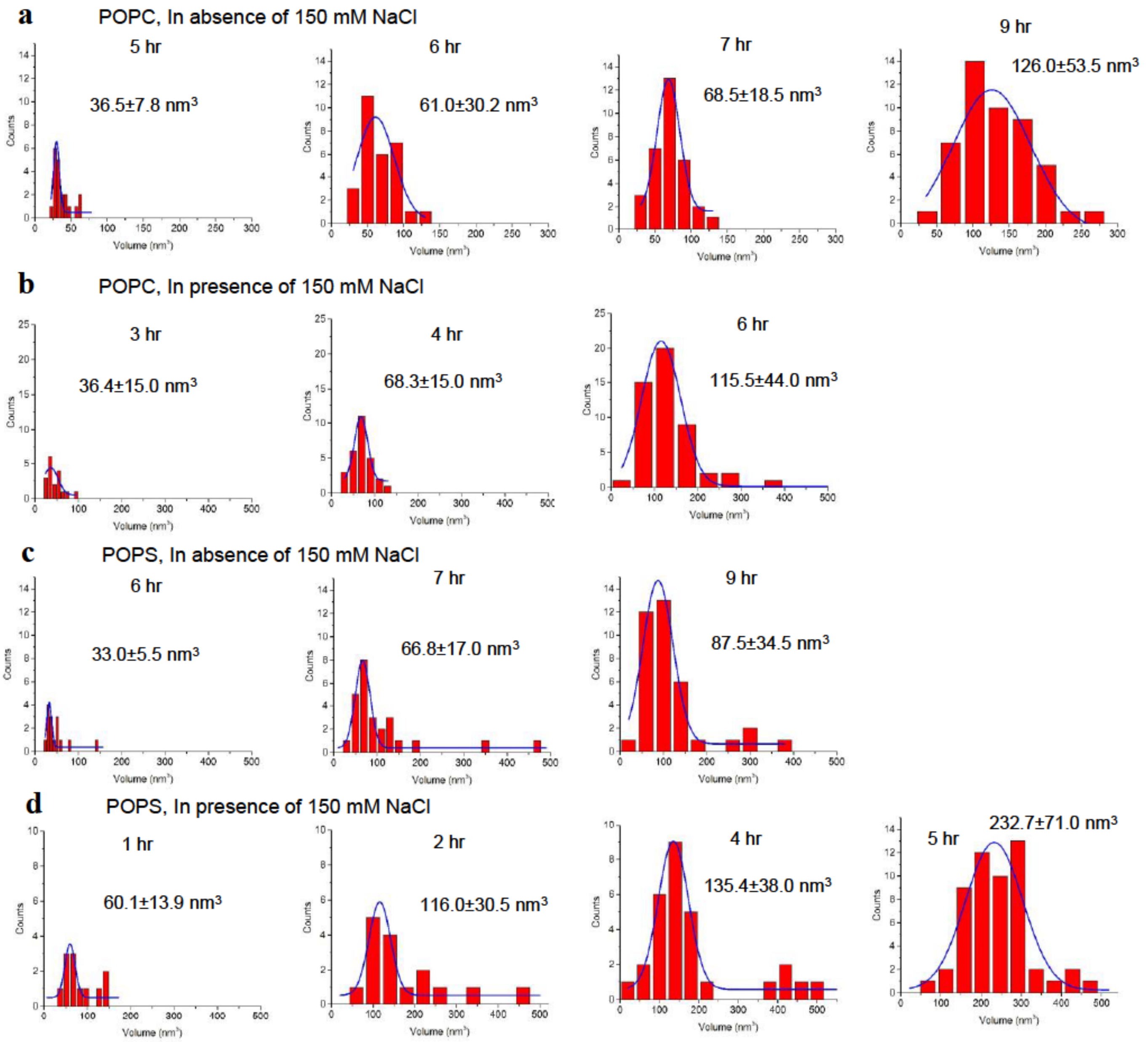


**Supplementary Figure 3. Quantitative analysis of 10 nM Aβ42 aggregation on POPC and POPS SLBs in absence or presence of 150 mM NaCl.** (**a**) and (**b**) show the volume distribution of the aggregates formed at different time points on the POPC SLBs in the absence and presence of 150 mM NaCl, respectively. (**c**) and (**d**) show the aggregate volumes on POPS SLBs in the absence and presence of 150 mM NaCl, respectively.

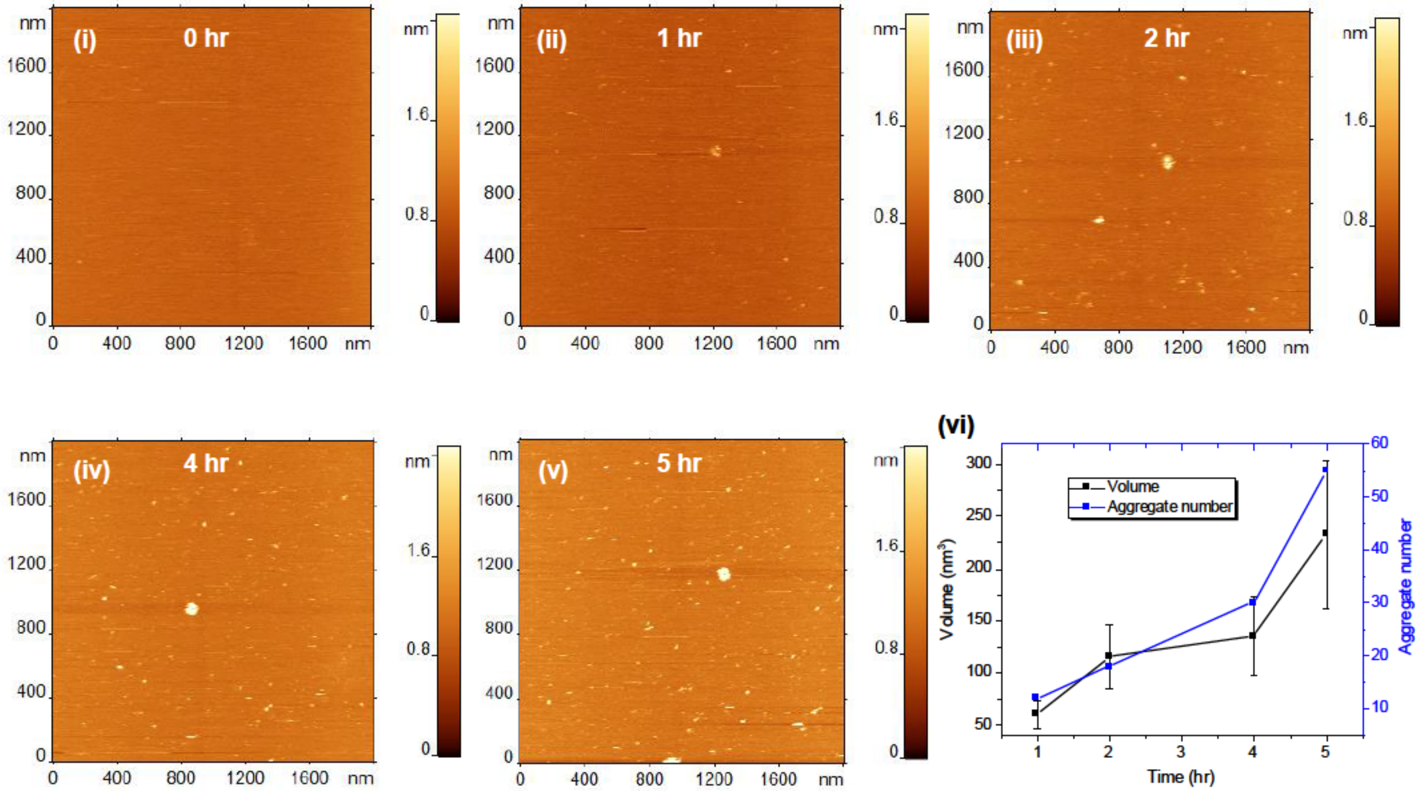


**Supplementary Figure 4. Time-lapse AFM imaging of 10 nM Aβ42 on POPS SLB, in presence of 150 mM NaCl.** Frame (i)-(v) shows the images taken at different points. The blobs on the surface are Aβ42 aggregates. Frame (i) shows clean surface. Aggregates start forming in the consecutive frames. Frame (vi) shows the plot for number and volume of aggregates as a function of time. Both the number and volume of the aggregates increase with time.

**a** Aggregation on POPC-POPS SLB in absence of 150 mM NaCl

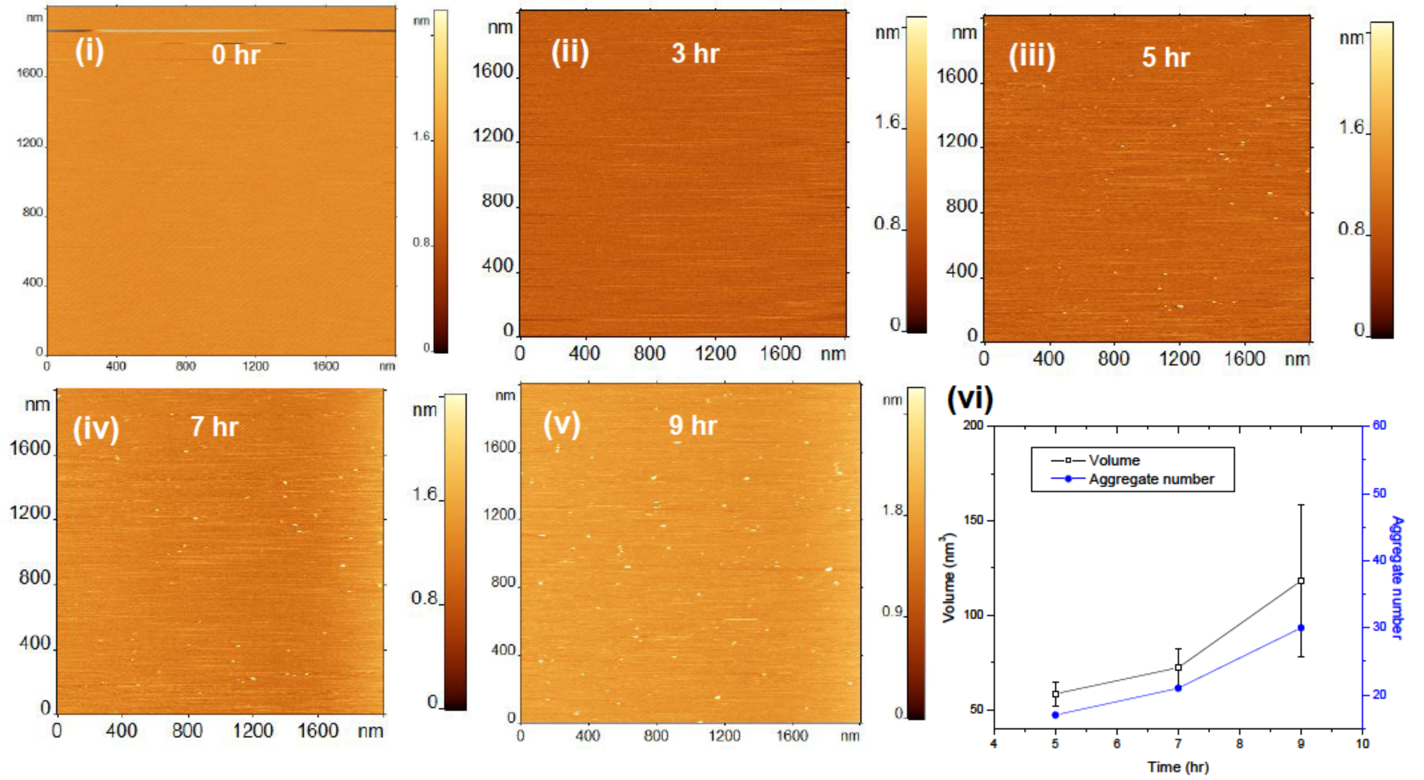


**b** Aggregation on POPC-POPS SLB in presence of 150 mM NaCl

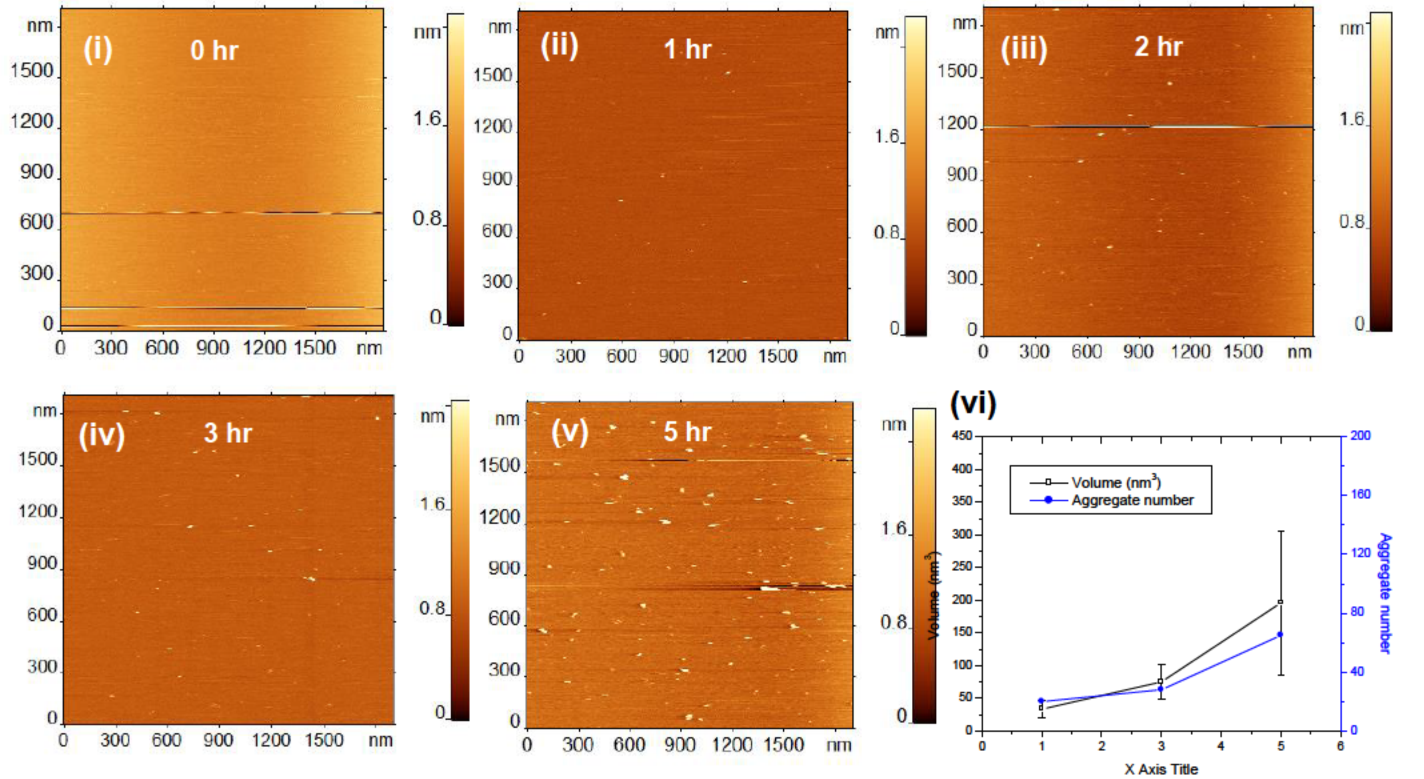


**Supplementary Figure 5. Time-lapse AFM images of 10 nM Aβ42 on POPC-POPS SLB.** Images recorded (**a**) in the absence and (**b**) in presence of 150mM NaCl, respectively. Frames (i)-(v) show the SLB surface at different time points. Frame (vi) shows the plot for count and volume of aggregates as a function of time. Aggregates start appearing much faster (after 1 hr of incubation) in presence of NaCl (Frame ii in b). After 5 hr of incubation, surface contains more aggregates in presence of NaCl (Frame v in b), compared to no NaCl condition (Frame iii in b).

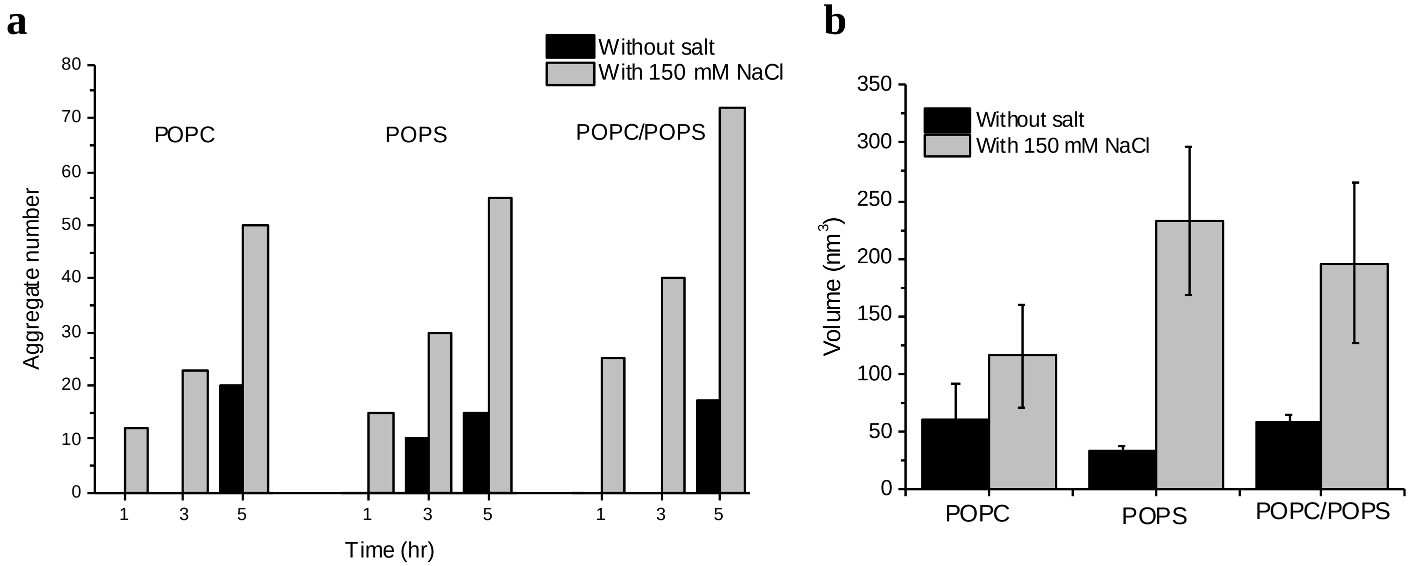


**Supplementary Figure 6. Volume and number of aggregates formed on different bilayer surfaces.** (**a**) Comparison of number of aggregates formed after 1 hr, 3 hr and 5 hr of incubation with NaCl (gray bars) and without NaCl (black bars) on three SLB surfaces. (**b**) Comparison of the volume of aggregates formed on POPC, POPS and POPC-POPS SLBs with and without salt condition after 5 hr of incubation. The volume of the aggregates are much larger in presence of NaCl in all three surfaces.

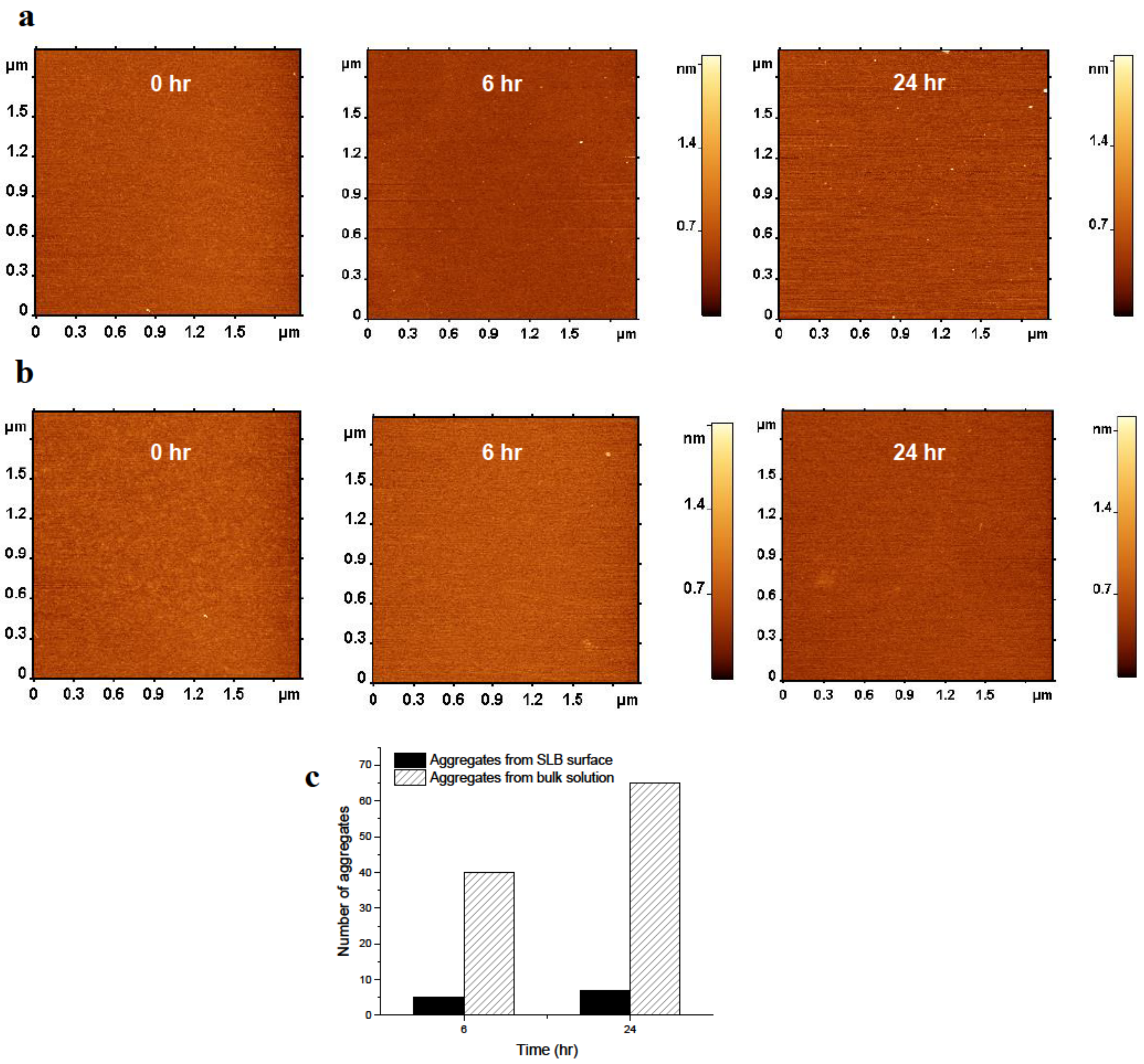


**Supplementary Figure 7. Bulk experiment showing the dissociation of the aggregates from the SLB surface.** AFM images of Aβ42 aggregates present in bulk solution at different time intervals (**a**) in presence of POPS SLB surface and (**b**) in absence of SLB surface. AFM images clearly show the presence of higher number of aggregates at 24 hr panel in the presence of SLB surface compared to its absence. (**c**) The bar diagram shows the comparison in the number of aggregates present in two situations. The numbers are computed from two 2 μm × 2 μm images for each time point.

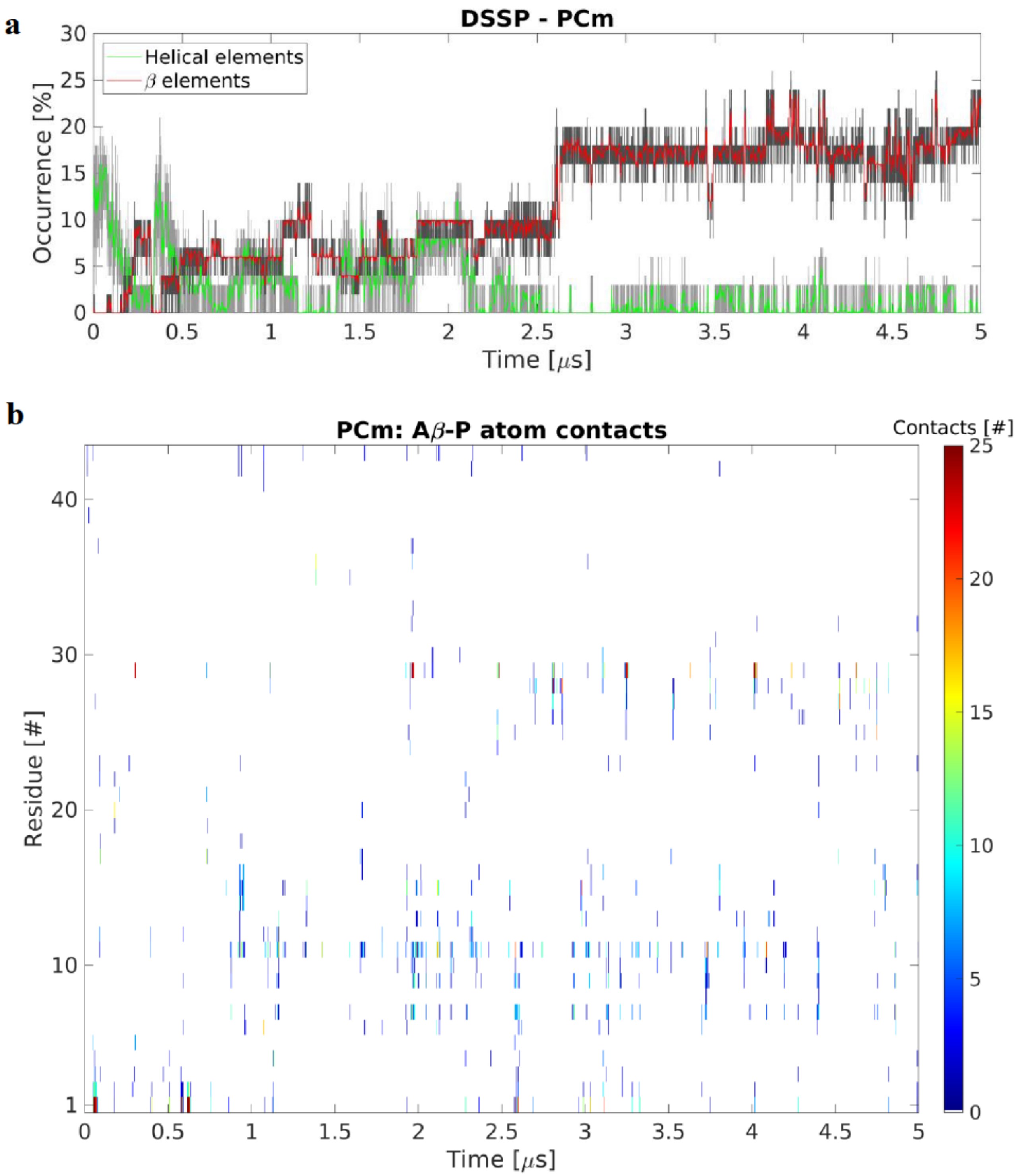


**Supplementary Figure 8**. **Interaction of Aβ42 monomer with POPC bilayer (PCm system)**. (**a**) Evolution of secondary structure β- (sheet and bridge), red, and helical (α-, π–, and 3/10-helices), green, elements of the Aβ42 protein as determined by DSSP (v3). The monomer starts with very little β-structure and ~12% helical content and quickly undergoes transition to conformations with 5-10% β-content, followed by an abrupt transition around ~2.5 μs to ~17% β-content that remains for the rest of the simulation. The graphs are moving averages using a 1 ns window; raw data is presented as dark and light grey graphs, respectively. (**b**) Time-resolved contact map between residues of the Aβ42 monomer and the P atoms of the POPC headgroups. Interactions are focused around the N-terminal and residues 20-30. Color represent total number of contacts (<6 Å).

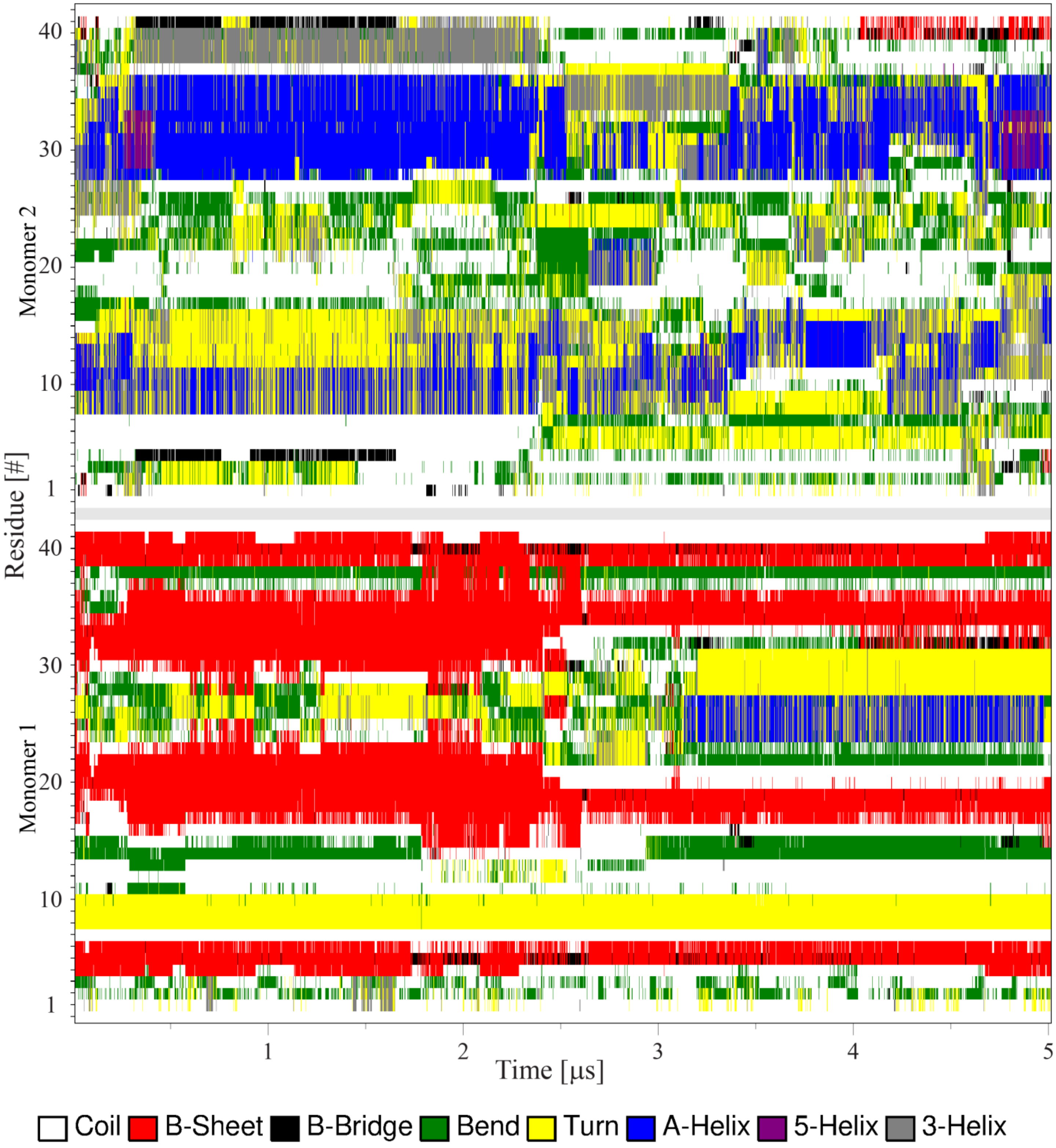


**Supplementary Figure 9. Conformational analysis of the interaction of a membrane-bound and a free Aβ42 monomer with POPC bilayer (PCmm system).** Map of protein secondary structure versus time as determined by DSSP (v3). The initially membrane-bound monomer (Mon1) retains a small N-terminal β-strand throughout the simulation, while larger central and C-terminal strands undergo dramatic change as the simulation progresses. The structure of the initially free monomer, Mon2, fluctuates widely but remains largely helical throughout the simulation.

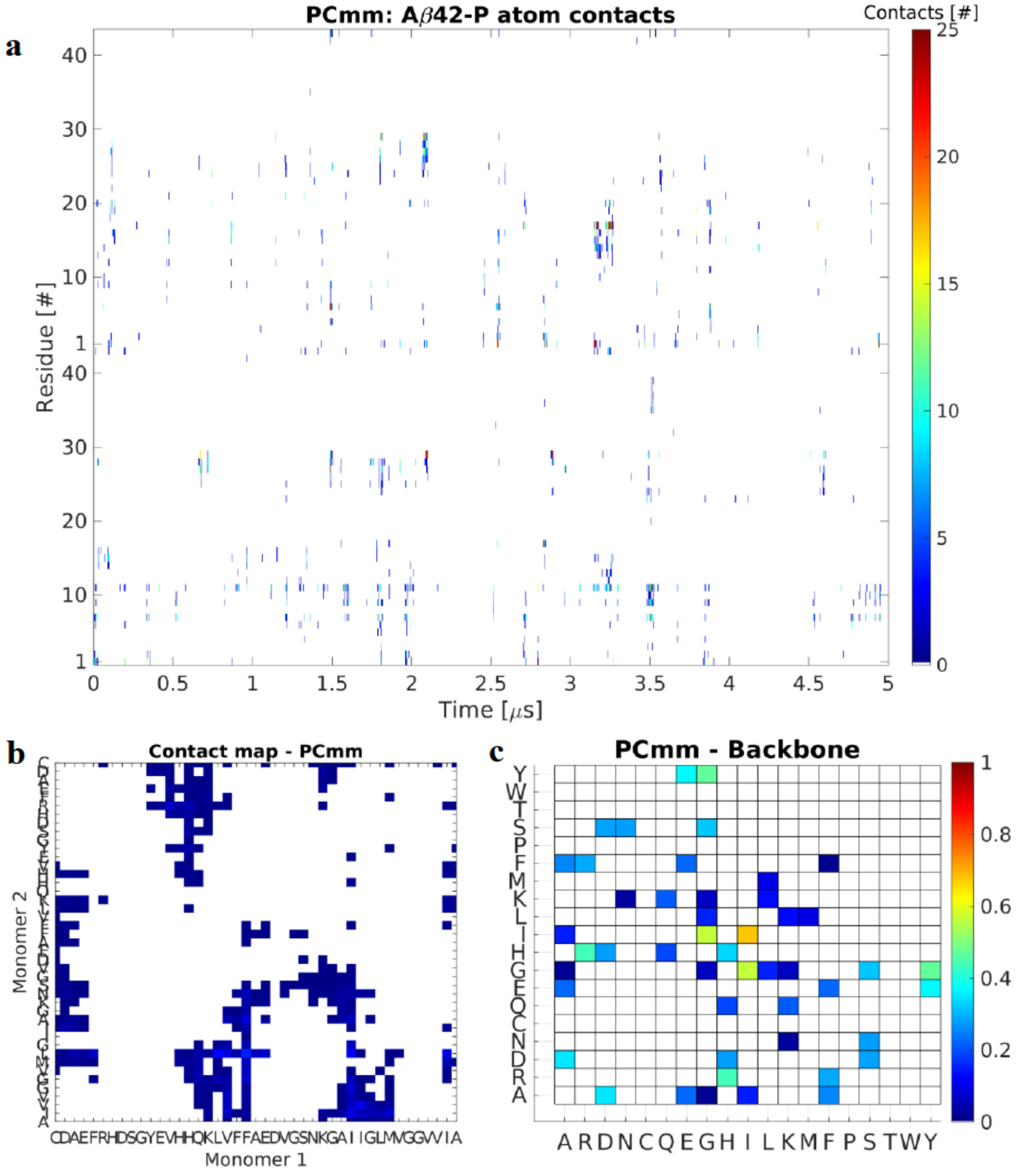


**Supplementary Figure 10. Analysis of contacts during the interaction of membrane-bound and free Aβ42 monomers with POPC bilayer (PCmm system).** (**a**) Interaction map between protein residues and P atoms of the lipid headgroups. Similar to the monomer interactions, the on-surface assembled dimer interacts with the membrane through the N-termini and residues 20-30 of the monomers within the dimer. The monomers within the dimer rarely interact extensively with the membrane at the same time. (**b**) Normalized contact map based on the residue contacts of individual residues interacting with the opposite monomer. Mon1 central and C-terminal regions interact extensively with the central and C-terminal regions of Mon2; color represent total number of contacts (<6 Å) between the residue pairs. (**c**) Pivot dihedral correlation map showing the normalized residue-wise interactions important for stability of dimer. Interactions involving charged and hydrophobic residues are the most important for the dimer stability, in particular involving D, H, R, Y, and I.

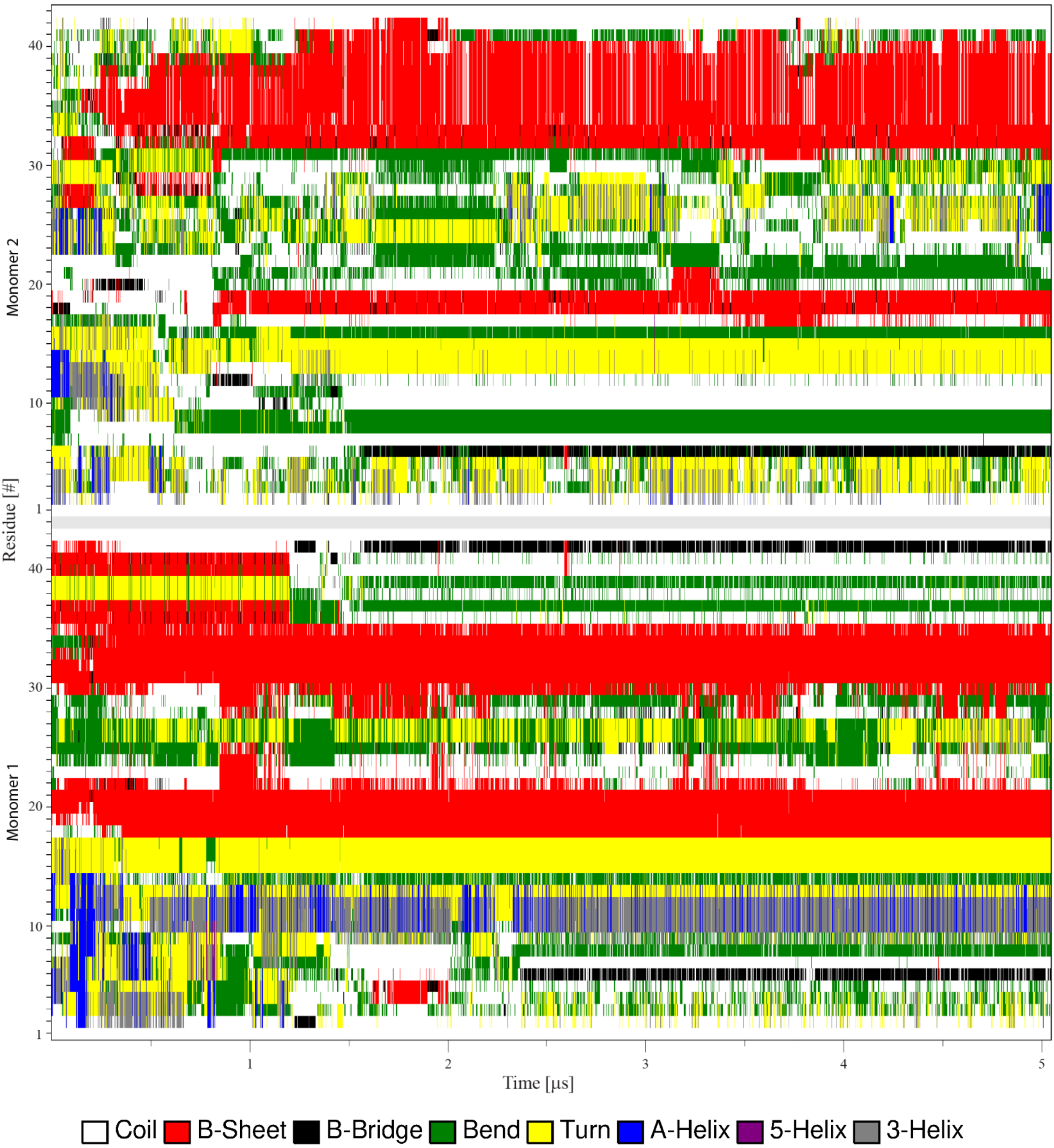


**Supplementary Figure 11. Conformational analysis of the interaction of Aβ42 dimer with POPC bilayer (PCd system).** Map of protein secondary structure versus time as determined by DSSP (v3). Mon1 experiences large changes in the termini, N-terminal adopting a small helical segment while the C-terminal loses a short β-turn; the central β-sheet remains stable throughout the simulation. Mon2 on the other hand, gains a large β-strand in the C-terminal region and a small strand in the central region.

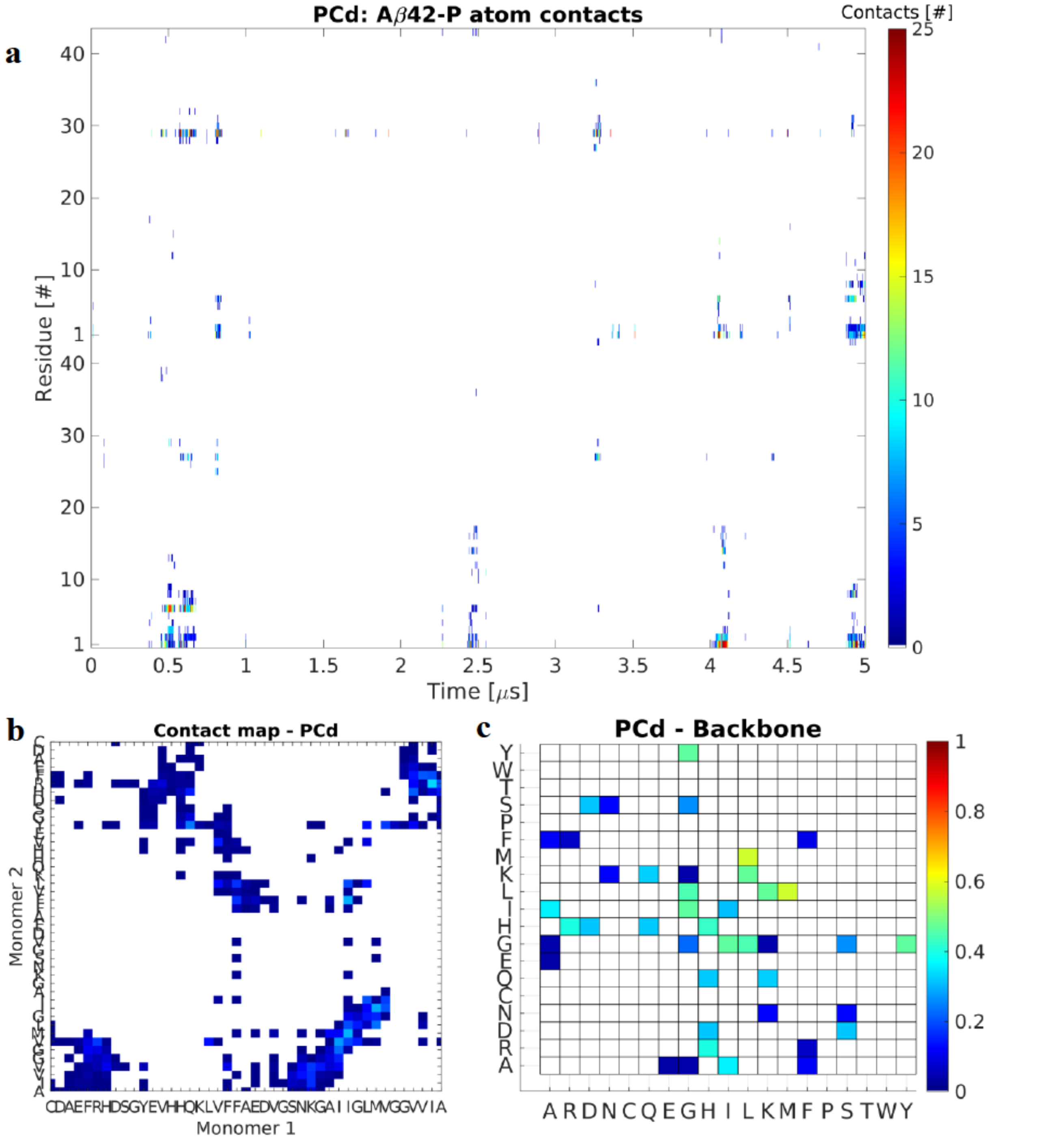


**Supplementary Figure 12. Analysis of the interaction of an Aβ42 dimer with POPC bilayer (PCd system).** (**a**) Time-resolved map of interactions between individual residues and the PC headgroups. Color represent total number of contacts (<6 Å) between the residue and the P atoms of the headgroups. The dimer sporadically interacts with the membrane, almost always through the N-terminal and residues 20-30. (**b**) Normalized map, based on the residue contacts, between residue of one monomer with the opposite monomer. The monomers interact through N-C, C-C, and central regions. Color represent total number of contacts (<6 Å) between the residue pairs. (**c**) Pivot dihedral correlation map showing the normalized residue-wise interactions important for stability of dimer. Interactions involving residues D, H, L, M, R, and Y are particularly important for the stability of the dimer, while other interactions (such as through residues I, Q, and S) are frequent but not as important.

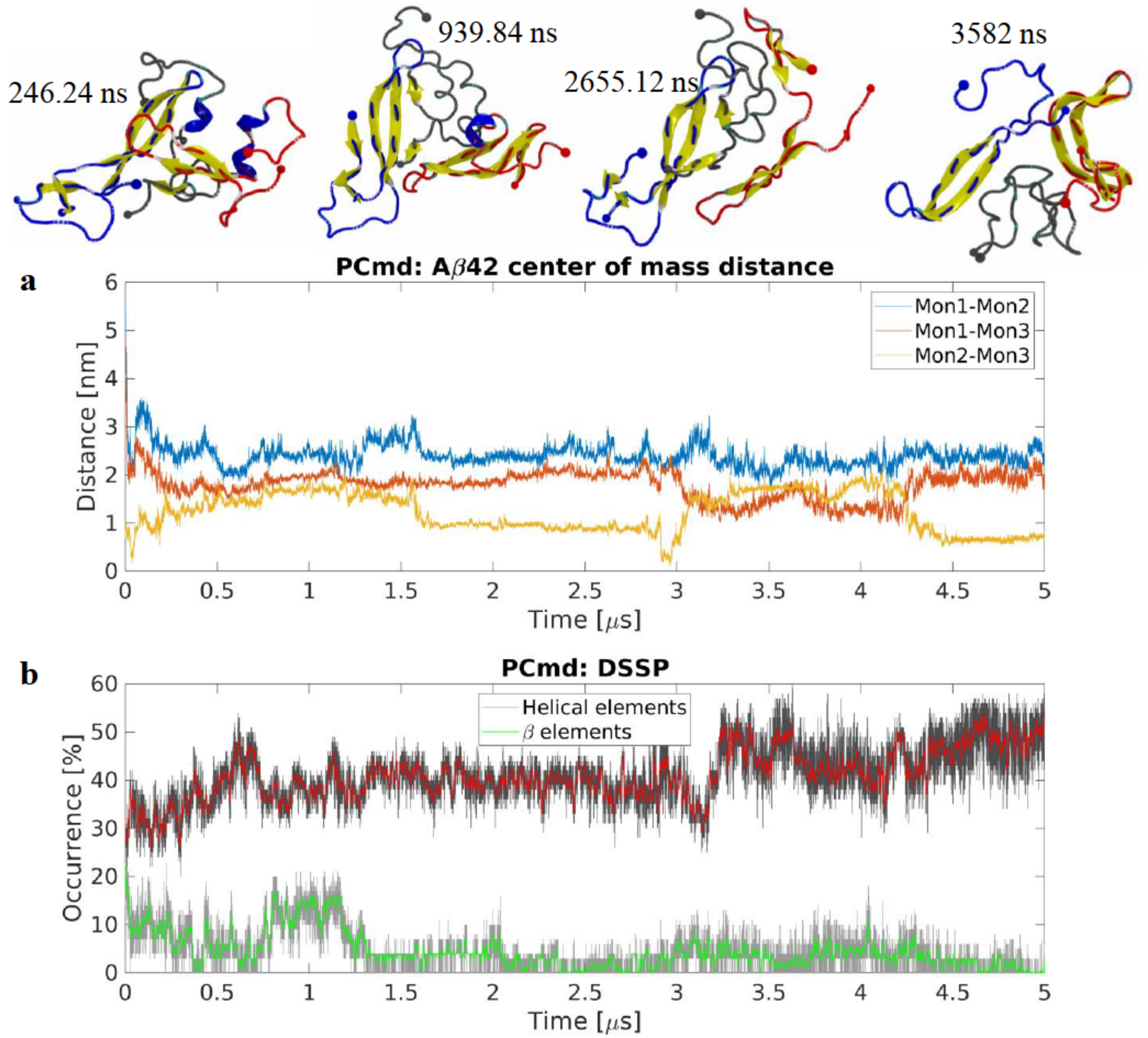


**Supplementary Figure 13. Trimer formation on POPC from membrane-bound Aβ42 monomer and free dimer (PCmd system).** (**a**) Center of mass (CoM) distances between monomers showing the distances between the surface-bound monomer, Mon1, and the free dimer, Mon2 and Mon3. The trimer rapidly forms and the initially free dimer is re-arranged through the interactions of Mon1, seen as increased Mon2-Mon3 CoM distance; as the newly formed trimer stabilizes it undergoes conformational change, seen as change in the CoM distances. Snapshots show the change of secondary structure of Aβ42 monomers within the trimer. The initially surface-bound monomer, Mon1, interacts with the dimer through both monomers in the dimer, as the simulation progresses the interaction involves an inter-peptide β-sheet that is then disrupted due to Mon3 being pushed away from the interaction interface as the trimer adopts a triangular conformation. Molecules are in cartoon representation following VMD coloring scheme (yellow β-strands and purple α-helices), N- and C-terminal Cα are represented as a large and a small sphere respectively; blue, red, and grey sphere and backbone colors represent Mon1, Mon2, and Mon3 respectively. (**b**) Evolution of secondary structure β- (sheet and bridge), red, and helical (α-, π–, and 3/10-helices), green, elements as determined by DSSP (v3). The trimer experiences a significant increase in β-content as the simulation progresses. The graphs are moving averages using a 1 ns window; raw data is presented as dark and light grey graphs, respectively.

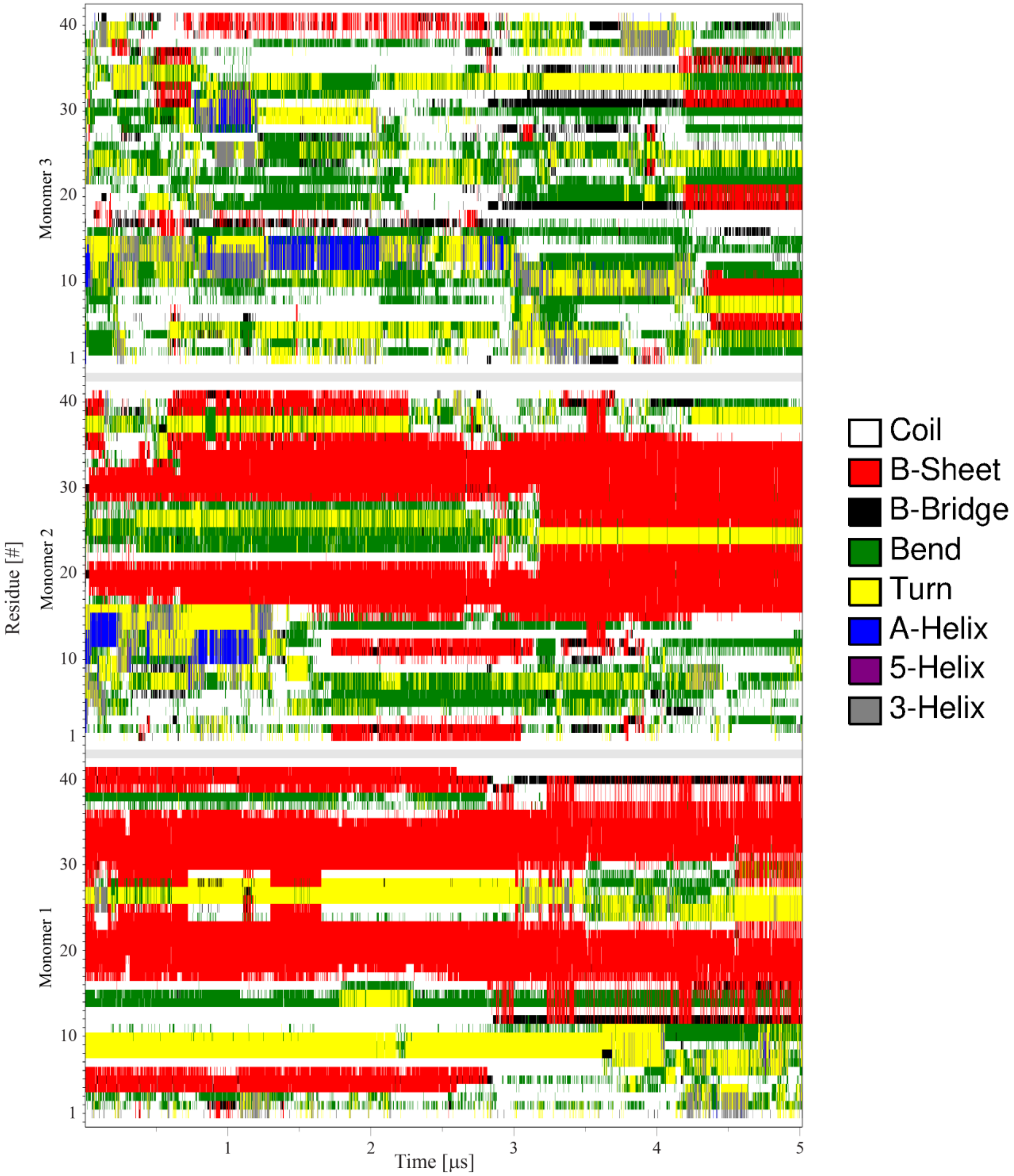


**Supplementary Figure 14. Conformation analysis of trimer formation on POPC, from a membrane-bound Aβ42 monomer and a free dimer (PCmd system).** Time-resolved map of the protein secondary structure for the initially membrane-bound Monomer, Mon1, and the initially free dimer, Mon2 and Mon3. Mon1 experiences little to no change in secondary structure for the majority of the simulation; around ~3 µs and onward the small N-terminal β-strand is disrupted and the central and C-terminal β-strands grow larger. Mon2 initially loses the small C-terminal β-strand and a helical segment appears at residues 10-15, this rapidly changes and the central and C-terminal strands become larger at the same time as the strands of Mon1 become larger. Mon3 on the other hand remains largely free of β-strands and helices throughout the simulation and adopts turn and bend structures in all segments; towards the end of the simulation small β-strands form in the N-terminal, central, and the C-terminal regions of the monomer.

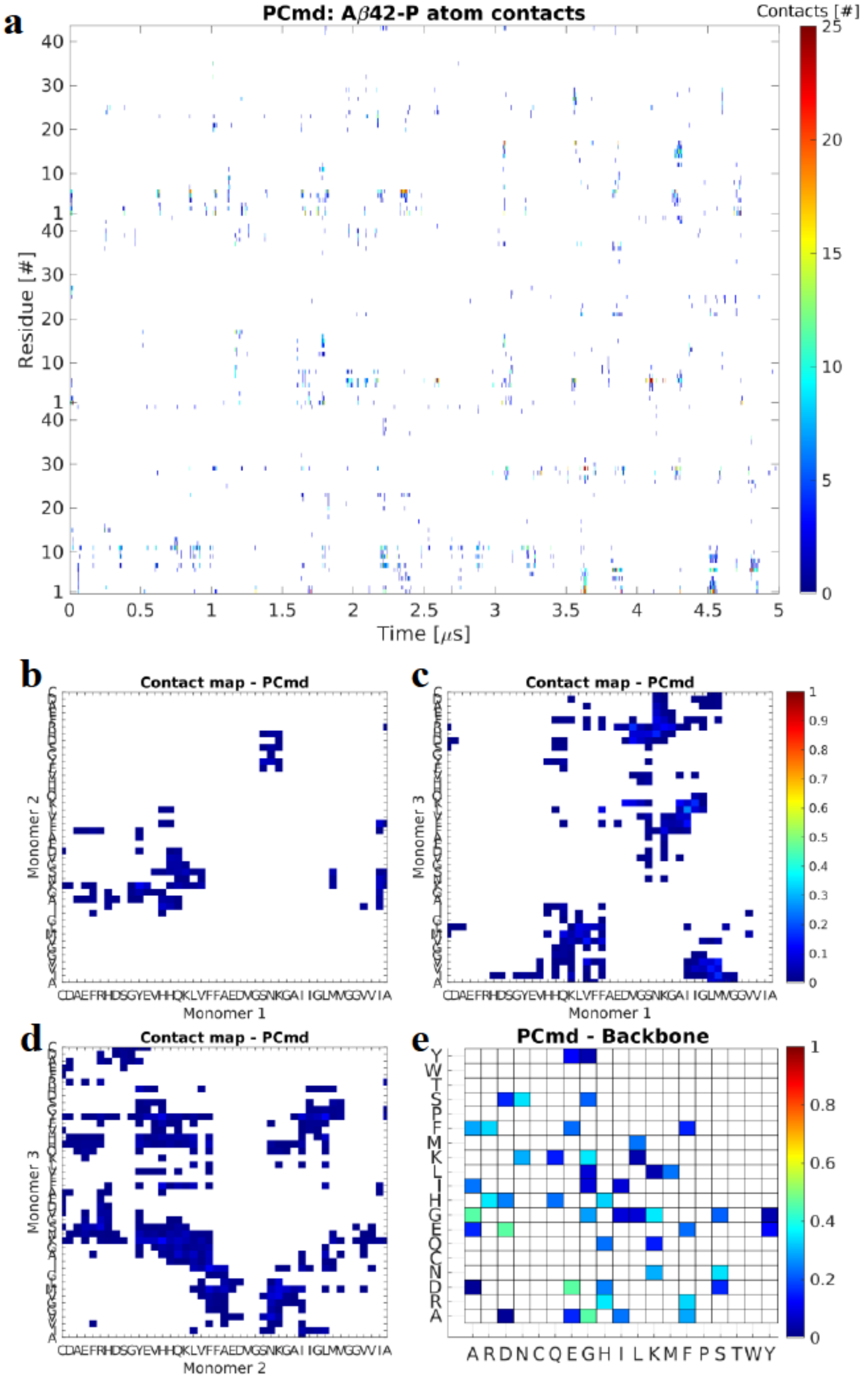


**Supplementary Figure 15. Analysis of contacts during trimer formation on POPC from membrane-bound Aβ42 monomer and free dimer (PCmd system).** (**a**) Time-resolved map of interactions between individual residues and the P atoms of the lipid headgroups. The Aβ42 molecules interact transiently with the membrane with the majority of interactions happening through the N-terminal residues. Color represent total number of contacts (<6 Å) between the residue and the headgroups. (**b-d**) Normalized map, based on the residue contacts, between residue of one monomer with the opposite monomer in the PCmd system. Majority of the interactions between Mon1 and Mon2 occur through the central regions of the proteins; Mon1 and Mon3 interact through N-C and the central regions with majority of the interaction focused around KLVFFA segment; Mon2 and Mon3, the original dimer, interact through all segments of the protein with majority of interactions happening in the central region. Color represent total number of contacts (<6 Å) between the residue pairs. (**e**) Pivot dihedral correlation map showing the normalized residue-wise interactions important for stability of the trimer. Interactions involving residues A, D, E, K, N, and R are particularly important for the stability.

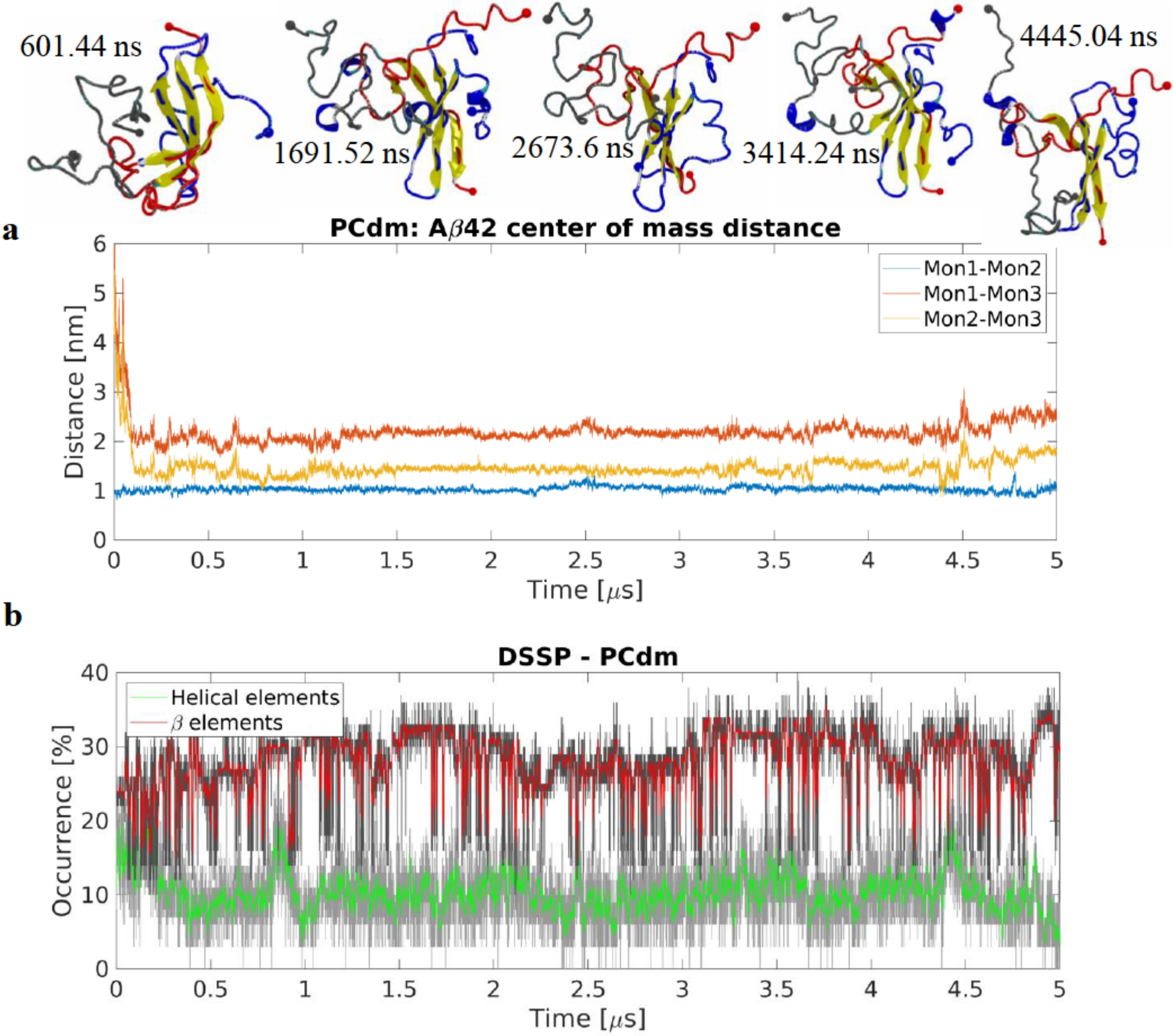


**Supplementary Figure 16. Trimer formation on POPC from membrane-bound Aβ42 dimer and free monomer (PCdm system).** (**a**) Center of mass distances between monomers showing the distances between the surface-bound dimer, Mon1 and Mon2, and the free monomer, Mon3. The trimer rapidly forms with the initially free monomer interacting with Mon2. Throughout the simulation, the initial dimer remains stable and does not experience significant conformational change. Snapshots showing change of secondary structure of the trimer show that the initially free monomer undergoes significant conformational change as it interacts with the initial dimer. The monomers within the initial dimer experience little change in secondary structure. Aβ42 molecules are in cartoon representation following VMD coloring scheme (yellow β-strands and purple α-helices), N- and C-terminal Cα are represented as a large and a small sphere respectively; blue, red, and yellow sphere and backbone colors represent Mon1, Mon2, and Mon3 respectively. (**b**) Evolution of secondary structure β- (sheet and bridge), red, and helical (α-, π–, and 3/10-helices), green, elements as determined by DSSP (v3). The graphs are moving averages using a 1 ns window; raw data is presented as dark and light grey graphs, respectively. The trimer experiences a large change in the secondary structure during the simulation; the initial ~15% helical content drops to around ~10% while the β-structure content fluctuates significant as the simulation progresses. The final β-content is ~10% higher compared to the initial.

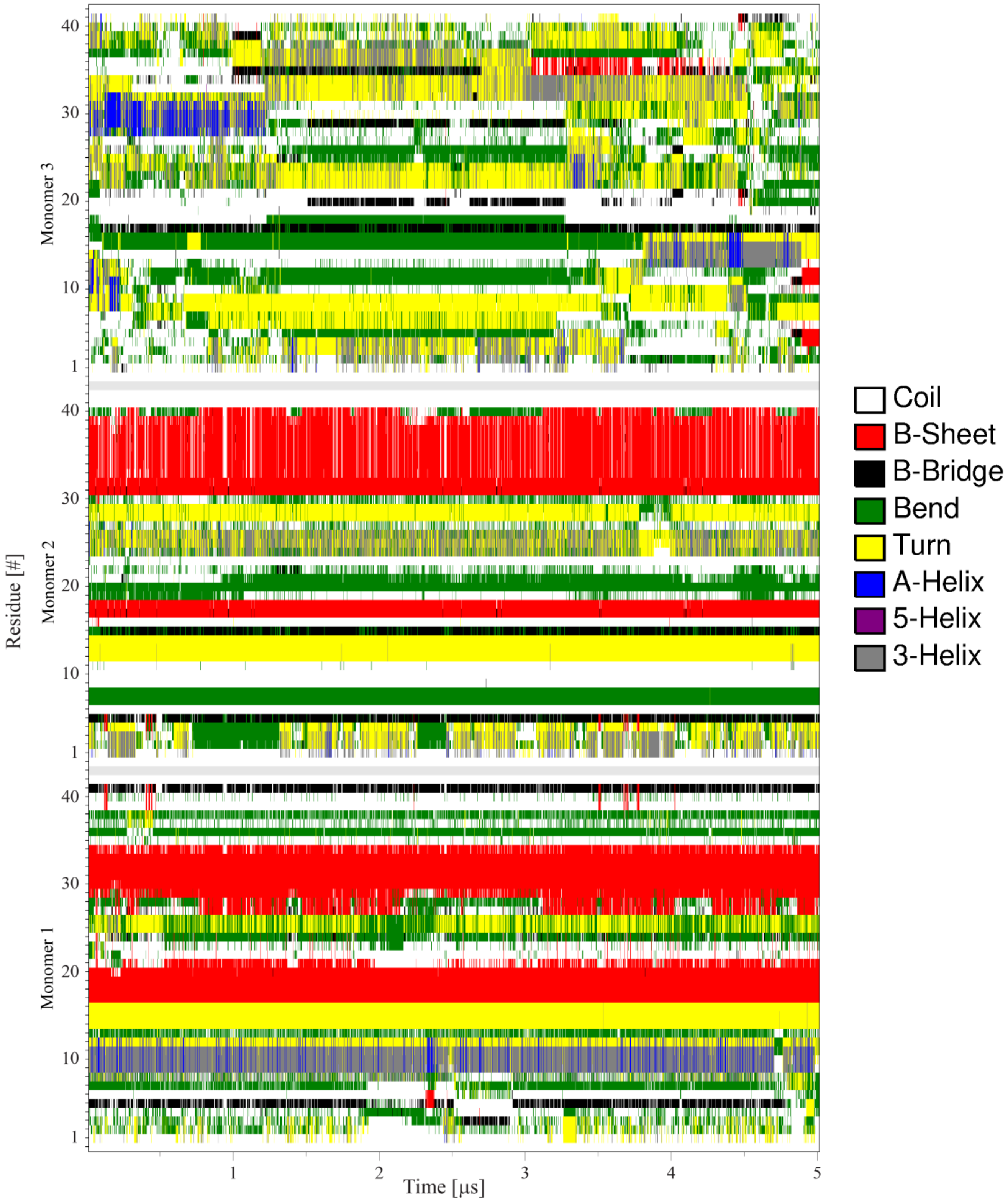


**Supplementary Figure 17. Secondary structure analysis of trimer formation on POPC from membrane-bound Aβ42 dimer and free monomer (PCdm system).** Time-resolved map of the protein secondary structure for Mon1, Mon2, and Mon3. The monomers with the initial dimer experiences fluctuations in the secondary structure, in particular in the C-terminal β-strands of Mon1 and Mon2. Similarly, the secondary structure of the N-terminal segments of the two monomers within the dimer also fluctuate throughout the simulation. Mon3, the initially free monomer, undergoes structural change in all segments, with the most dramatic changes happening in the C-terminal and the N-terminal that eventually lead to the formation of a small β-strand in the N-terminal.

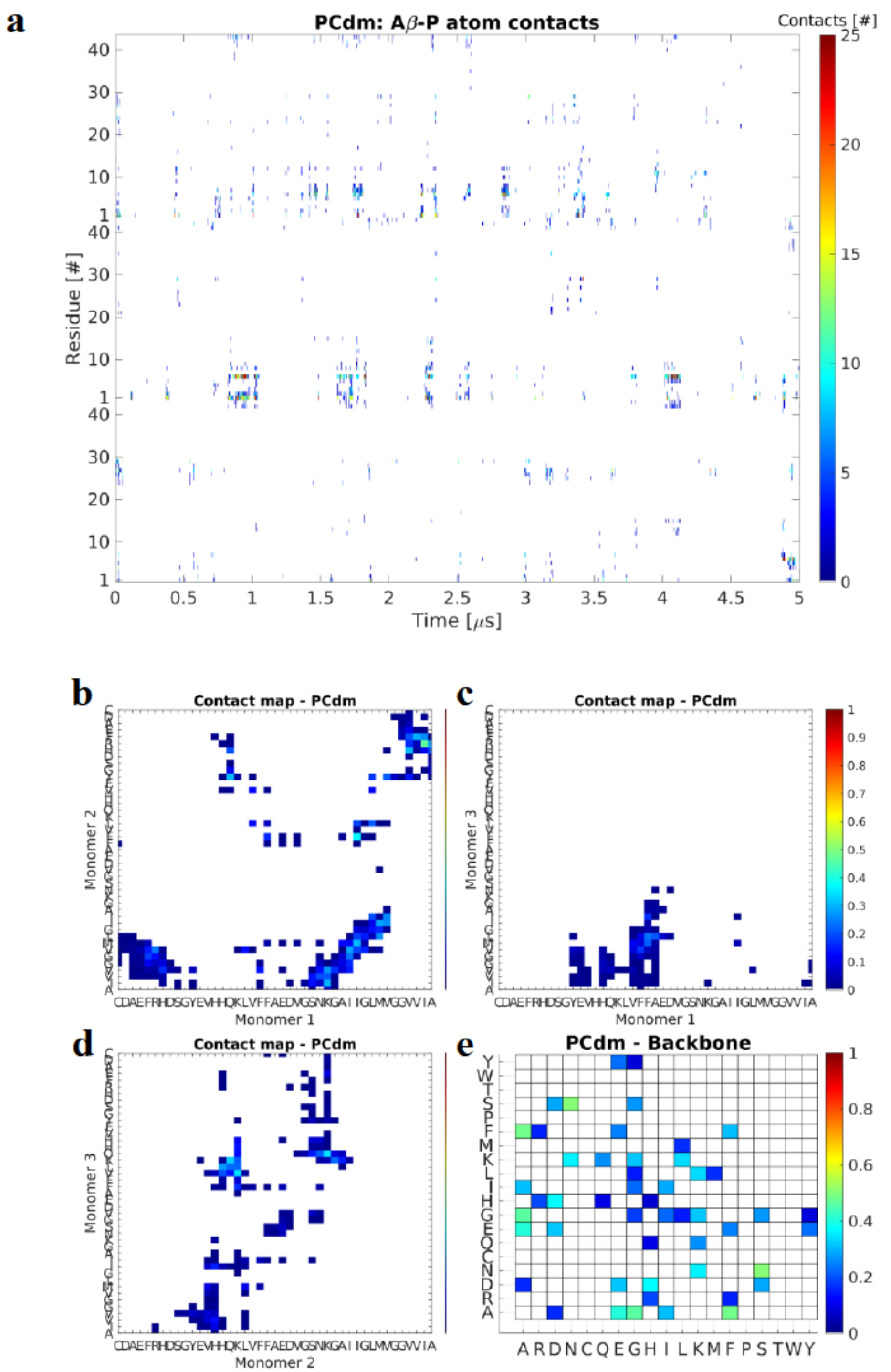


**Supplementary Figure 18. Analysis of interactions during trimer formation on POPC from membrane-bound Aβ42 dimer and free monomer (PCdm system).** (**a**) Time-resolved map of interactions between individual residues and the P atoms of the lipid headgroups. Majority of the interaction of Mon1 happens through the C-terminal, while Mon2 and Mon3 interact through the N-terminal residues. Color represent total number of contacts (<6 Å) between the residue and the lipid headgroups. (**b-d**) Normalized map, based on the residue contacts, between residue of one monomer with the opposite monomer in the PCmd system. Mon1 and Mon2 interact through N-C and C-central regions while Mon1 and Mon3 interact only through the residues in the C-central segments. Mon2-Mon3 interactions are facilitated through the central segments of the monomers with contributions from N-C interactions. Color represent total number of contacts (<6 Å) between the residue pairs. (**e**) Pivot dihedral correlation map showing the normalized residue-wise interactions important for stability of the trimer. Interactions through A, E, F, H, N, and S are very important for the stability.

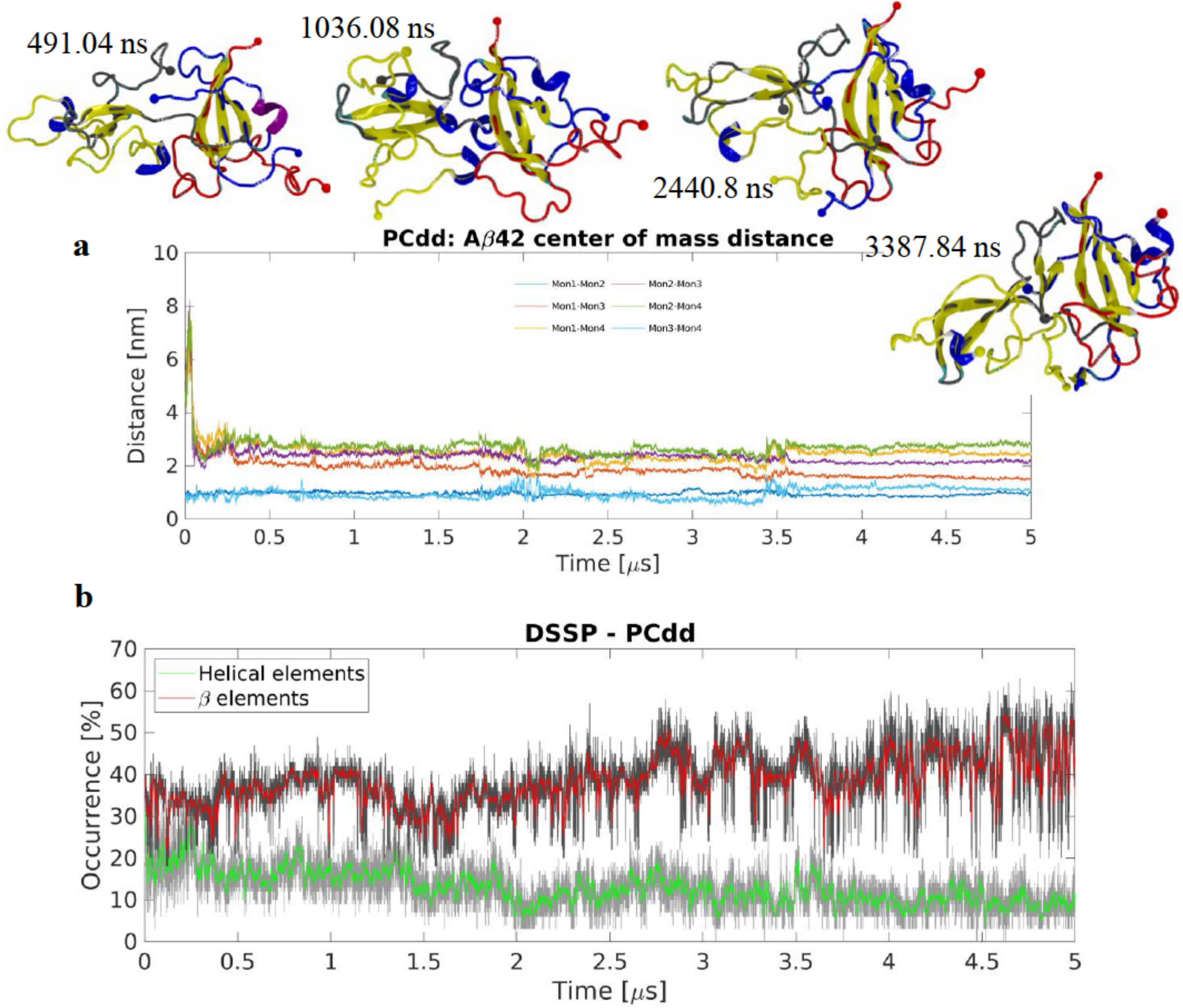


**Supplementary Figure 19. Tetramer formation on POPC from membrane-bound Aβ42 dimer and free dimer (PCdd system).** (**a**) Center of mass distances between monomers showing the distances between the surface-bound dimer, Mon1 and Mon2, and the free dimer, Mon3 and Mon4. After the formation of the tetramer, the individual monomers do not experience dramatic changes in their arrangement. Snapshots show the change in secondary structure of Aβ42 molecules as cartoon representation. The monomer largely maintain their initial secondary structures throughout the simulation, with Mon3 and Mon4 experiencing the largest amount of change as appearing and disappearing β-strands. Colors follow VMD coloring scheme (yellow β-strands and purple α-helices), N- and C-terminal Cα are represented as a large and a small sphere respectively; blue, red, yellow, and grey sphere and backbone colors represent Mon1, Mon2, Mon3, and Mon4 respectively. (**b**) Evolution of secondary structure β- (sheet and bridge), red, and helical (α-, π–, and 3/10-helices), green, elements as determined by DSSP showing that the tetramer steadily loses helical structures as the simulation progresses while experiencing an increase in β-structure content at the same time. The graphs are moving averages using a 1 ns window; raw data is presented as dark and light grey graphs, respectively.

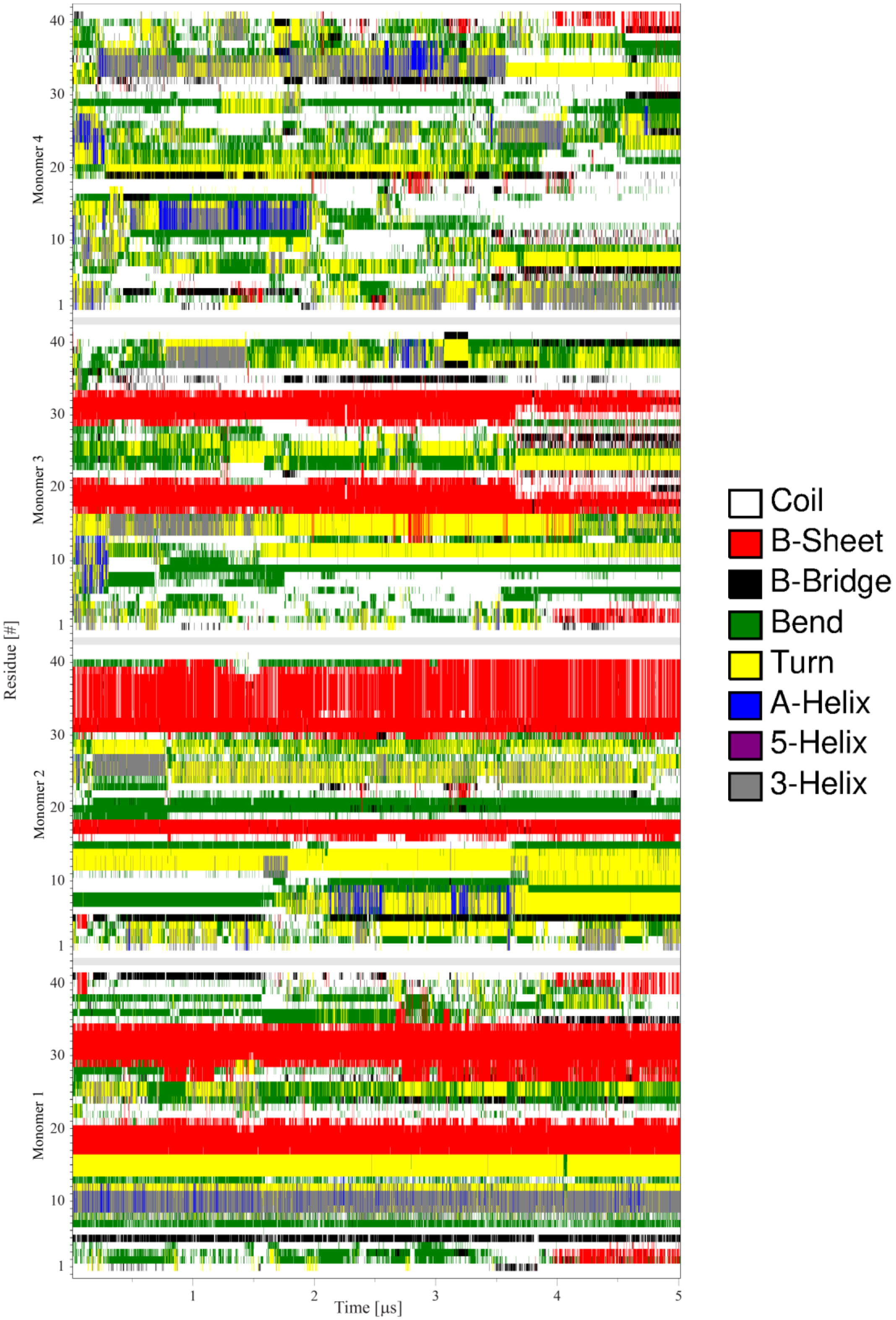


**Supplementary Figure 20. Conformational analysis of tetramer formation on POPC from membrane-bound Aβ42 dimer and free dimer (PCdd system).** Time-resolved map of the protein secondary structure for Mon1, Mon2, Mon3, and Mon4. The initially surface-bound dimer, Mon1 and Mon2, largely maintains its initial structure and only shows dynamic behavior in the N-termini of the monomers within the dimer. Whereas the initially free dimer, Mon3 and Mon4, is more dynamics, in particular Mon4 experiences significant change in secondary structure adopting small β-structure and helical segments before finally assuming a conformation with a small C-terminal β-strand.

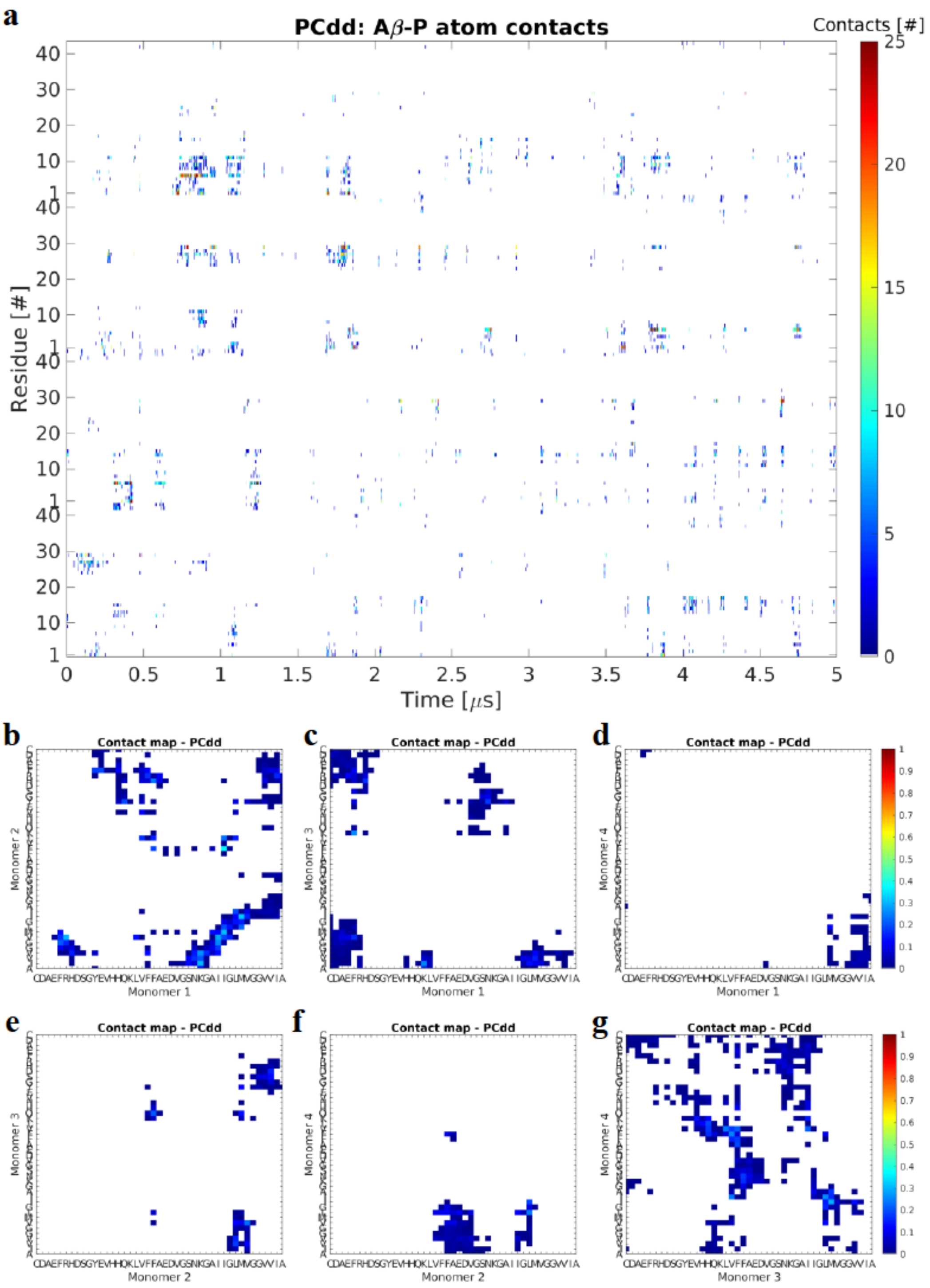


**Supplementary Figure 21. Interaction maps for tetramer formation on POPC from membrane-bound Aβ42 dimer and free dimer (PCdd system).** (**a**) Time-resolved map of interaction between individual residues and the P atom of the lipids headgroup. As seen in the previous simulations, the interactions are primarily through the N-termini of the individual monomers. Color represent total number of contacts (<6 Å) between the residue and the lipid headgroups. (**b-g**) Normalized pairwise contact maps showing the interaction between the monomers. Mon1-Mon2 interactions primarily happen through N-C and C-C terminal segments; Mon1-Mon3 interact through their C-termini and C-central region; Mon1-Mon4 interact through a small number of C-terminal residues; Mon2 interacts with Mon3 and Mon4 using residues in the central segment; and Mon3-Mon4 interactions are dominated by residues from the central regions of the monomers. Color represent total number of contacts (<6 Å) between the residue pairs.

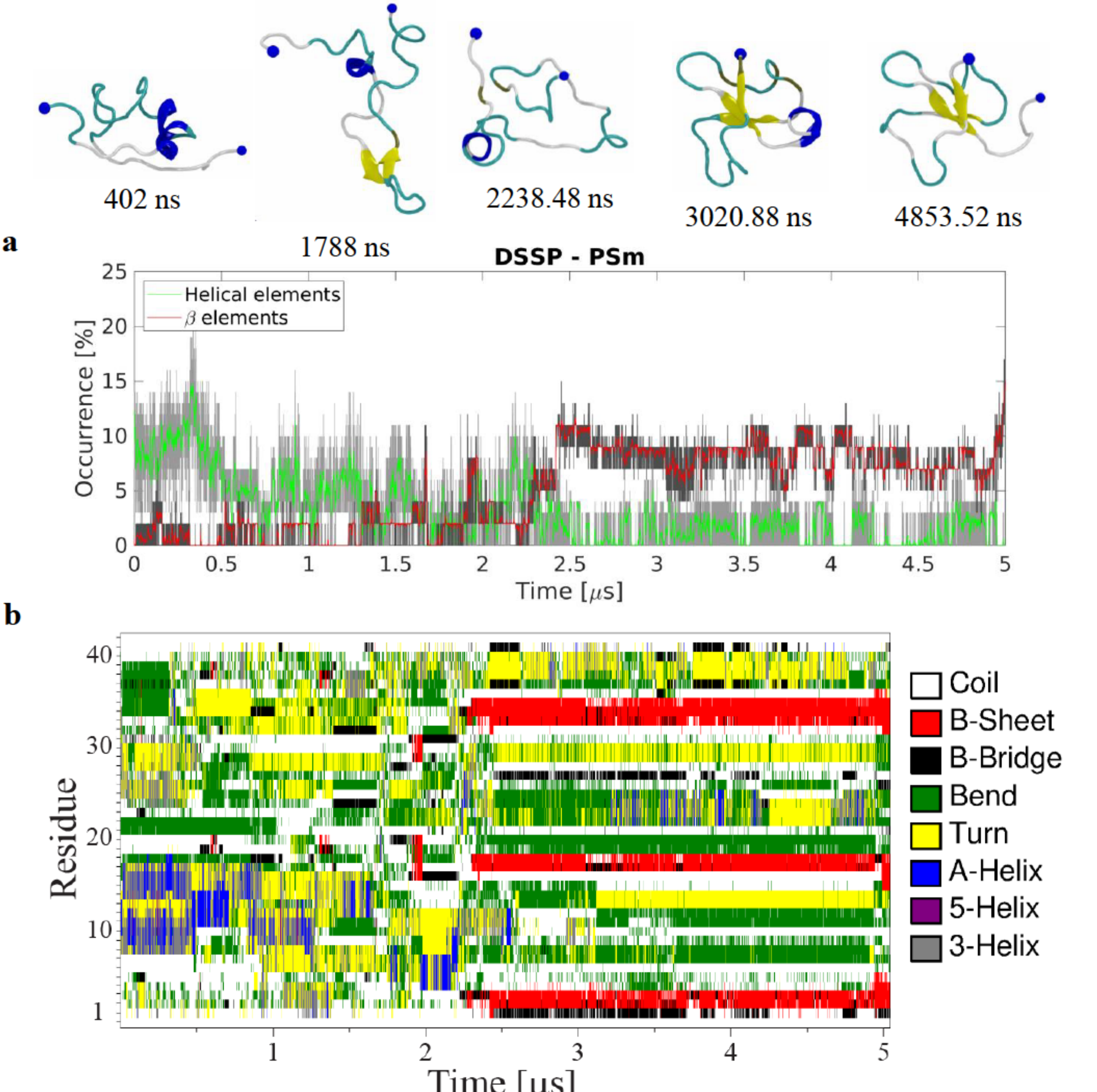


**Supplementary Figure 22. Interaction of Aβ42 monomer with POPS bilayer (PSm system). (a)** Evolution of secondary structure β- (sheet and bridge), red, and helical (α-, π–, and 3/10-helices), green, elements as determined by DSSP. After an initial increase in helical content the monomer helical content remains fluctuating around ~1-5% while the β-structure content sees a rapid growth around ~2.5 μs that plateaus for the rest of the simulation around ~7.5%. The graphs are moving averages using a 1 ns window; raw data is presented as dark and light grey graphs, respectively. Snapshots show the adoption of a transient N-terminal α-helix that is later converted into a small helical segment before the monomer finally adopts a small three-strand β-sheet. Colors follow VMD coloring scheme (yellow β-strands and purple α-helices), N- and C-terminal Cα are represented as a large and a small sphere respectively. (**b**) Time-resolved secondary structure map of the protein showing the residue-specific change over time as the monomer goes from a helical conformation to a conformation with a small β-sheet that spans the N-terminal, central, and C-terminal regions.

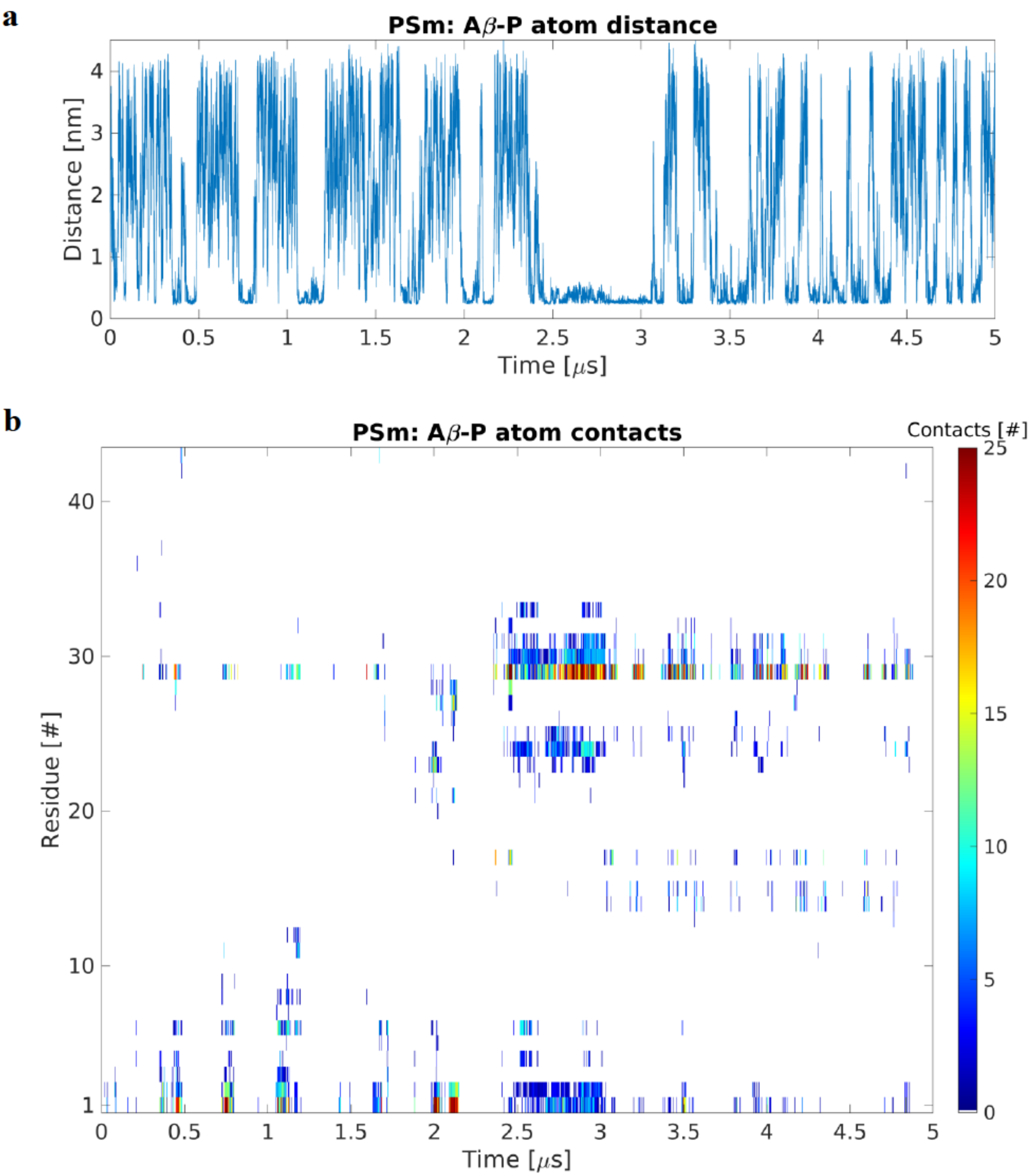


**Supplementary Figure 23. Interaction of Aβ42 monomer with POPS bilayer (PSm system). (a)** Minimum distance between the Aβ42 monomer and the P atoms of the POPS headgroups. The graph shows that while the monomer does interact with the bilayer quite often, it does not stay on the surface for longer periods of time. (**b**) Time-resolved map of interactions between individual residues and the P atoms of the lipid headgroups. As on POPC, the interaction with the membrane is facilitated by residues in the N-terminal and the 20-30 segment. Color represent total number of contacts (<6 Å) between the residue and the lipid headgroups.

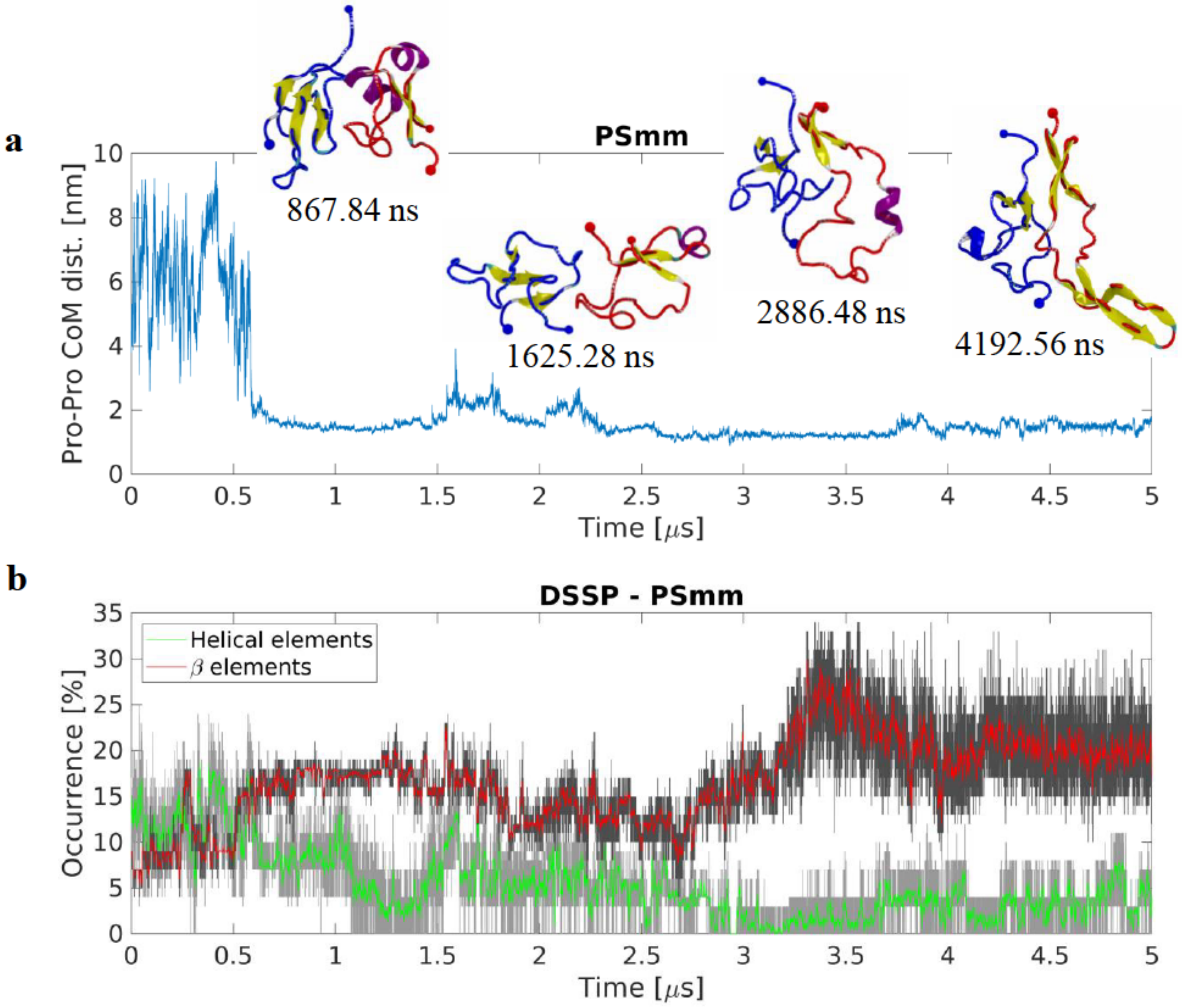


**Supplementary Figure 24. Interaction of bilayer-bound Aβ42 monomer with free monomer on POPS bilayer (PSmm system).** (**a**) Center of mass distance between monomers in the PSmm system showing the distance between the surface-bound monomer, Mon1, and the free monomer, Mon2. A stable dimer is formed after ~520 ns and remains in the same configuration for the rest of the simulation. Snapshots showing the dynamics of the interaction process and the change in the arrangement and secondary structure of the monomers within the dimer. Colors follow VMD coloring scheme (yellow β-strands and purple α-helices), N- and C-terminal Cα are represented as a large and a small sphere respectively; Mon1 and Mon2 backbone and sphere colors are blue and red, respectively. (**b**) Evolution of secondary structure β- (sheet and bridge), red, and helical (α-, π–, and 3/10-helices), green, elements as determined by DSSP. The formation of the dimer causes a dramatic increase of the β-structure content and a decrease in the helical content. This change is further enhanced around ~2.6 μs and results in a final β-structure content of ~20%. The graphs are moving averages using a 1 ns window; raw data is presented as dark and light grey graphs, respectively.

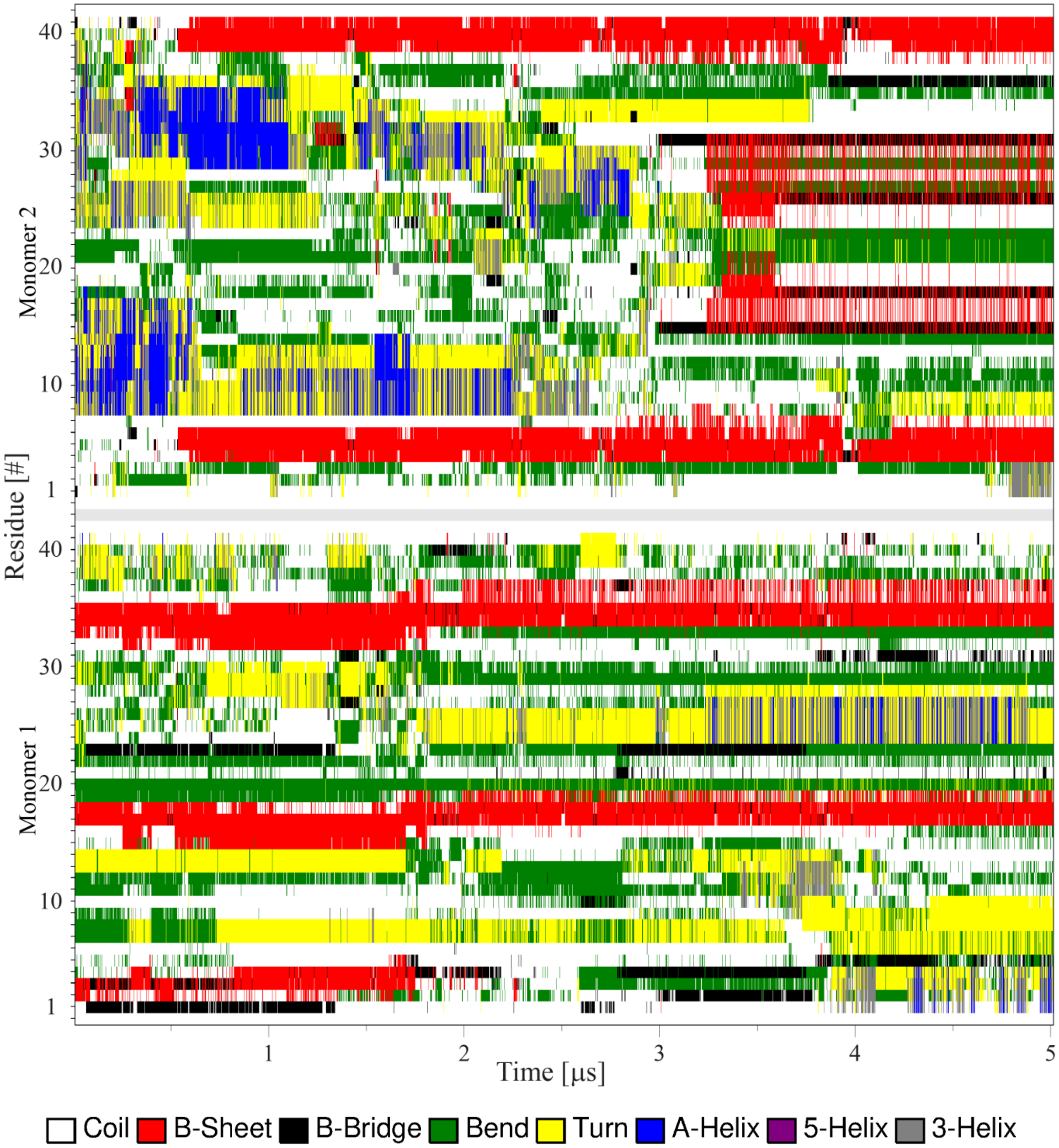


**Supplementary Figure 25. Analysis of secondary structure during interaction of bilayer-bound Aβ42 monomer with free monomer on POPS bilayer (PSmm system).** Time-resolved map of the protein secondary structure for the Aβ42 monomers. Mon1, the initially surface-bound monomer, maintains its structure with little change for the first ~1.8 µs of the simulation but then loses the N-terminal β-strand while the central and C-terminal strands become smaller. During the same time, after the dimer has formed (at ~520 ns), Mon2 undergoes a helix-to-strand conversion that starts with the loss of the N-terminal and central helix, followed by appearance of a C-terminal helix and β-strand and an N-terminal strand. The newly formed strands remain stable for the remaining simulation time, while the C-terminal helix is converted into two additional strands in the central and C-terminal regions.

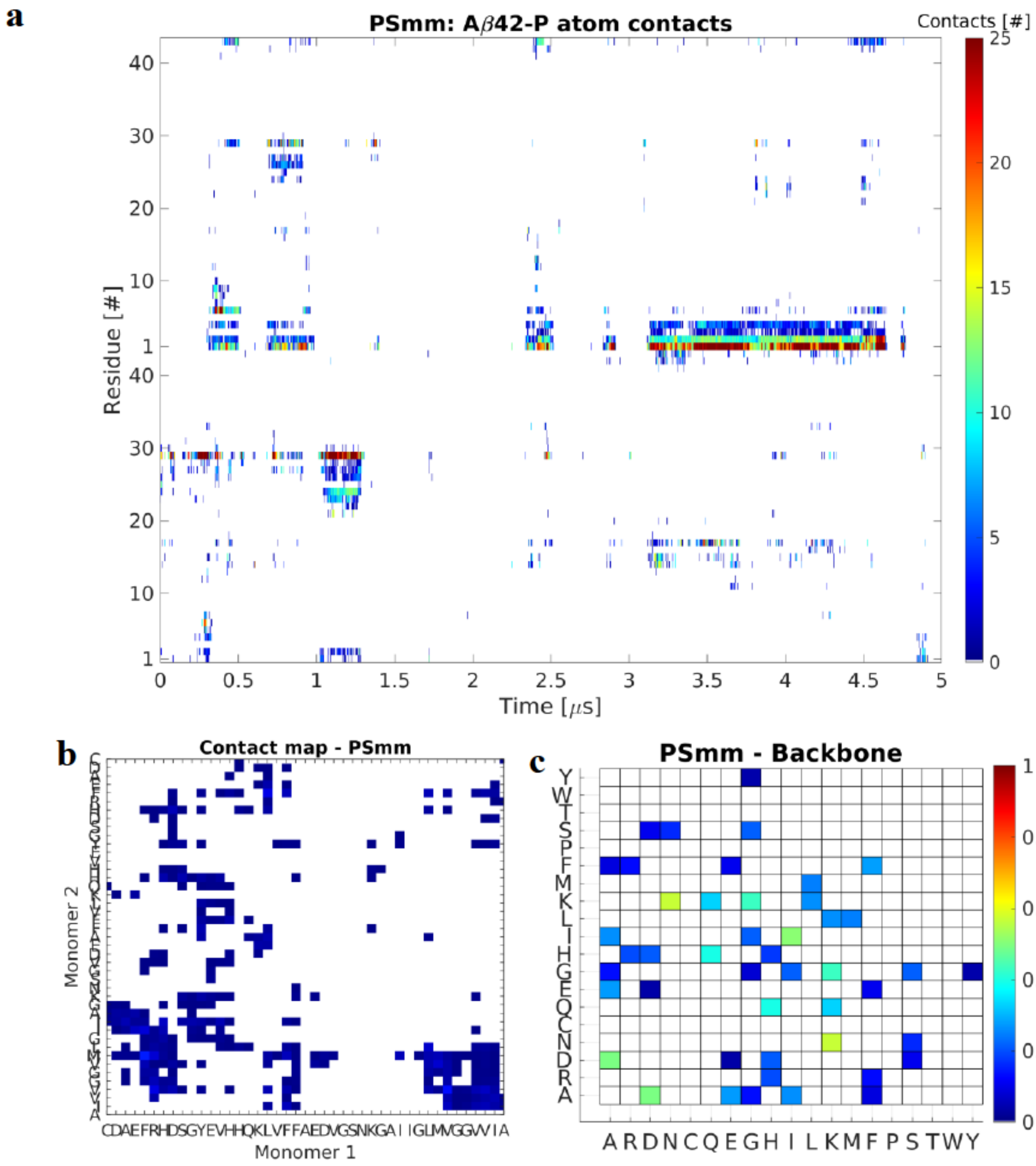


**Supplementary Figure 26. Analysis of contacts during interaction of bilayer-bound Aβ42 monomer with free monomer on POPS bilayer (PSmm system).** (**a**) Time-resolved map of interactions between individual residues and the P atoms of the lipid headgroups. As with previous results, the monomers interact with the membrane through N-terminal and residues 20-30. However, unlike before at around ~3.2 μs the dimer remains on the surface for ~1.6 μs; consequently this is the period in which helix-strand conversion in the dimer happens (Fig. S25). Color represent total number of contacts (<6 Å) between the residue and the lipid headgroups. (**b**) Normalized per-residue contact map showing that the dimer is stabilized through N-C and C-C interactions with a significant contribution from N-central region interactions as well. Colors represent total number of contacts (<6 Å) between the residue pairs. (**c**) Pivot dihedral correlation map showing the normalized residue-wise interactions important for stability of the dimer. In particular residues A, D, I, K, N, and Q are very important for the dimer.

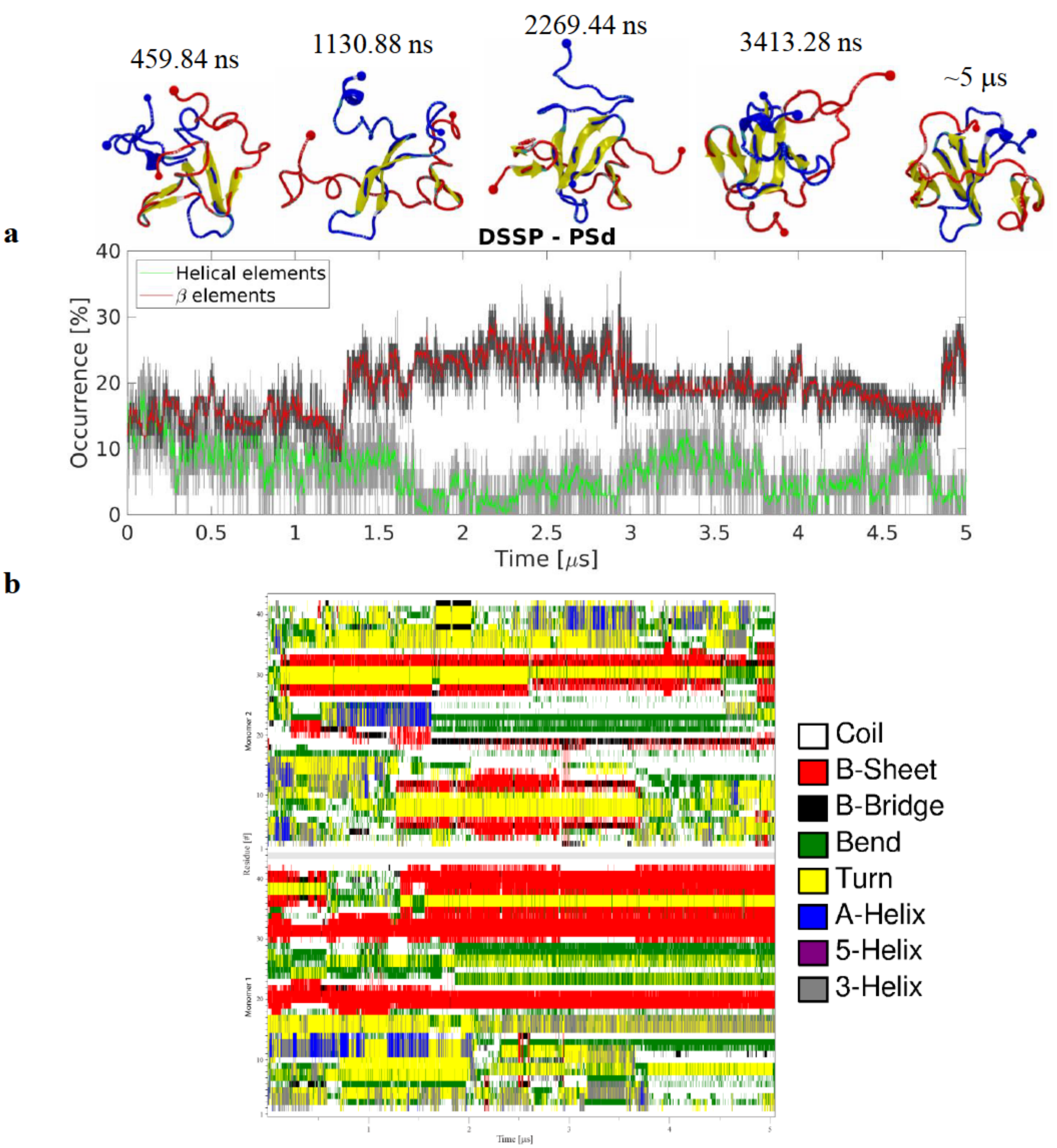


**Supplementary Figure 27. Interaction of Aβ42 dimer with POPS bilayer (PSd system). (a)** Evolution of secondary structure β- (sheet and bridge), red, and helical (α-, π–, and 3/10-helices), green, elements as determined by DSSP showing that the dimer undergoes a stepwise increase in β-content while helical content fluctuates between 1-10%. The graphs are moving averages using a 1 ns window; raw data is presented as dark and light grey graphs, respectively. Snapshots show the change in monomer arrangement and secondary structure of the Aβ42 molecules as cartoon representation following VMD coloring scheme (yellow β-strands and purple α-helices). N- and C-terminal Cα are represented as a large and a small sphere respectively; blue and red sphere and backbone colors represent Mon1 and Mon2, respectively. (**b**) DSSP plot showing the evolution of protein secondary structure as a function of time. Significant change of structure occur in Mon1 C-terminal and the Mon2 N-terminal with both experiencing increase in β-structure.

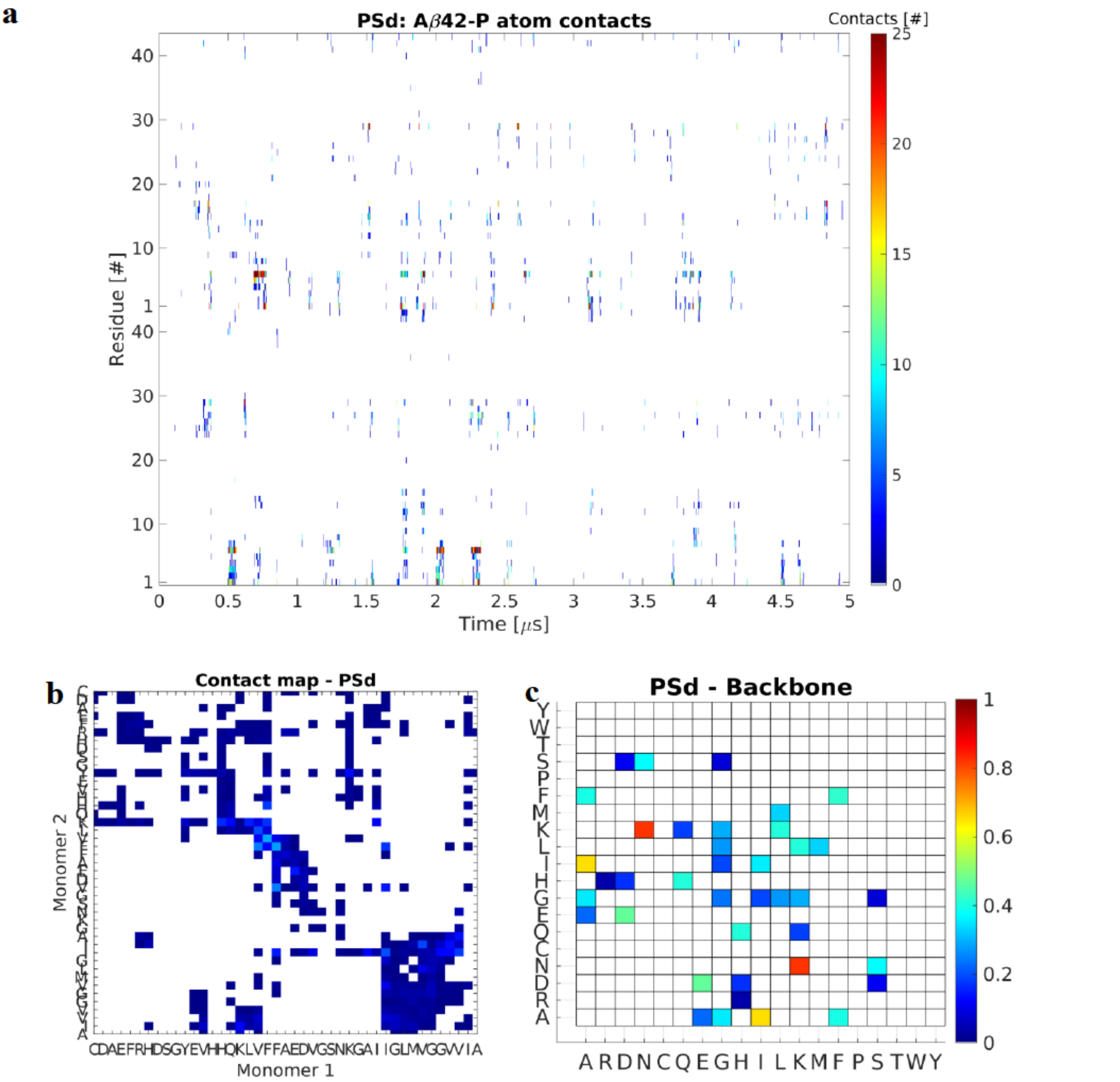


**Supplementary Figure 28. Analysis of interactions of Aβ42 dimer in presence of POPS bilayer (PSd system).** (**a**) Time-resolved map of interactions between individual residues and the P atom of the lipid headgroups. Color represent total number of contacts (<6A) between the residue and the lipid headgroups. N-terminal and residues 20-30 are responsible for the majority of the interactions with the membrane. Mon2 also has significant contribution from residue 10-20. (**b**) Normalized residue contact map, showing the interaction between the monomers within the dimer. Majority of interactions occur through the central regions of the monomers with significant contribution from C-C terminal interactions. Colors represent total number of contacts (<6 Å) between the residue pairs. (**c**) Pivot dihedral correlation map showing the normalized residue-wise interactions important for stability of the complex. Interactions involving residues A, D, E, F, I, K, L, and N are critically important for the dimer.

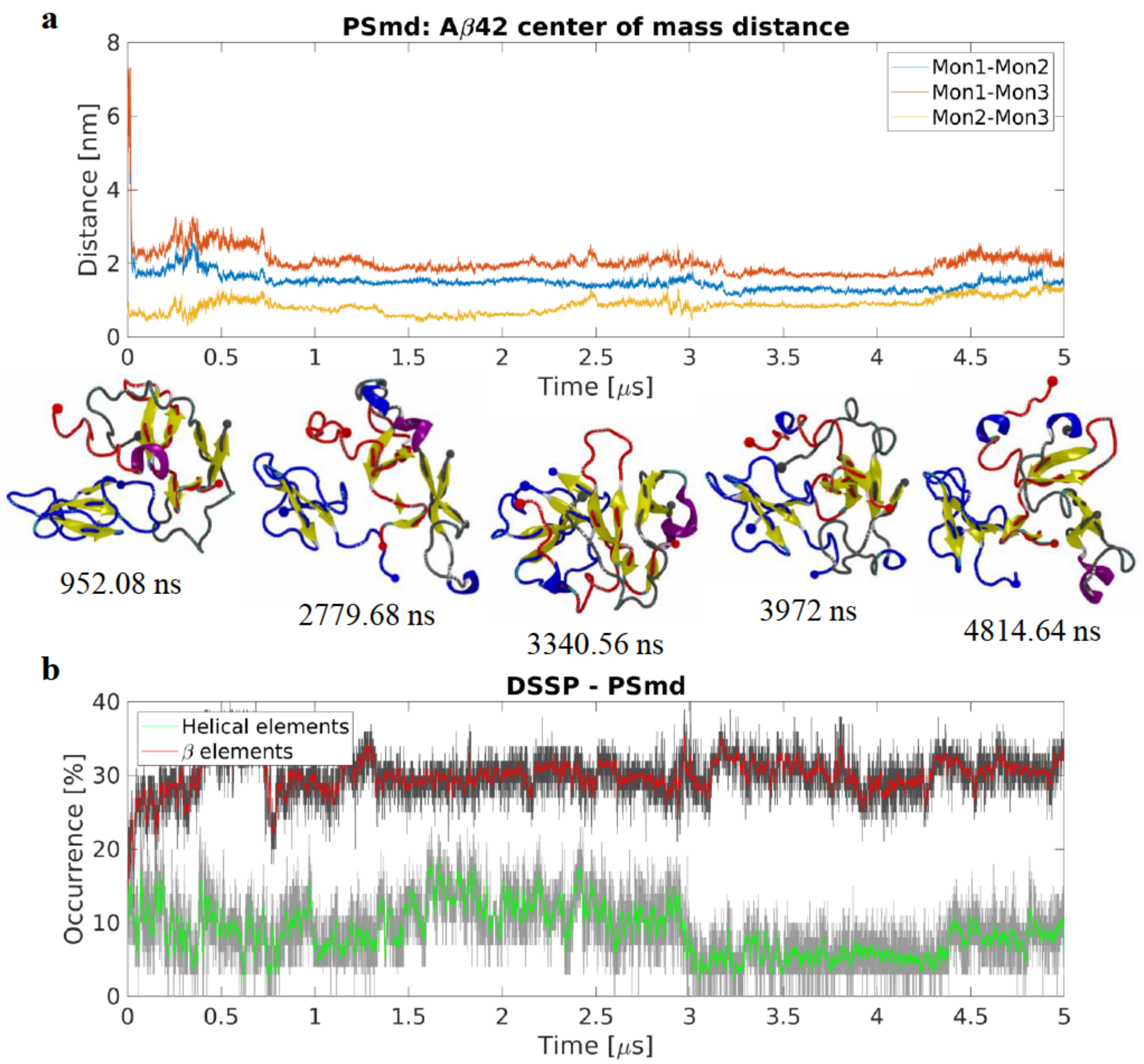


**Supplementary Figure 29. Trimer formation on POPS from membrane-bound Aβ42 monomer and free dimer in the (PSmd system).** (**a**) Center of mass distance between monomers showing the distance between the surface-bound monomer, Mon1, and the free dimer, Mon2 and Mon3. After the rapid formation of the trimer the arrangement of the monomers undergo small changes, seen as fluctuations in the CoM distances. Snapshots show arrangement and structure of the monomers within the trimer. Aβ42 molecules are in cartoon representation following VMD coloring scheme (yellow β-strands and purple α-helices), N- and C-terminal Cα are represented as a large and a small sphere respectively; blue, red, and grey sphere and backbone colors represent Mon1, Mon2, and Mon3 respectively. (**b**) Evolution of secondary β- (sheet and bridge), red, and helical (α-, π–, and 3/10-helices), green, elements as determined by DSSP. After an initial increase in β-structure content in the trimer the β-content fluctuates around ~30%. Helical content remains around ~10% for the majority of the simulation, although it also experiences fluctuations. The graphs are moving averages using a 1 ns window; raw data is presented as dark and light grey graphs, respectively.

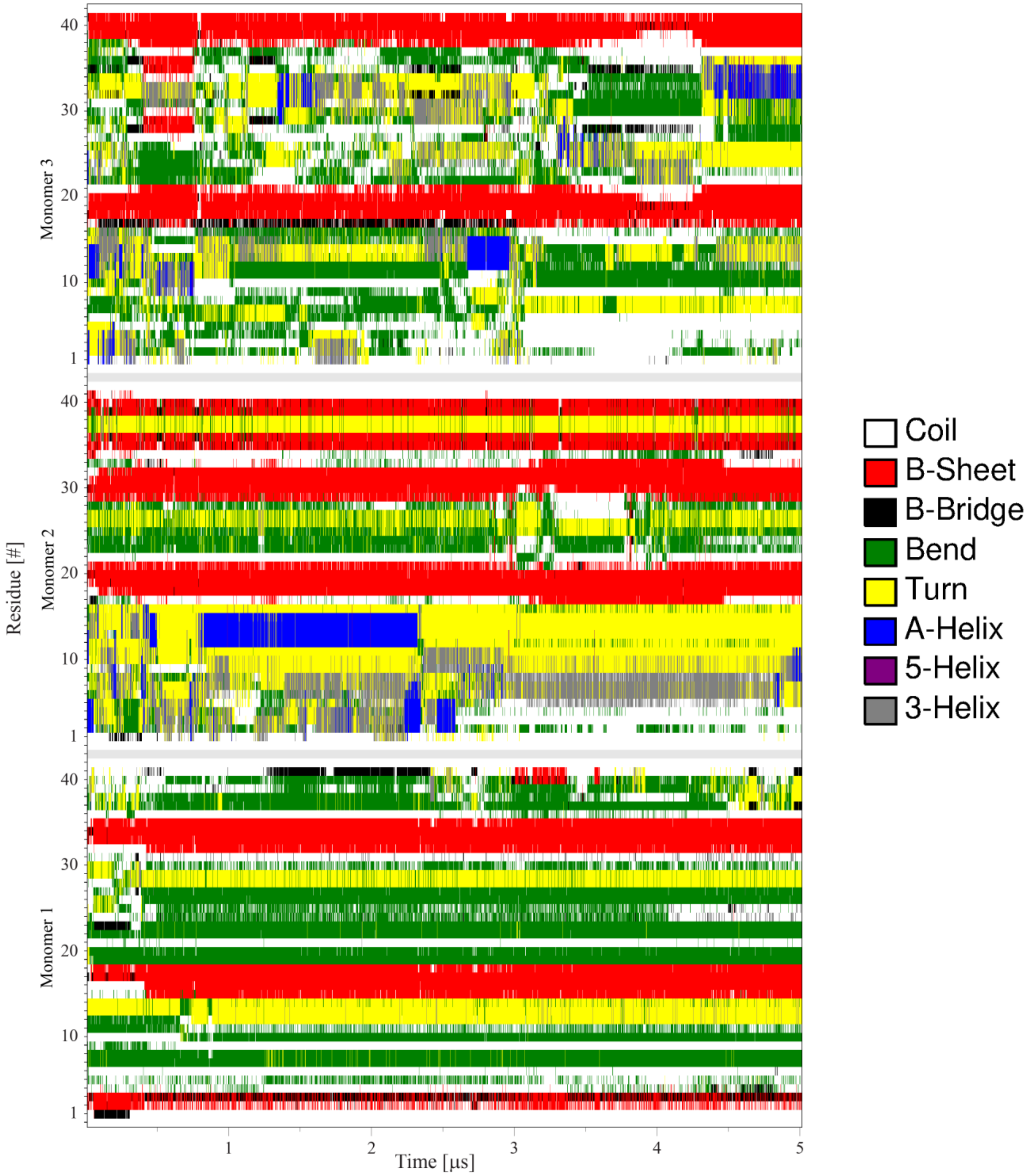


**Supplementary Figure 30. Conformational analysis of interaction of bilayer-bound Aβ42 monomer with free dimer on POPS bilayer (PSmd system).** DSSP plot showing the evolution of protein secondary structure as a function of time. The initially membrane-bound monomer, Mon1, does not experience a significant change in structure as it maintains the small β-strands in the termini and central region. Mon2 maintain three small β-strands in the central and C-terminal regions while it experiences fluctuations in the secondary structure of the N-terminal region. Mon3 is the most dynamic monomer in the system and experiences significant changes in all regions, however it also has short β-strands in the C-terminal and central (17-21) regions.

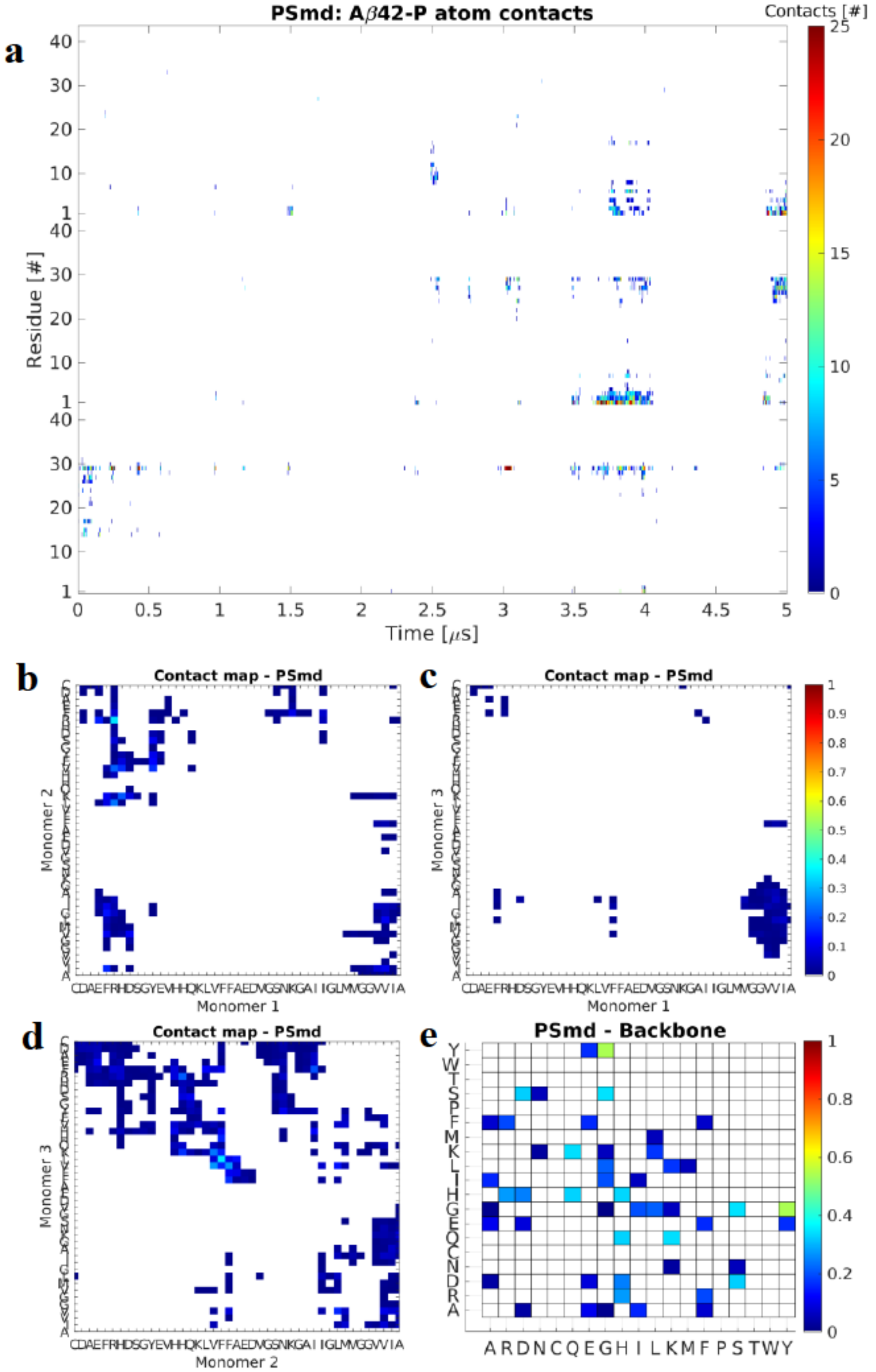


**Supplementary Figure 31. Analysis of interactions of bilayer-bound Aβ42 monomer with free dimer on POPS bilayer (PSmd system).** (**a**) Time-resolved map of interactions between individual residues and the P atoms of the lipid headgroups. Mon1 interacts with the membrane exclusively through residues 12-30; Mon2 interacts through N-terminal and residues 20-30; and Mon3, interacts through residues 1-20. Colors represent total number of contacts (<6 Å) between the residue and the lipid headgroups. (**b-d**) Normalized residue contact maps showing the regions important for interactions between the monomers. Mon1-Mon2 interact through their N- and C- termini; Mon1-Mon3 interact almost exclusively through C-C terminal interactions; while Mon2-Mon3 interact through the central regions as well as the N- and C-termini. Color represent total number of contacts (<6 Å) between the residue pairs. (**e**) Pivot dihedral correlation map showing the normalized residue-wise interactions important for stability of the trimer. Interactions involving residue H, K, Q, S, and Y are identified as important for the trimer.

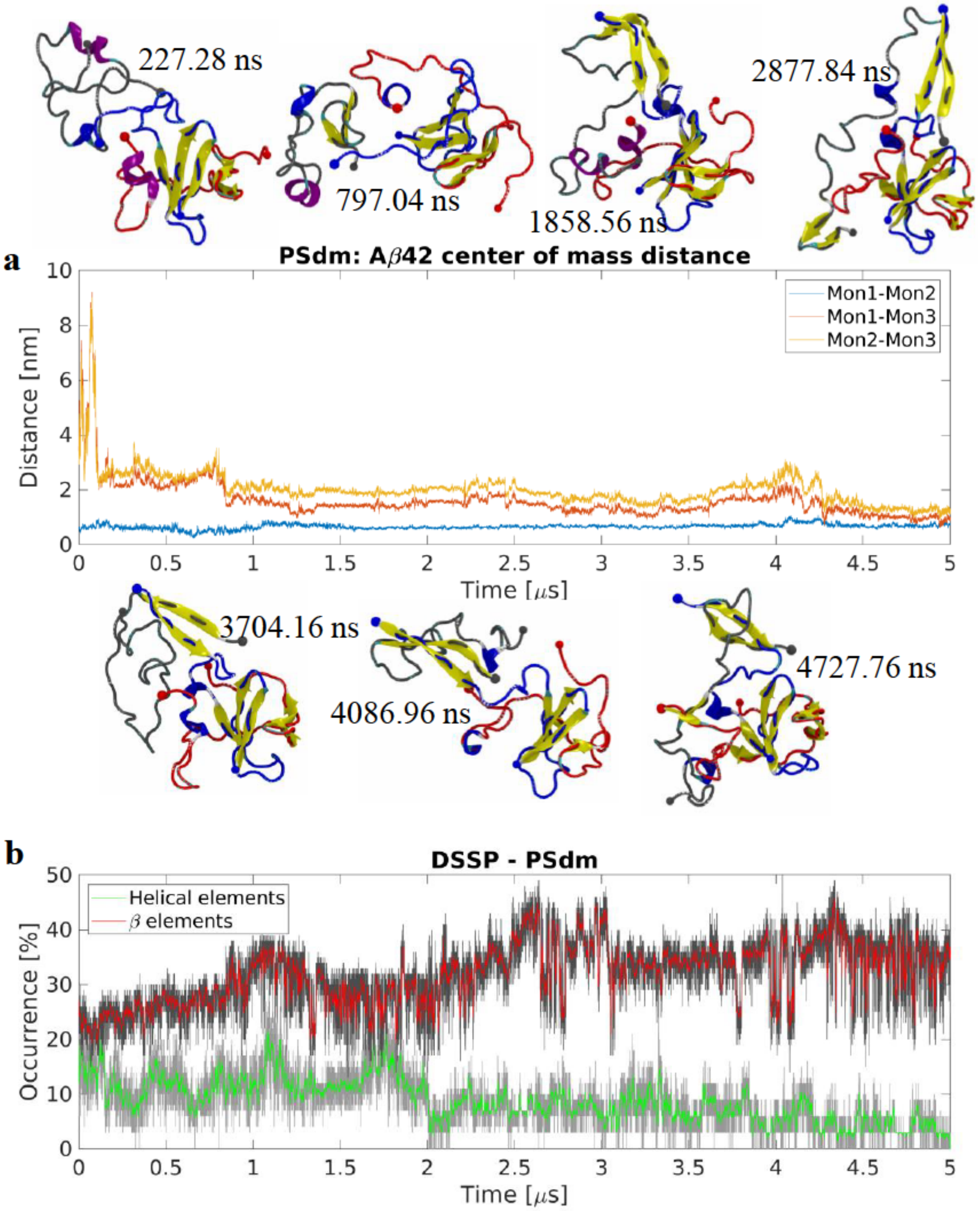


**Supplementary Figure 32. Interaction of bilayer-bound Aβ42 dimer with free monomer on POPS bilayer (PSdm system).** (**a**) Center of mass distance between the surface-bound dimer, Mon1 and Mon2, and the free monomer, Mon3, showing the rapid formation of the trimer. Snapshots show the change of secondary structure as well as the re-arrangement of the monomers within the trimer as the simulation progresses. The initial dimer remains as a compact unit while Mon3 remains connected in an extended configuration. Aβ42 molecules are in cartoon representation following VMD coloring scheme (yellow β-strands and purple α-helices), N- and C-terminal Cα are represented as a large and a small sphere respectively; blue, red, and grey sphere and backbone colors represent Mon1, Mon2, and Mon3 respectively. (**b**) Evolution of secondary structure β- (sheet and bridge), red, and helical (α-, π–, and 3/10-helices), green, elements as determined by DSSP showing the steady increase in β-content while helical content slowly decreases. The graphs are moving averages using a 1 ns window; raw data is presented as dark and light grey graphs, respectively.

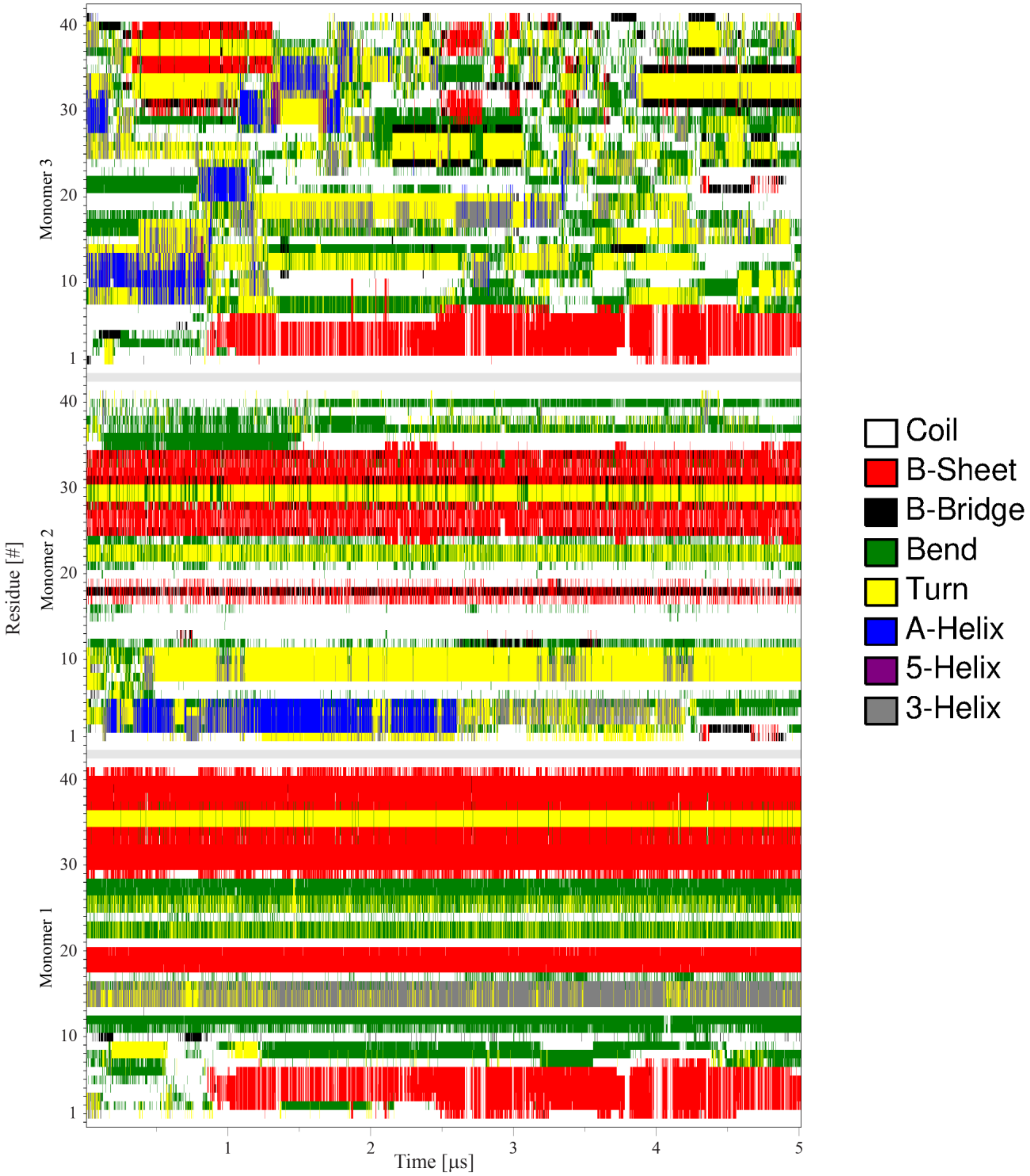


**Supplementary Figure 33. Analysis of secondary structure during interaction of bilayer-bound Aβ42 dimer with free monomer on POPS bilayer (PSdm system)**. DSSP plot showing the evolution of protein secondary structure as a function of time. Mon1 structure is very stable and, apart from the appearance a β-strand in the N-terminal region, does not dramatically change from its initial conformation. Mon2 also has the same tendencies, and the central and C-terminal regions remain largely unchanged during the simulation while the N-terminal forms an α-helix before returning back to its original conformation. Mon3, the initially free monomer, is the most dynamic in terms of structural change as it undergoes several dramatic transitions: within the first 2 μs, the C-terminal transitions from an α-helix to a short β-strand and back to a helix, the central region gains an α-helix that is quickly lost again, and the N-terminal α-helix is converted into a short β-strand that remains for the rest of the simulation. The central and C-terminal regions experience continuous structural change for the rest of the simulation before finally adopting a short C-terminal β-strand.

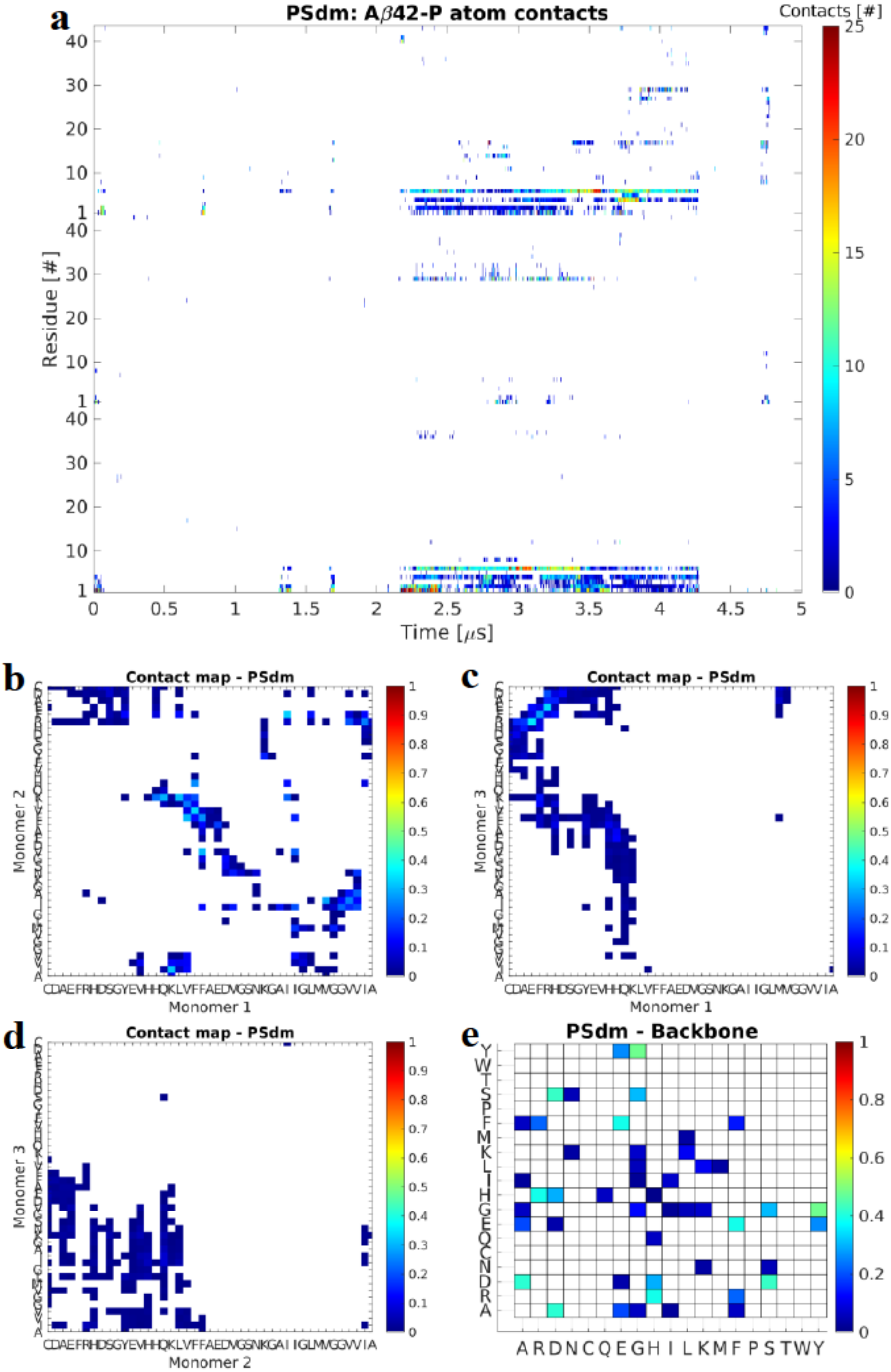


**Supplementary Figure 34. Analysis of contacts during interaction of bilayer-bound Aβ42 dimer with free monomer on POPS bilayer (PSdm system)**. (**a**) Time-resolved map of interactions between individual residues and the P atoms of the lipid headgroups. During the first ~2.2 μs the trimer sporadically interacts with the membrane through the N-termini of the monomers following which the trimer stays bound to the membrane surface for the next ~2 μs through the N-termini of Mon1 and Mon3 and to a lesser extent the C-terminal residues of Mon2. Color represent total number of contacts (<6 Å) between the residue and the lipid headgroups. (**b-d**) Normalized residue contact maps showing that Mon1-Mon2 interactions are facilitated by the central regions of the monomer as well as the C-termini. Mon1-Mon3 interaction happened through N-C interactions and N-central regions, while Mon2-Mon3 interactions occur primarily through the N-C termini of the monomers. Colors represent total number of contacts (<6 Å) between the residue pairs. (**e**) Pivot dihedral correlation map showing the normalized residue-wise interactions important for stability of the trimer. In particular residues A, D, F, H, S, and Y are important contributor to the trimer stability.

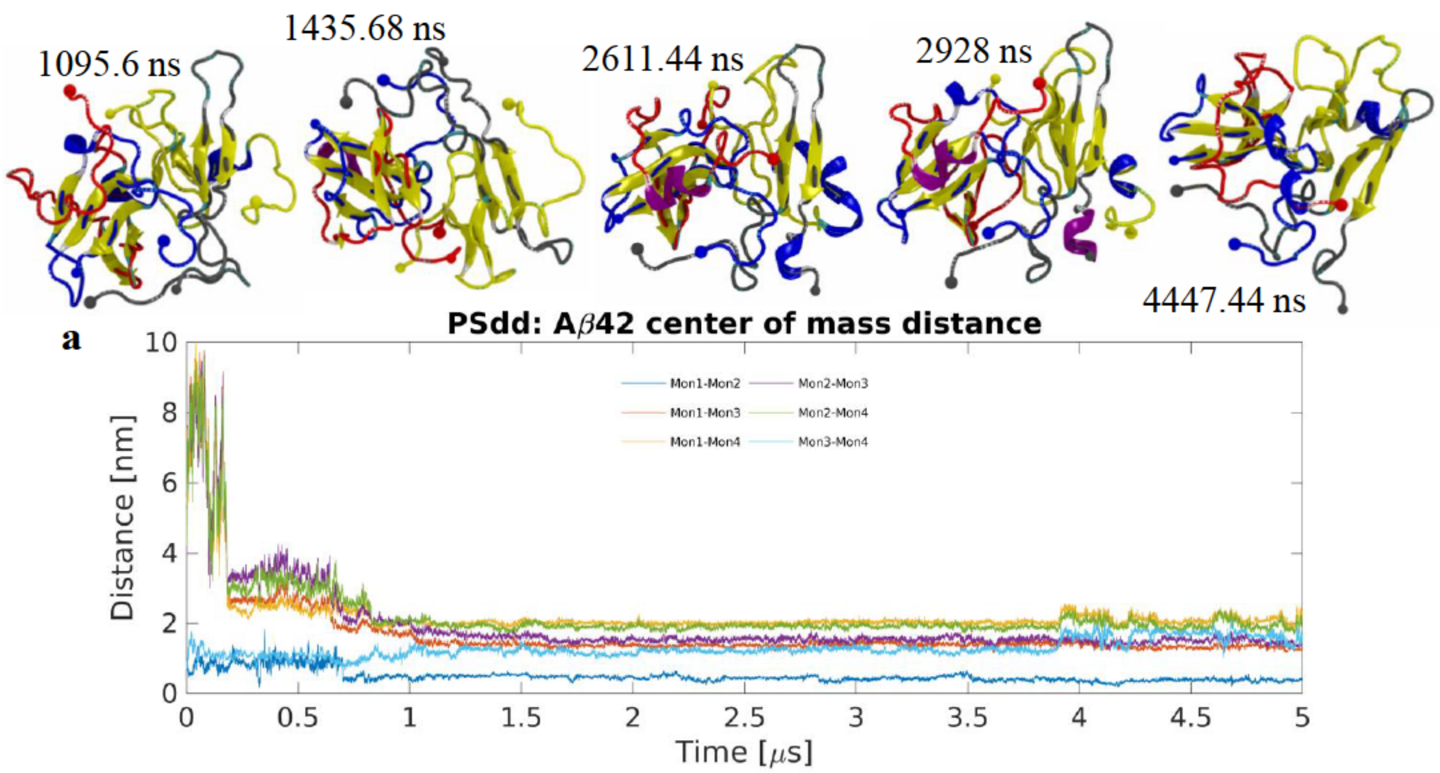


**Supplementary Figure 35. Tetramer formation on POPS from membrane-bound Aβ42 dimer and free dimer (PSdd system).** (**a**) Center of mass distance between monomers showing the distance between the surface-bound dimer, Mon1 and Mon2, and the free dimer, Mon3 and Mon4. After the tetramer has formed the monomers undergo a re-arrangement that causes the CoM to decrease and remain around ~2 nm for the rest of the simulation. Snapshots show that while the monomers within the tetramer undergo small change in the secondary structure during the simulation and that the initial dimers remain as compact subunits of the tetramer. Aβ42 molecules are in cartoon representation following VMD coloring scheme (yellow β-strands and purple α-helices), N- and C-terminal Cα are represented as a large and a small sphere respectively; blue, red, grey, and yellow sphere and backbone colors represent Mon1, Mon2, Mon3, and Mon4 respectively. (**b**) Evolution of secondary structure β- (sheet and bridge), red, and helical (α-, π–, and 3/10-helices), green, elements as determined by DSSP showing that the β-content increases a small amount while the helical content slowly decreases as the simulation progresses. The graphs are moving averages using a 1 ns window; raw data is presented as dark and light grey graphs, respectively.

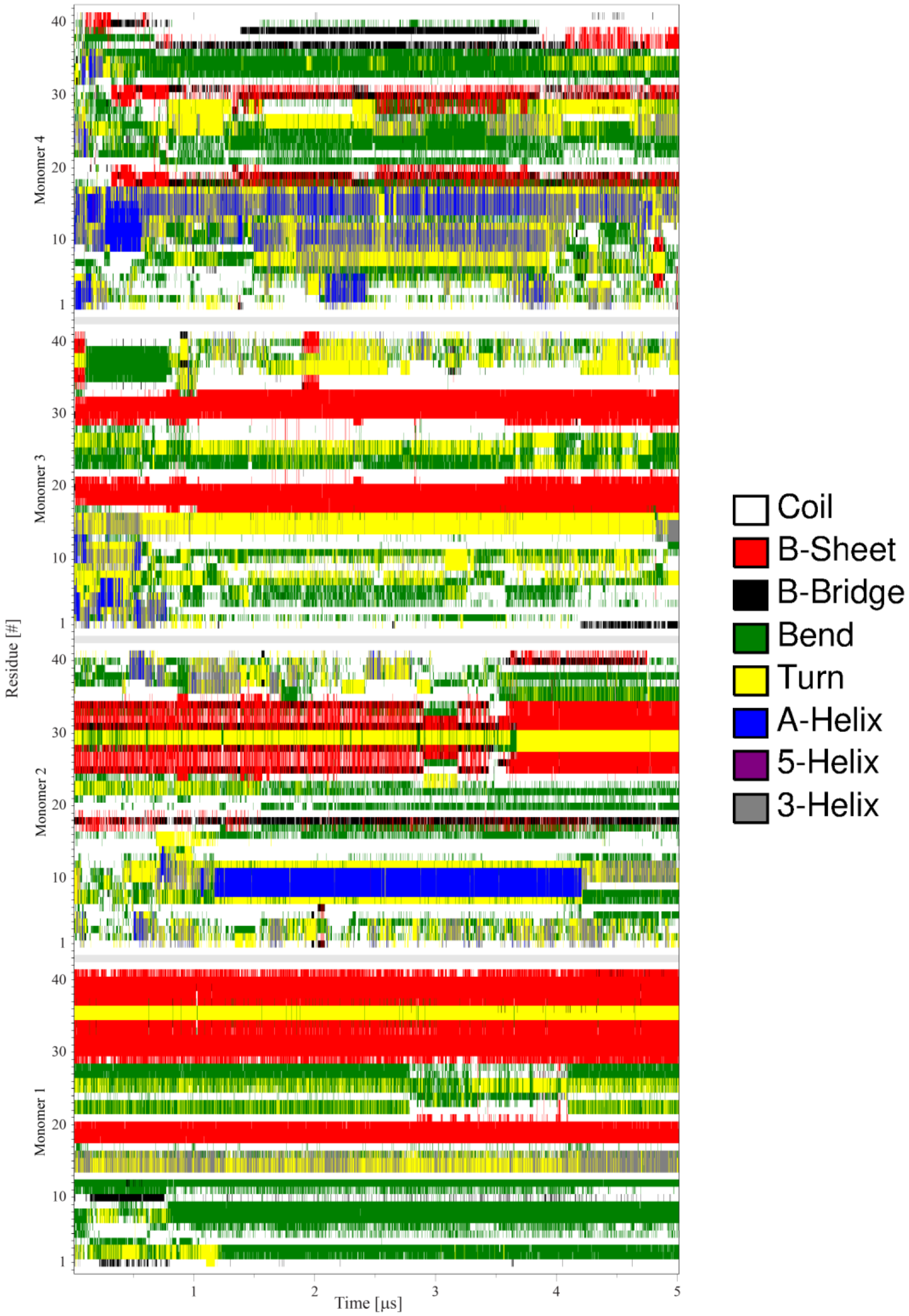


**Supplementary Figure 36. Conformational analysis of tetramer formation on POPS from membrane-bound Aβ42 dimer and free dimer (PSdd system).** DSSP plot showing the evolution of protein secondary structure as a function of time. Mon1 and Mon3 show no significant change in their structures during the simulation; Mon2 on the other hand is dynamic, in regards to secondary structure, and its N-terminus adopts helical structure that eventually goes back to a less structured configuration. The C-terminal also experiences fluctuation in structure but to a much smaller degree and largely remains as β-structure. Mon4 has a very dynamic N-terminus and central region that rapidly transition to and from small segments of α-helices, while the C-terminus remains largely unstructured before adopting a small β-strand at the end of the simulation.

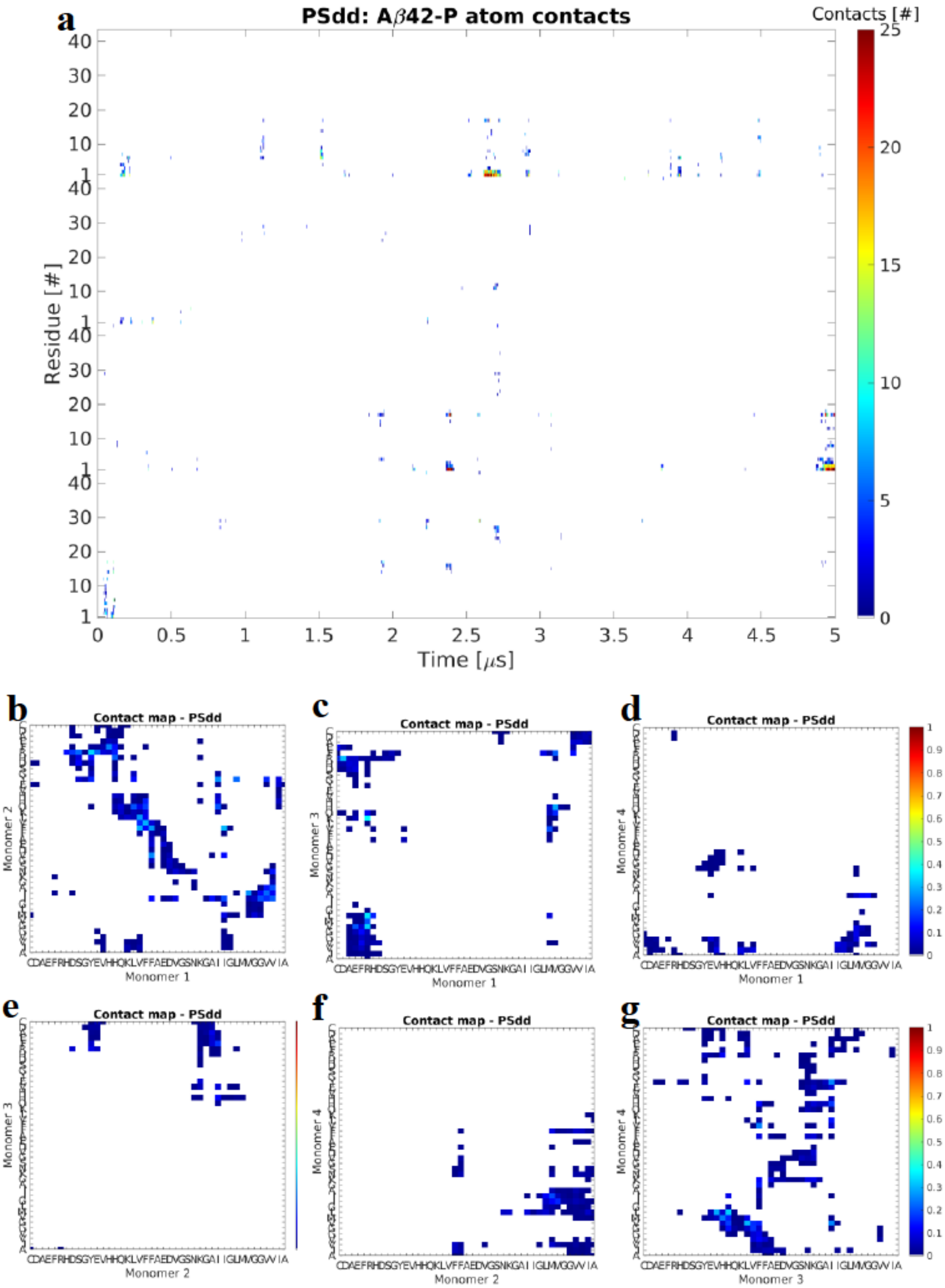


**Supplementary Figure 37. Analysis of interactions during tetramer formation on POPS from membrane-bound Aβ42 dimer and free dimer (PSdd system).** (**a**) Time-resolved map of interaction between individual residues and the P atom of the lipids headgroup. As seen in the previous simulations, the interactions are primarily through the N-termini of the individual monomers, in particular Mon2 and Mon4. Colors represent total number of contacts (<6 Å) between the residue and the lipid headgroups. (**b-g**) Normalized pairwise contact maps showing the interaction between the monomers. Mon1-Mon2 interactions primarily happen through C-C termini and the central segments; Mon1-Mon3 interact through N-C- and C-C termini; Mon1-Mon4 interact through a small number of residues in the N- and C-termini and C-C termini; Mon2 interacts with Mon3 through residues in the N-terminus and central segment and with Mon4 through C-C terminal interactions. Mon3-Mon4 interactions are dominated by residues from the central regions of the monomers and C-central interactions. Colors represent total number of contacts (<6 Å) between the residue pairs.
